# A cellular and transcriptomic dissection of the human breast for studying mechanisms of cell and tissue function

**DOI:** 10.1101/2022.10.25.513729

**Authors:** Katelyn Del Toro, Rosalyn W. Sayaman, Kate Thi, Yamhilette Licon-Munoz, William C. Hines

## Abstract

A fundamental question in biology, central to our understanding of cancer and other pathologies, is determining how different cell types coordinate to form and maintain tissues. Recognizing the distinct features and capabilities of the cells that compose these tissues is critical. Unfortunately, the complexity of tissues often hinders our ability to distinguish between neighboring cell types and, in turn, scrutinize their transcriptomes and generate reliable and tractable cell models for studying their inherently different biologies. In a companion article, we introduced a novel method that permits the identification and purification of the twelve cell types that compose the human breast—nearly all of which could be reliably propagated in the laboratory. Here, we explore the nature of these cell types. We sequence mRNAs from each purified population and investigate transcriptional patterns that reveal their distinguishing features. We describe the differentially expressed genes and enriched biological pathways that capture the essence of each cell type, and we highlight transcripts that display intriguing expression patterns. These data, analytic tools, and transcriptional analyses form a rich resource whose exploration provides remarkable insights into the inner workings of the cell types composing the breast, thus furthering our understanding of the rules governing normal cell and tissue function.

## Introduction

Understanding the biology of tissues and ultimately developing effective cancer therapies requires us to recognize the symphony of interactions in tissues that occur among its many different cell types—and the myriad of molecules they synthesize. Determining how cells communicate with each other and their microenvironments is critical, as is defining the roles of stromal cells in regulating epithelial cell behavior, shaping tissues, and promoting or preventing overt cancer^1–4^. To this puzzle, there are many pieces. Clarifying and appreciating the roles of each cell type requires a substantial foundation in our ability to identify and distinguish among these distinct tissue elements so we may learn how they fit together to form a functioning organ. Tumors are caricatures of normal tissues^5^, yet the intricate cellular composition and arrangement of cells in many tissues, including the breast, have yet to be adequately dissected, leaving many unanswered questions about these cells’ unique and essential qualities.

The human dissections and inspiring observations made by Andreas Vesalius, William Harvey, and other Renaissance anatomist-physicians produced astonishing breakthroughs in medical science^6, 7^. In a companion article and with a similar passion for anatomical dissection, we applied modern tools of histology and cell and molecular biology to examine the fabric of the human breast—to learn its cellular composition and study its mechanistic underpinnings. Using immunofluorescence and advanced flow cytometry, we developed methods to resolve, isolate, quantify, and culture each breast cell type, reconciling every cell in the tissue. The product of this labor was a fluorescence-activated cell sorting (FACS) strategy that used redundant markers and a rigorous gating strategy to identify the twelve distinct breast cell types (Figure 1a, Figure 1—figure supplement 1)]. These populations included two separate luminal epithelial cell types, myoepithelial cells, adipocytes, leukocytes, pericytes, vascular smooth muscle cells, erythrocytes, adipose-derived mesenchymal stem cells (fibroblasts), lymphatic and vascular endothelial cells, and a newly identified rare epithelial cell population with unknown function.

**Figure 1.**
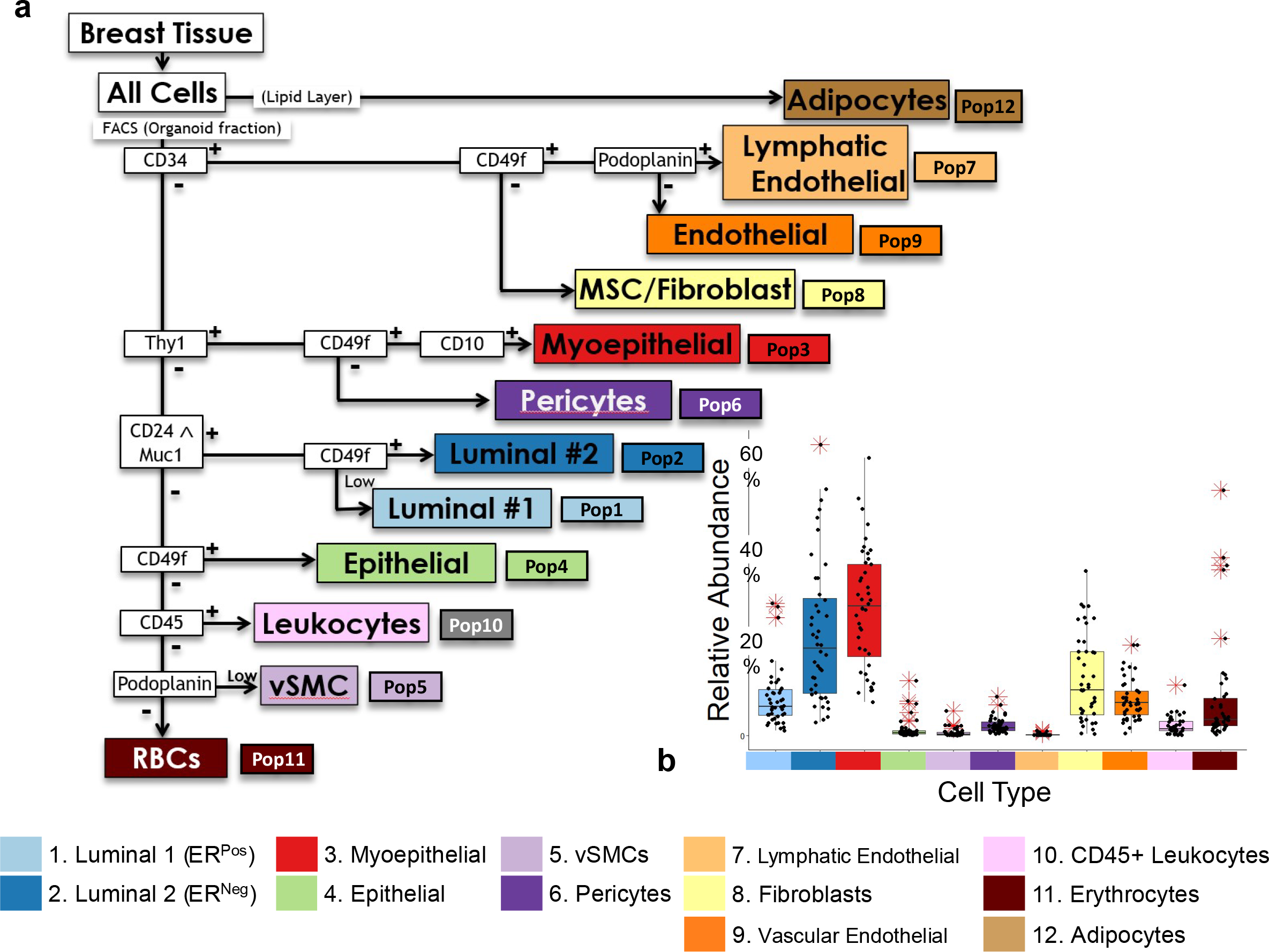
FACS gating strategy for comprehensive cell type isolation. (a) Prior to FACS isolation, breast tissues were enzymatically digested with collagenase. This was followed by low-speed centrifugation, which yielded an organoid pellet, supernatant, and an adipocyte-enriched lipid layer. RNA was isolated from the lipid layer (adipocytes, Pop12) and stored for later sequencing. The remaining cell types were FACS purified from the trypsin-dissociated organoid cell pellet, according to the outlined gating scheme. Each cell type was assigned a population number that corresponds to each population’s isolation strategy (Pops 1-12). **(b)** Relative abundance of each cell type from 30 different individuals (proportion of total cell counts, as measured by FACS analysis of 44 independent FACS experiments). Outliers (*) are defined as points falling (1.5 x IQR) above/below the 75^th^ and 25^th^ quartile.

Although this FACS method resolved leukocytes as a single CD45^Pos^ fraction, the strategy is adaptable and can be further refined to scrutinize the array of leukocyte populations in the tissue. We now routinely sort and culture nine of the twelve breast cell types (excluded are red blood cells, leukocytes, and vascular smooth muscle cells). Some of these primary models are the first of their kind from this tissue, including breast pericytes, lymphatic endothelial cells, and our newly defined epithelial cell population. Their isolation and culture add to the repertoire of available models and permit functional studies that were not previously possible^8^.

Here, we asked the critical question of what defines each breast cell type—and sought to identify their characteristic features and functions. Unsurprisingly, hierarchical clustering of transcriptional profiles echoed FACS results by cleanly separating the eleven sequenced cell types—revealing remarkable correlations among biological replicates (of each cell type, isolated from random individuals). Principal component analysis (PCA) established genes that heavily contribute to cellular diversity. We used PCA-derived gene lists to identify the distinguishing features of epithelial, mesenchymal, endothelial, and leukocyte cell lineages—each being a group of cell types with similar origins and characteristics. These included well-recognized features, such as keratinization of epithelial cells, muscle contraction by mesenchymal cells, vasculogenesis by endothelial cells, and more. While these results validated our analytic approach, they revealed many other processes we had not previously associated with a specific cell lineage. Shifting to a more gene-centric approach, we investigated what families of genes contributed to cell-type-specific functions by identifying the gene families that varied most across the spectrum of cell types. Many gene families showed little to no change in expression (e.g., exportins), whereas others were extensively differentially expressed (e.g., laminins, integrins, Rho GTPase activating proteins, and more). Differences in these genes’ expression, and the biological processes in which they are involved, were further solidified—and others identified—via comparisons between cell-type pairs and application of gene set enrichment and pathway analyses to reveal the ‘essence’ of each cell type. We highlighted six comparisons that we believed to be of the highest interest as they reveal features of every major cell type and identify those distinguishing closely related cell types. Moreover, we have performed all 55 possible pairwise cell-type comparisons and provided the results in a user-friendly analysis tool (Pathway Analyzer^9^) and web application (Breast Cell Atlas: Transcript Finder^10^). These tools allow one to thoroughly engage with the dataset and actively explore each cell type’s expressed genes and enriched biological pathways.

## Results and Discussion

### Identification of cell lineages and analytic pipeline

The breast is a compound tubuloalveolar gland whose subcutaneous tissue comprises, at a minimum, 12 major cell types. In an adjoining article, we describe the development and validation of an antibody panel and gating strategy capable of isolating these cell types^8^, cell populations we numbered from one to twelve (which refers to their respective FACS gate using the isolation strategy outlined in Figure 1a and Figure 1—figure supplement 1). These cell populations are described as follows: ER^Pos^ (Pop1/Lum1) and ER^Neg^ (Pop2/Lum2) luminal epithelial cells, myoepithelial cells (Pop3), a rare epithelial cell population (Pop4), vascular, smooth muscle cells (vSMCs, Pop5), pericytes (Pop6), lymphatic endothelial cells (Pop7), fibroblasts (Pop8), vascular endothelial cells (Pop9), CD45^Pos^ leukocytes (Pop10), erythrocytes (Pop11), and adipocytes (Pop12). These different cell types are often broadly classified as members of epithelial, endothelial, mesenchymal, or hematopoietic lineages (Figure 2b).

**Figure 2.**
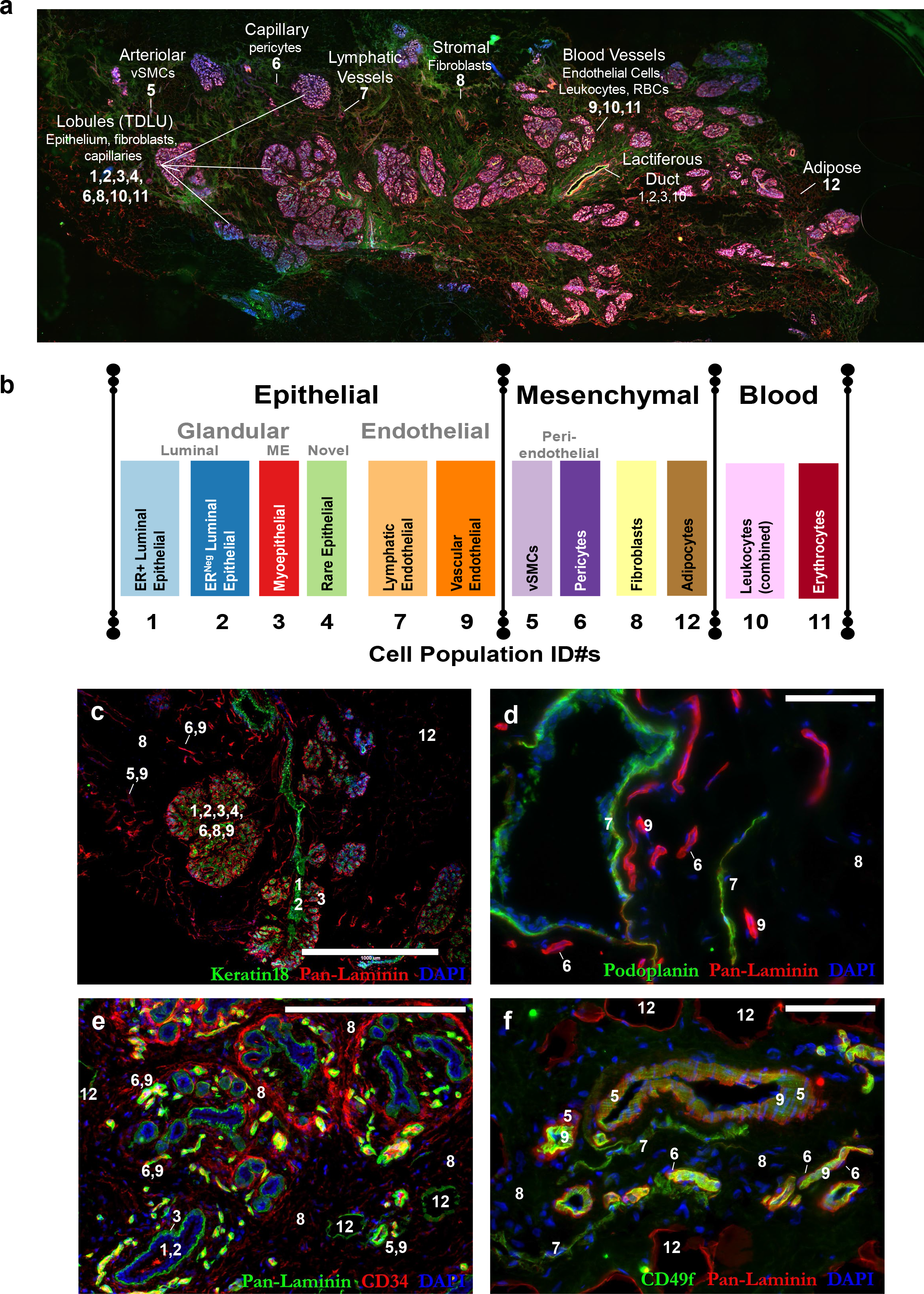
IHC staining of cell types in the breast. (a) Representative 4x scan of normal breast tissue (reduction mammoplasty tissue from 23-year-old female). The general location of the 12 cell types are indicated by the their FACS population identification numbers, Pops1-12. **(b)** Chart of the 12 breast cell types and their assigned colors (used throughout this article). Cell types are grouped by their common histological classification: epithelial, mesenchymal, and blood cell types. Epithelial types are further subdivided into glandular and endothelial types, whereas the mesenchymal perivascular types are indicated (pericytes and vascular smooth muscle cells). **(c-f)** spatial position of the cell types in normal tissue immunostained with: **(c)** keratin 18 (green; luminal epithelial cells) and pan-laminin (red; epithelial and vascular basement membrane, and adipocytes). Staining reveals a collecting duct within several breast lobules (center) composed of epithelial (Pops 1-4), fibroblasts (Pop8), and capillaries that consist of pericytes (Pop6) and vascular endothelial cells (Pop9). Stromal adipocytes are also present (Pop12, upper-right). Scale = 1000μm. **(d)** podoplanin (green) and pan-laminin (red) staining reveal podoplanin-staining lymphatic vessels (Pop7) and vascular blood vessels (Pop9). Laminin embedded pericytes (Pop6) are also discernible on the capillaries; Scale = 200μm; **(e)** CD34 (red) and pan-laminin (green) staining reveals a cross-section of three-and a-half breast lobules within a bed of CD34^Pos^ fibroblasts. Endothelial cells forming capillaries and larger blood vessels also stain for CD34, both inside and outside the lobular areas, e.g., larger vessels are visible at lower right, along with pan-laminin staining adipocytes (Pop12); Scale = 200μm **(f)** CD49f (green) and pan-laminin (red) reveals a large blood vessel (center) surrounded by vascular smooth muscle cells (Pop5). Endothelial cells (Pop9) and pericytes (Pop6) of smaller capillaries are present, as are lymphatic vessels (Pop7) and adipocytes (Pop12); Scale = 200μm.

In the early days of this study, we had not yet completed identifying all populations revealed by FACS, and several remained a mystery even after finalizing our isolation strategy. Establishing their identities was particularly challenging. We were aided by several helpful clues, however. One was the distinct morphologies the cell types displayed when cultured. These observations allowed us to determine whether a particular population appeared epithelial-like, e.g., Pop4 epithelial cells, or presented with more mesenchymal-like characteristics, e.g., Pop6 pericytes. We also backtracked and stained tissues using the same antibody combinations used in FACS and hunted for cell types in the tissue that shared identical staining patterns. Ultimately, the transcriptome information from RNA-sequencing provided definitive evidence that solidified each cell type’s identity.

With their identities and histological locations now solved (Figure 2a-f, Figure 2—figure supplement 1-2)^11^, we shifted our attention to exploring the individual biologies of these different cell types by diligently dissecting their transcriptomes through a comparative lens provided by the comprehensive collection of cell types we had isolated and sequenced. Our comprehensive analytic pipeline is described in Figure 3 (Figure 3, Figure 3—figure supplement 1).

**Figure 3.**
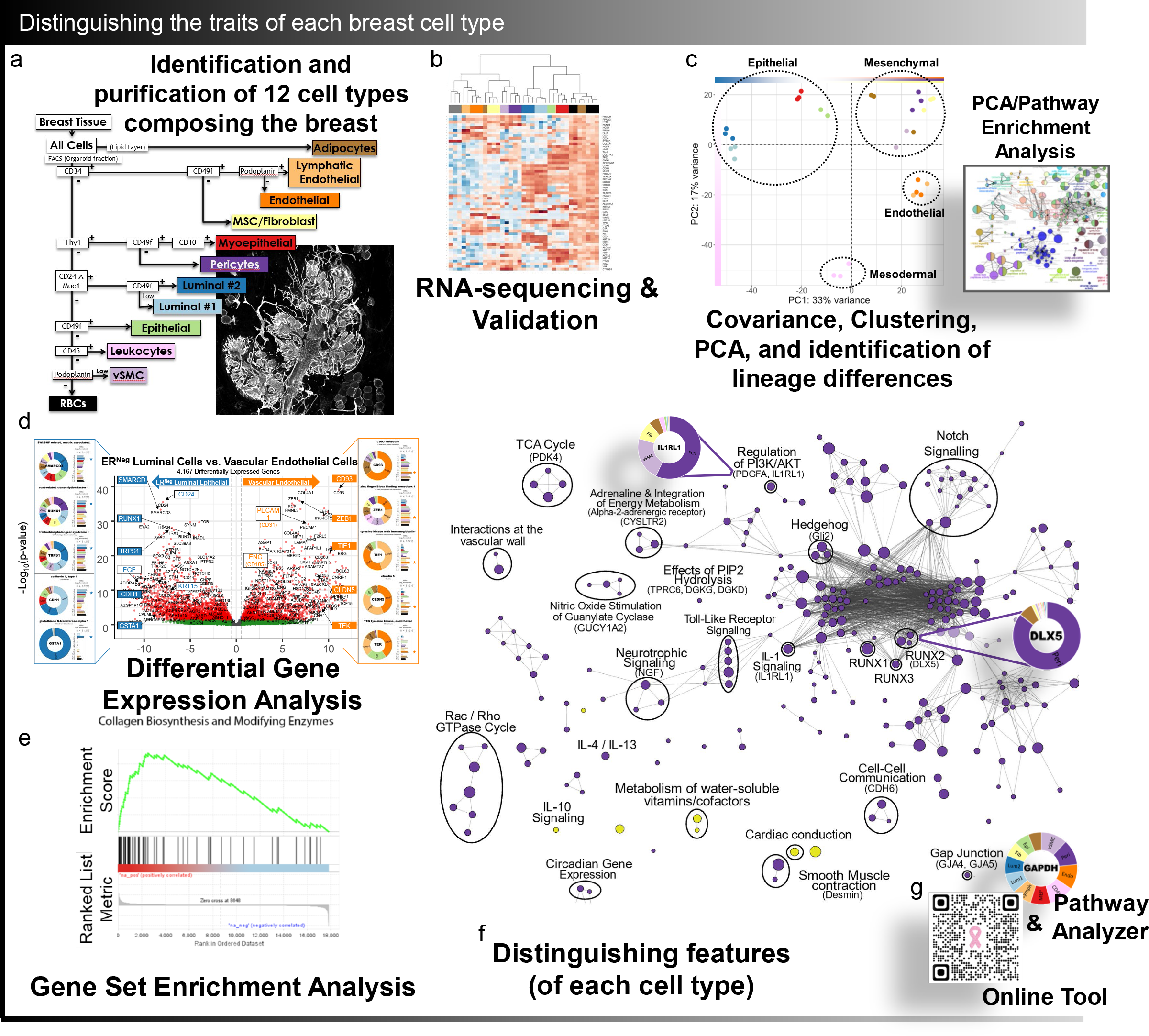
Distinguishing the traits of each breast cell type. Shown is a sketch of the RNA-sequencing workflow used for analyzing the eleven sequenced cell types. This began with **(a)** developing a FACS isolation method for separating each breast cell type (as detailed in Fig.1 and adjoining article). This procedure was scaled up and cells were sorted and RNA-sequenced. Analysis of the RNA-sequencing data involved: **(b)** validation and exploration of cellular relationships by hierarchical clustering, **(c)** covariance matrix and quality control analyses, identification of unique cell markers, and principal component analysis (PCA). PCA-derived gene lists were used to perform gene ontology enrichment analyses that revealed the shared characteristics of cell lineage. **(d)** Differential gene expression analysis was performed and used to identify gene families that varied most among breast cell types; Pairwise comparisons identified transcripts whose levels varied between sample pairs. Differentially expressed genes between cell pairs were ranked and analyzed by **(e)** Gene Set Enrichment Analysis, leading to the identification of key pathways and processes that differ between cell types. GSEA results were further analyzed by **(f)** Constructing and analyzing enrichment maps that identified the central biological themes and distinguishing features of each cell type. Analyzing the data required the development of software code and packages that are available for download: Pathway Analyzer (Open Science Framework, https://osf.io/), and online Transcript Finder (http://breastcancerlab.com/breast-atlas).

### mRNA sequencing and validation of FACS purified cell types

To investigate the unique biology of the breast cell types, we FACS isolated each population to sequence and analyze their transcriptomes. This process entailed processing and collagenase-treating breast tissues (reduction mammoplasty tissues free of proliferative disease), followed by low speed (<100g) centrifugation. The first population collected was the lipid layer that had phase-separated under centrifugal force (Pop12 adipocytes). All other cell types were FACS-purified from the pelleted tissue ‘organoids’ following their trypsinization to single-cell suspensions (Figure 1 and Figure 1-figure supplement 1). While developing and testing our FACS antibody panel, we typically stained between 100,000 to 500,000 cells^8^. However, many more were needed for mRNA library construction and sequencing here, so we scaled up our operation by increasing the number of cells stained by roughly 1000-fold.

FACS purifying cells for RNA-sequencing was performed on freshly collected tissues. This process required relatively large amounts of tissue (>30 grams) and sort times (14 to 18 continuous hours for each tissue) to produce cell yields that could support deep RNA-sequencing. In addition to the above lipid layer, the sorting provided 11 FACS-purified cell populations from each specimen. This exercise yielded hundreds of thousands to millions of cells for the more abundant cell types (Figure 3-table supplement 1). In contrast, yields were much lower for the rarer types, which produced a range of only 24,000-130,000 cells. After cell purification, the RNA isolated from each sample correlated with cell counts, and in pilot experiments, we found it essential to isolate RNA immediately following FACS to preserve its integrity. In two independent sequencing runs, we sequenced 36 samples representing eleven of the twelve identified breast cell types (red blood cells did not produce measurable RNA and thus were not sequenced). All cell populations were isolated from four random individuals (aged 16, 23, 27, and 29 years/old), and nearly all were sequenced in triplicate (from three of the four individuals, Figure 3—table supplement 2). In only two instances did we not have enough RNA of sufficient quality to sequence in triplicate –and both were rare populations with low FACS yields (Pop5 vascular smooth muscle cells and Pop7 lymphatic endothelial cells, Figure 1b, Figure 4a). We also isolated and sequenced RNAs from snap-frozen tissues and unsorted cell suspensions, which we used as mixed cell controls. After sequencing mRNAs from the samples and aligning the FASTQ sequencing data to the human genome, we calculated the average mRNA counts across all samples to be 39.5 million paired-end reads—all with Phred^12^ quality scores of 25 or more, indicating the sequencing was of high depth and quality. We found these early sequencing results encouraging and moved forward with quality control analyses.

**Figure 4.**
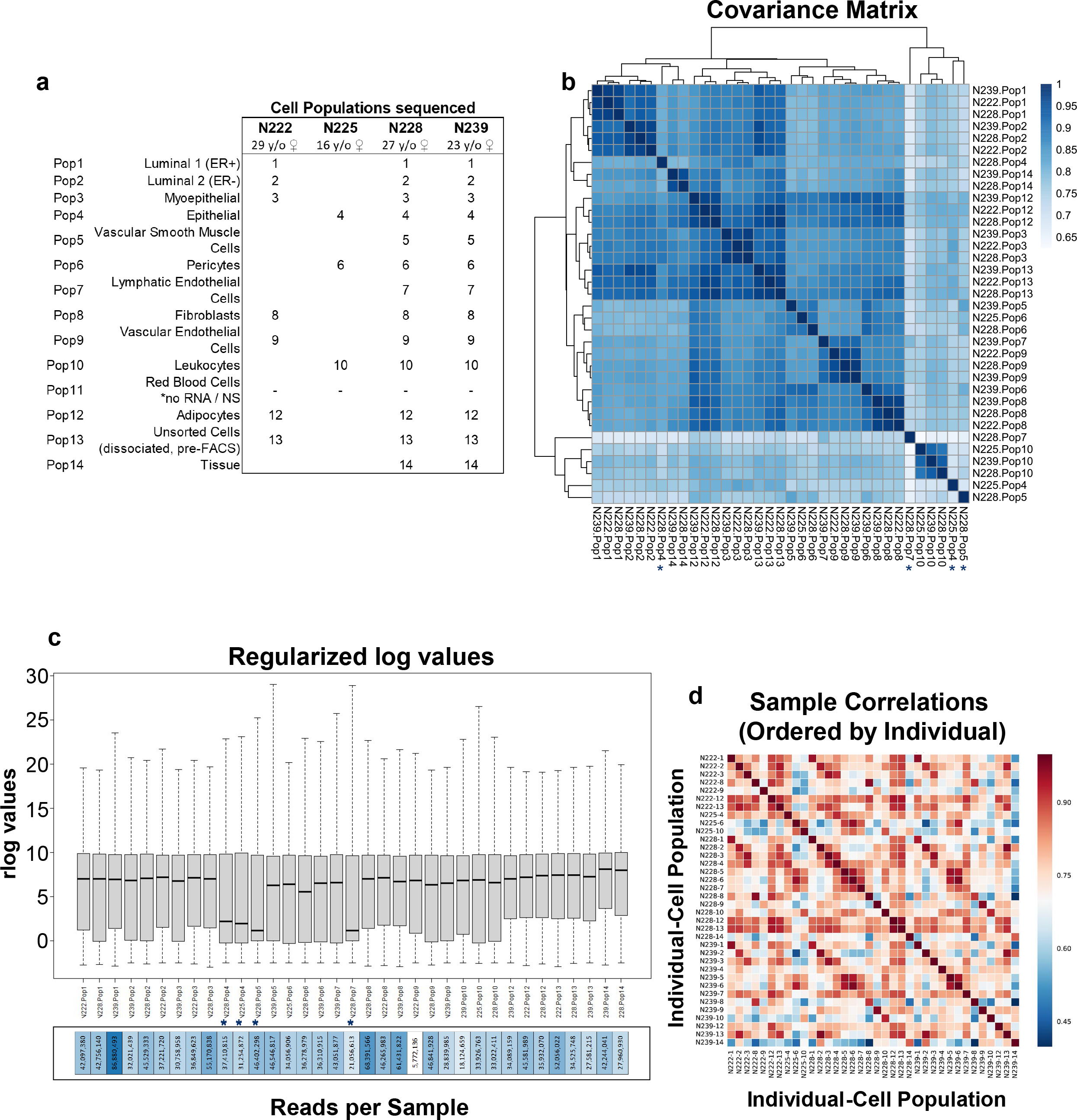
RNA-seq quality assessment. (a) Sample metadata for the mRNA-sequenced cell populations. **(b)** Covariance matrix displaying the correlation of gene expression for all pairwise combinations of samples. Samples N228-Pop4, N225-Pop4, N228-Pop5, N228-Pop7 have been flagged for lower correlations between replicates (*). These cell populations are the least abundant types in the breast and are among those that produced the lowest FACS cell yields **(c)** distribution of regularized log transcript levels in each sample. **(d)** Sample correlation matrix demonstrates absence of sample/batch effects.

Nearly all sequenced samples demonstrated excellent read counts. However, covariance analysis, assessment of Cook’s distances, and distribution of regularized log values^13^ identified five samples with reduced transcript complexity and lower correlations. All were the rarer cell types, i.e., Pop4 epithelial cells, Pop5 vSMCs, and lymphatic endothelial cells (Figure 4b,c).

Nevertheless, we determined that only Pop4 epithelial cells from individual N239 needed to be removed from critical downstream analyses. We retained the four other low-complexity samples because they clustered well with their cell-type counterparts (hierarchical clustering and PCA) and provided meaningful insights for many transcripts (Figure 4b, Figure 3—table supplement 2). These samples were: Pop4 epithelial cells (from individuals N225 and N228), Pop5 vSMCs (from N228), and Pop7 lymphatic endothelial cells (from individual N228). Although outlier genes existed in these samples, concordance between replicates was high for most others.

Despite these minor issues, the sequencing data across the board was impressively robust. We did not detect sequencing artifacts, such as correlations with sequencing run or individuals from which the cells were derived, as all samples clustered tightly based on their inherent cell type (Figure 4b,d). The substantial number of reads detected per sample, the highly correlated replicates, and the large number of unique transcripts detected in these specimens (21,559) indicated that the dataset was rich and of excellent quality. Indeed, these data were immediately helpful in determining the identities of several obscure cell populations.

Before we had sequenced the mRNAs from the cell types, there were several FACS populations whose identities were a mystery (as discussed in the adjoining article)^8^. Most notably were populations #4, 5, and 6, which we ultimately identified as a rare epithelial cell type, vascular smooth muscle cells, and pericytes. RNA-sequencing provided conclusive evidence that helped identify these cell types by revealing their relationships to other populations and supplying information on expressed markers. After identifying these three populations, we explored all remaining cell types. Reassuringly, transcript levels of genes encoding cell surface markers used to sort each cell type tracked nicely with their respective FACS-staining levels (Figure 4—figure supplement 1a,b; Figure 1a). And GAPDH, often used as a uniformly-expressed internal control gene, was indeed expressed equally among the cell types, as were the transcripts of other widely-used internal control genes (Figure 4—figure supplement 1c, Figure 3—table supplement 2). In addition, analysis of hundreds of other genes, such as those uniquely expressed by each cell type or those used as cell type markers, i.e., genes encoding CD antigens, validated and helped us build confidence in our dataset (Figure 5, Figure 5—figure supplement 1). To facilitate the exploration of these RNA-sequencing data, we have created a publicly available tool for visualizing transcript levels in each cell type (http://breastcancerlab.com/breast-atlas/).

**Figure 5.**
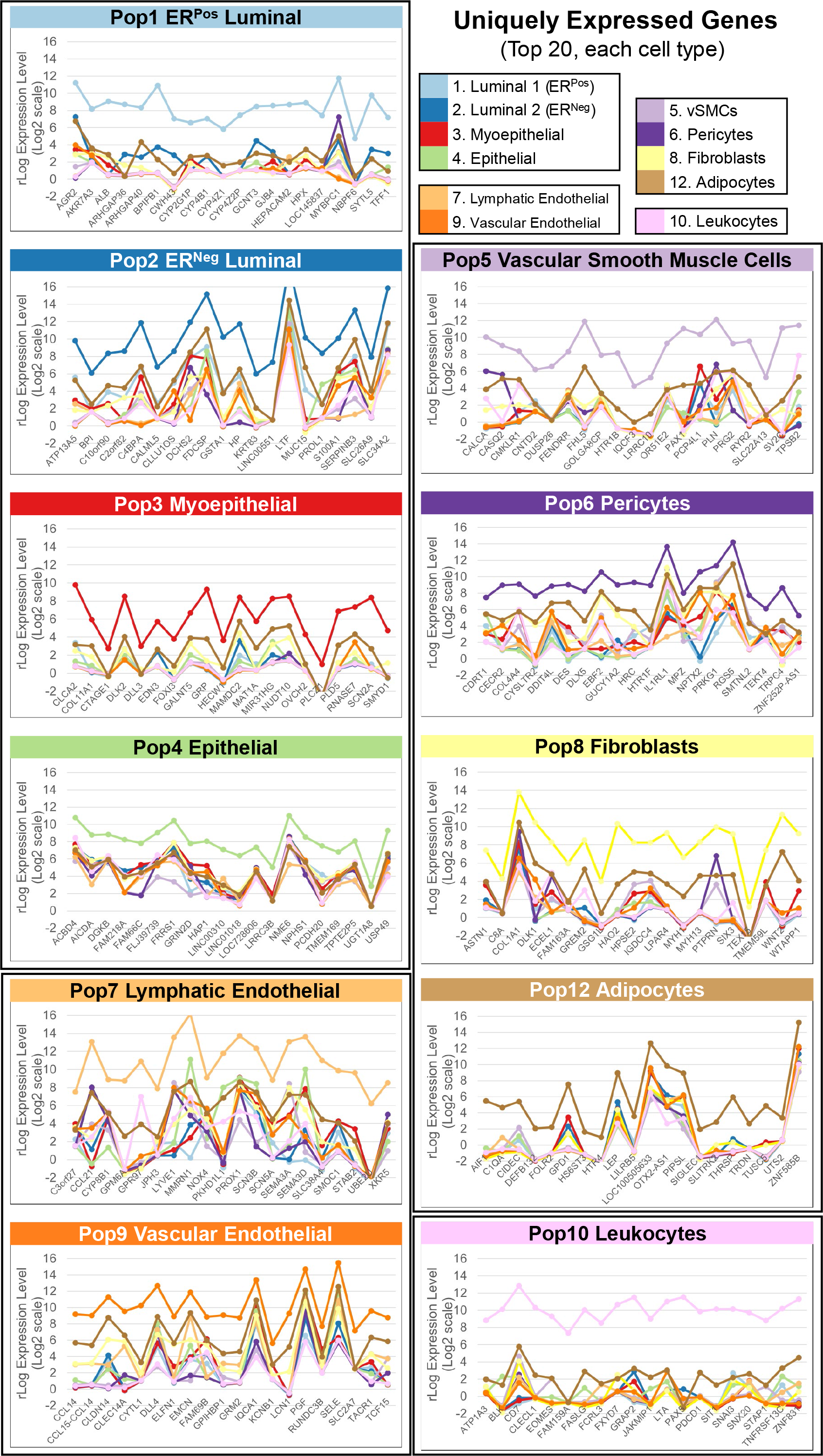
Uniquely expressed genes. A sliding threshold (fold-change) was used to identify the top 20 most uniquely expressed genes in each of the eleven sequenced cell types. Notable genes included: **a)** albumin (ALB) and connexin 30.3 (GJB4) in ER-positive luminal cells, **(b)** lactotransferrin (LTF) in ER-negative luminal cells, **(c)** alpha-1 chain of collagen XI (COL11A1) and notch ligand delta-like 3 (DLL3) in myoepithelial cells, **(d)** protocadherin 20 (PCDH20) in Pop4 epithelial cells, **(e)** lymphatic vessel endothelial hyaluronan receptor 1 (LYVE1) and prospero homeobox 1 (PROX1) in lymphatic endothelial cells, **(f)** endothelial selectin (SELE) and delta like canonical notch ligand 4 (DLL4) in vascular endothelial cells, **(g)** calcitonin related polypeptide alpha (CALCA) and 5-hydroxytryptamine receptor 1b (HTR1B) in vascular smooth muscle cells, **(h)** desmin (DES) and 5-hydroxytryptamine receptor 1f (HTR1F) in pericytes, **(i)** alpha-1 chain of collagen I (COL1A1) and wnt ligand family member 2 (WNT2) in fibroblasts, **(j)** the lipid-binding protein ‘cell death inducing dffa-like effector c’ (CIDEC) and glycerol-3-phosphate dehydrogenase 1 (GPD1) in adipocytes, and **(k)** CD7 and fas ligand (FASLG) in leukocytes.

### Identification of cell-type specific transcriptional patterns

One of our early goals was to identify the distinct transcriptional patterns of the different breast cell types. To address this question and further explore the dataset, we clustered the samples’ transcriptomes using a standard Euclidean distance metric. Hierarchical clustering recapitulated the distinct nature of the different cell types and mixed cell controls (Figure 6). Clustering was robust, especially for the two luminal cell types, myoepithelial cells, pericytes, fibroblasts, vascular endothelial cells, and adipocyte samples (all of which displayed approximately unbiased (AU) *p-*values >0.95 when evaluated by multiscale bootstrap resampling^14^). Notably, biological replicates of each cell type clustered tightly together, and similar cell types formed adjacent clusters. These related pairs included: a) vascular and lymphatic endothelial cells (FACS Pops 7&9), b) vSMCs and pericytes (Pops5&6), c) ER^Pos^ and ER^Neg^ luminal epithelial cells (Pops 1&2), and d) myoepithelial cells and related Pop4 epithelial cells (Pops3&4). As expected, fibroblasts (Pop8) clustered with the other mesenchymal cell types (vSMCs and pericytes)— albeit at a greater distance— reflecting their inherently distinct features. In contrast, the most distant samples that stood far from the others were the CD45^Pos^ leukocytes (Pop10)—a separation that we anticipated, given their unique biology. The clustering confirmed that the eleven sorted cell types were indeed distinct and captured the underlying differences—and similarities— in their gene expression patterns.

**Figure 6.**
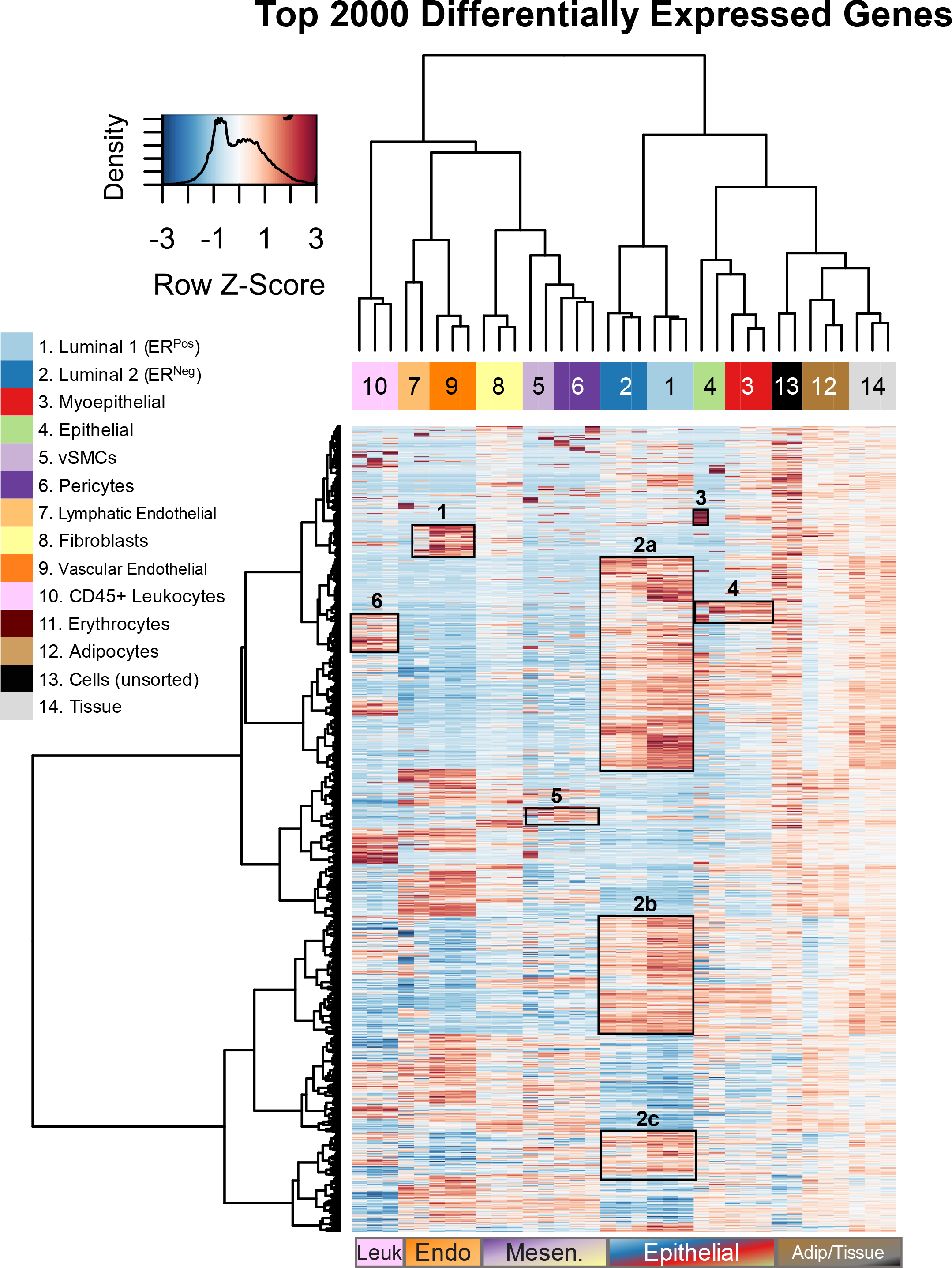
Transcript levels of top 2000 most differentially expressed genes. Heatmap with dendrogram of the top 2000 most differentially expressed genes across all FACS sorted populations, as well as unsorted cells and matched breast tissues (mixed cell controls). Clustering was performed using Euclidean distance measures and Ward clustering method on variance stabilized expression values (DESeq2; Variance Stabilized Transformation, VST). Adjusted p-values for these 2000 differentially expressed genes ranged between 4.9e-97 to 0.0017. All 2000 genes symbols can be viewed in Figure 3—table supplement 5 (when the data is ordered by adj. *p*-value).

Several features emerged after closely examining the hierarchical clustering of genes. Gene expression trends, in particular, were apparent across populations, and many clusters contained genes encoding prominent cell markers, which helped us identify or confirm each cell type’s identity (Figure 6). For example, p-selectin (SELP) and HOXD1 were genes in a cluster that differentiated endothelial cells. Epithelial cell adhesion molecule (EPCAM), zona occludens 3 (TJP3), and epidermal growth factor (EGF) were among the many genes in a cluster highly expressed by luminal epithelial cells. Calmodulin-like 3 (CALML3) and G-protein-coupled receptor 87 (GPR87) were highly expressed by Pop4 epithelial and myoepithelial cells, along with the chloride channel regulator CLCA2, which is regulated by the myoepithelial lineage marker p63 and is involved in basal cell adhesion. Finally, the perivascular cell cluster contained angiopoietin 1 (ANGPT1) and connexin 40 (GJA5), which are essential for blood vessel development and maintenance. The clustering thus demonstrated high agreement among the biological replicates and identified clusters of shared genes that provided substantial evidence confirming the identity of each cell population (or pointing to its identity).

The hierarchical clustering used to compare cell types was also notably stable. Whereas Figure 6 shows the clustering of the 2000 most differentially expressed genes, the pattern did not significantly change when we repeated the analysis with different inclusive thresholds, such as when we used the top 100, 200, 500, 1000, or 3000 genes that were most differentially expressed. Although the cellular relationships that emerged from these analyses were largely confirmatory, they helped us identify and characterize several populations early on. For example, before we had ascertained the identity of FACS Pop5 as vSMCs, its close clustering with related Pop6 pericytes here provided a critical clue^8^. The clustering also revealed similarities between the rare Pop4 epithelial cells and myoepithelial cells. However, it also showed that Pop4’s expression of some genes more resembled those found in luminal epithelial cells that distinguished this cell population from myoepithelial cells^8^. The molecular patterns that emerged provided information on each population and objective evidence of the relatedness among cell types. To explore these relationships—and features fundamental to each cell type—we needed to decipher their transcriptomes in finer detail.

### Distinguishing the features of cell lineage

Differentiated cells are often classified by their developmental history and the lineage from which they are derived. The different breast cells, for example, are often described as being mesenchymal (fibroblasts, pericytes, vSMC, and adipocytes), hematopoietic/blood (leukocytes, erythrocytes), endothelial (vascular and lymphatic), or epithelial derived (ER^Pos^ and ER^Neg^ luminal, myoepithelial, and Pop4 epithelial, Figure 2b). Although our primary goal was to identify the qualities that make each cell type unique, our sequencing of every breast cell type presented an unprecedented opportunity to explore the broad attributes of cell lineage through the comparative lens afforded by our dataset’s comprehensive nature. We accomplished this through principal component and pathway analysis, as this approach allowed us to analyze the samples en masse—and not focus on any particular pairwise comparison prematurely. As a result, these analyses produced an astonishing amount of data. Unfortunately, we can present only a fraction of it here. However, we have provided additional tools and tables to aid the exploration of all data and results presented herein (Figure 3—table supplements, Pathway Analyzer^9^, and the Breast Cell Atlas: Transcript Finder App^10^). Below are our analyses of the first four principal components that revealed the distinguishing features of the four broad cell lineages composing breast tissues.

Principal component analysis (PCA) of the transcripts from all cell types showed gene expression variances were distributed across multiple principal components (PCs), with the bulk of the variation (71%) being captured by the first four components (Figure 7—figure supplement 1). The projection of principal component 1 (PC1) and principal component 2 (PC2) markedly separated the four cell lineages in two-dimensional space. Genes contributing to PC1 were primarily responsible for distinguishing the mesenchymal and epithelial cell types, as these lineages projected at PC1’s opposing ends (Figure 7a). Interestingly, endothelial cells—which derive from mesenchyme but are classically regarded as a specialized epithelial cell type^15, 16^— projected as an independent group that aligned more closely with the mesenchymal cell types along PC1. In contrast, the variance explained by genes composing the second principal component (PC2) emerged from leukocytes, as these cells formed a single group at its distal end, far apart from the other cell types (Figure 7a). A similar analysis of principal components 3 and 4 resolved the endothelial cells and two subtypes of the mesenchymal and epithelial lineages, i.e., perivascular cells and myoepithelial/Pop4 epithelial cells (Figure 8a).

**Figure 7.**
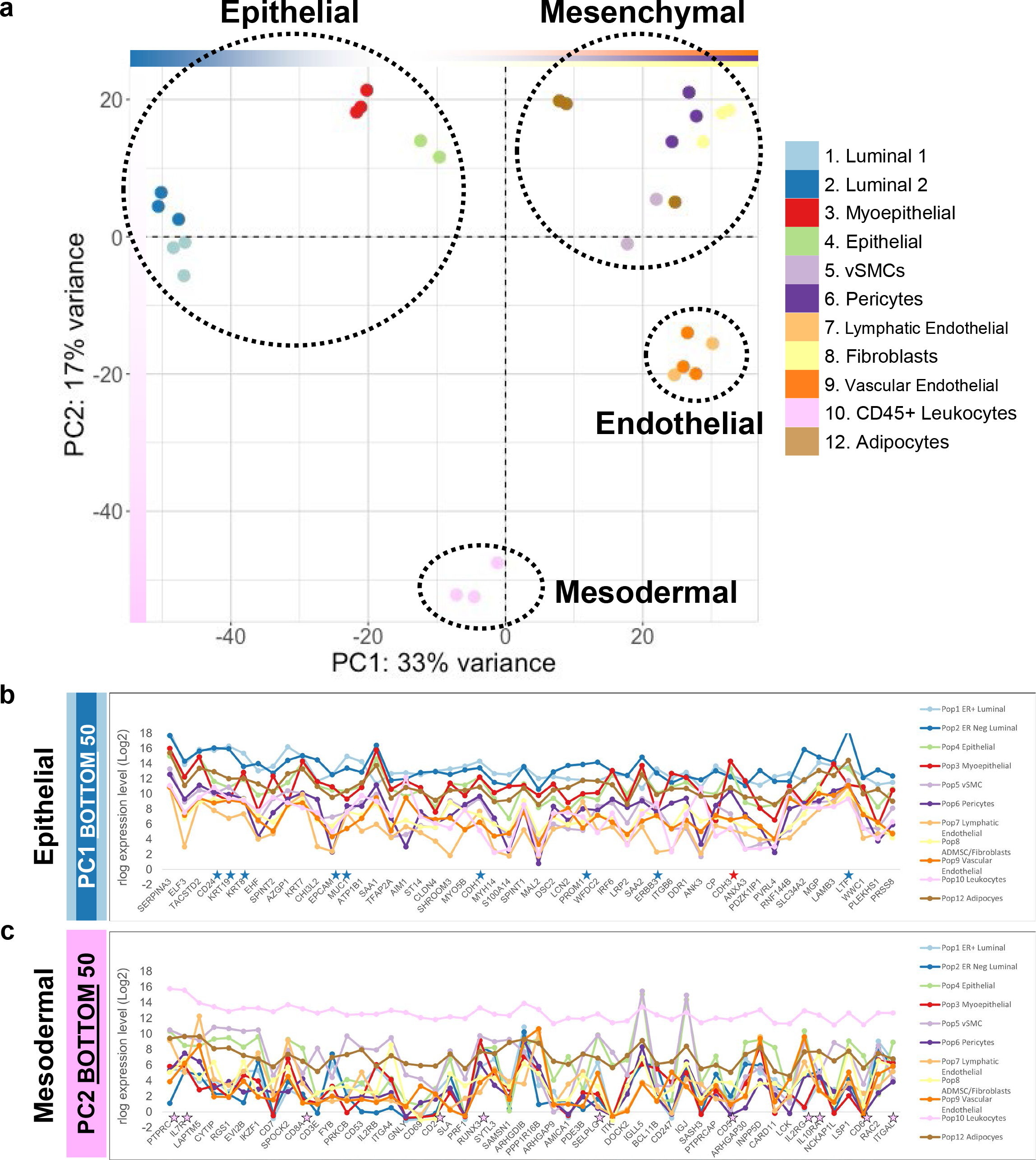
Principal Component Analysis. (a) projection of breast cell types on principal components 1 and 2 (PC1, PC2). PC1 separates the luminal ER^Pos^ and ER^Neg^, Pop4 epithelial, and myoepithelial cell types (epithelial lineages) from the mesenchymal (fibroblasts, adipocytes, pericytes and vSMCs) and endothelial (lymphatic and vascular) lineages. PC2 separates the CD45^Pos^ leukocytes (mesodermal lineage) from all other cell types. The cell types composing each of the epithelial, mesenchymal, endothelial, and mesodermal cell-lineages are indicated (dashed circles). **(b)** Median transcript levels of the 50 loading factors (genes) that contribute to the negative end of PC1(bottom end of the list, rlog/log2 scale). **(c)** Median mRNA expression of the 50 loading factors (genes) contributing to the negative end of PC2 (bottom end of the list) that separates the leukocyte samples (rlog, log2 scale). Stars (↔) denote commonly used markers for luminal cells (blue), myoepithelial cells (red), and leukocytes (pink).

**Figure 8.**
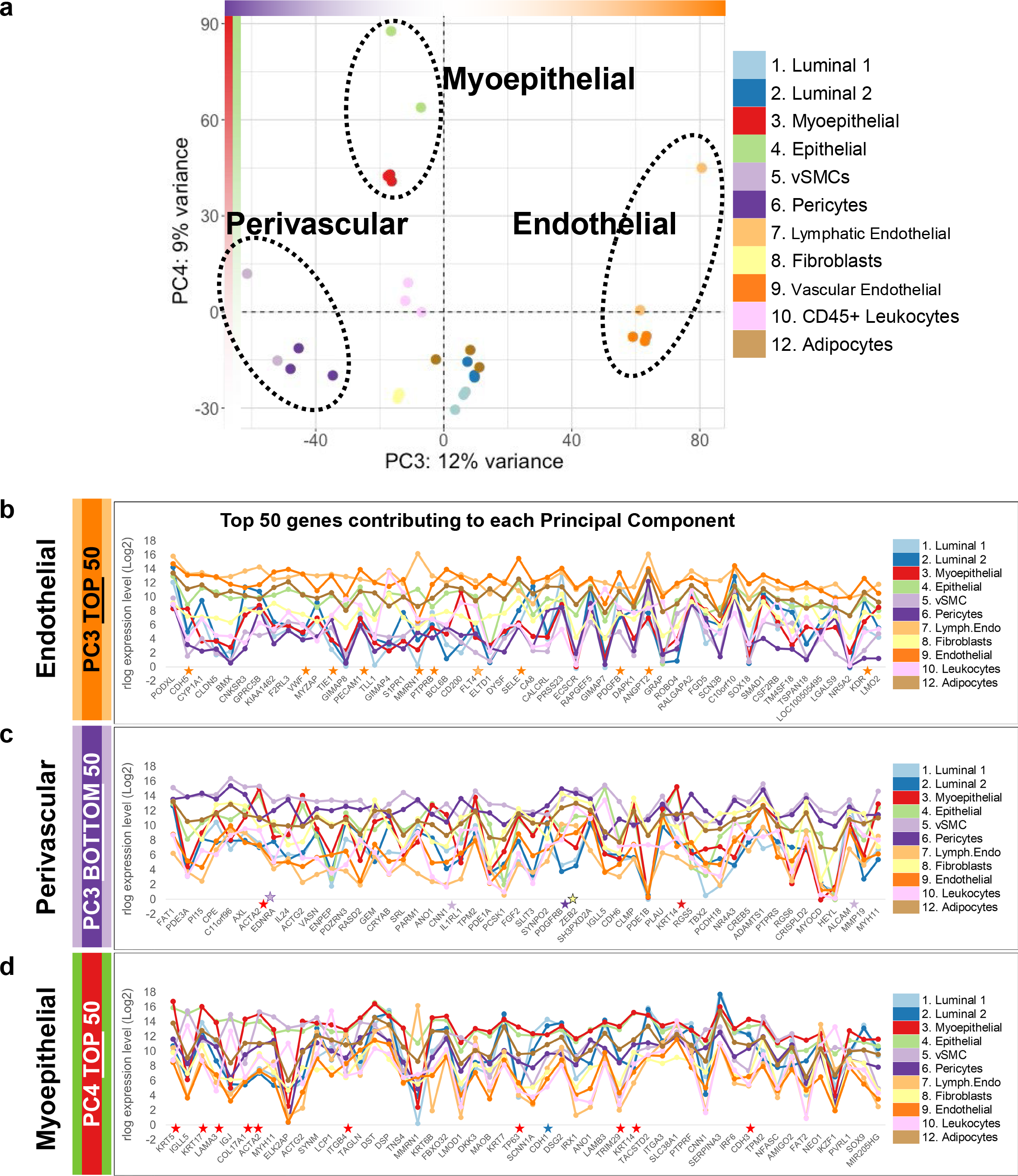
PCA plot of PC3 and PC4. (a) PCA projection of breast cell types on principal components 3 and 4 (PC3, PC4). PC3 separates the endothelial (most positive) and perivascular (most negative) cell types, whereas PC4 separates the myoepithelial and myoepithelial-related cell types (myoepithelial and Pop4 epithelial cells). Median mRNA expression of the 50 loading factors contributing to the **(b)** positive end of PC3 (top of the list; rlog, log2 scale), **(c)** negative end of PC3 (bottom of the list; rlog, log2 scale), and **(d)** the positive end of PC4 (top of the list; rlog, log2 scale). Stars (↔) denote common markers of endothelial cells (orange), myoepithelial cell (red), fibroblasts (yellow), pericytes (purple), and luminal cells (blue).

To gain further insights into the distinguishing attributes shared by members of each cell lineage, we examined the genes impacting their PCA groupings, i.e., the loading factors with the highest influence on each principal component (Figure 3—table supplement 3). We then used these unique gene lists to probe functionally grouped Gene Ontology^17^ and Reactome^18^ pathway annotation networks^19^.

#### **Epithelial Lineage** (ER^Pos^ & ER^Neg^ Luminal cells, Myoepithelial cells, Pop4 Epithelial cells)

We first analyzed the cluster defined by the epithelial lineages: the ER^Pos^ and ER^Neg^ luminal myoepithelial and Pop4 epithelial cell types (Figure 2b). Based on their positioning on PC1, we expected the genes with the highest contributions to PC1 to reflect those that distinguish the epithelial and mesenchymal lineages, and in particular, highlight genes associated with luminal epithelial cells—as the ER^Pos^ and ER^Neg^ luminal cells projected the farthest along on PC1. Analysis of gene expression in each cell type showed this was indeed the case (Figure 7b). Many of PC1’s top genes (at the negative epithelial end) were recognizable, as they are often used as markers of the epithelial lineage. They included keratins 6b, 8, 18, and 19, P-and E-cadherins, claudin 4, and Her2 (Figure 7b, Figure 9a-e; Figure 3—table supplement 3). Other familiar genes included integrin beta 6, the transcription factors ELF3, EHF, and TFAP2A; peptidase inhibitors SERPINA3 and SPINT2; serine protease ST14, cytoskeletal and associated proteins SHROOM3, MYO5D, and MYH14; the calcium-binding protein S100A14, among others (Figure 3— table supplement 3). These genes were among the 200 extracted from the negative epithelial end of PC1—a gene list we used to identify the biological pathways enriched within the epithelial cell lineage.

**Figure 9.**
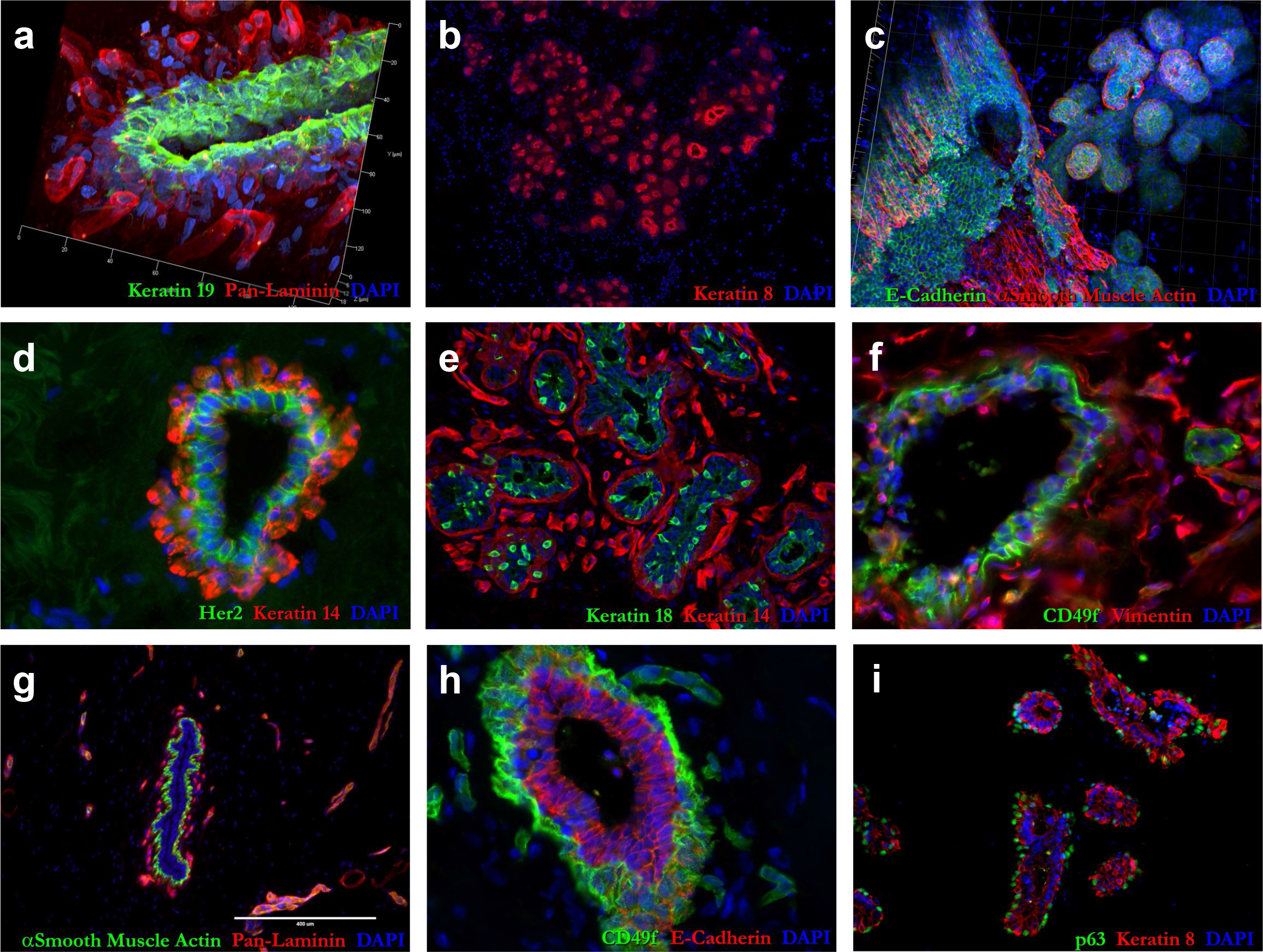
IHC validation of PCA markers. Tissue immunostaining of normal human breast tissue with markers contained in the principal components. **(a)** keratin 19 (green) and pan-laminin (red); **(b)** keratin 8 (red); **(c)** e-cadherin (green) and alpha smooth muscle actin (red); **(d)** Her2 (green) and keratin 14 red); **(e)** CD49f (green) and vimentin (red); **(f)** alpha smooth muscle actin (green) and pan-laminin (red); **(g)** CD49f (green) and e-cadherin (red); and **(h)** p63 (green) and keratin 8 (red). Loading factors at the negative end of PC1 (epithelial end) included: keratin 19/Krt19 (#5 on list), keratin 8/KRT8 (#6), E-cadherin/CDH1 (#22), Her2/ERBB2 (#85), and keratin 18/KRT18 (#138), and CD49f/ITGA6 (#5574). Loading factors at the positive end of PC1^No^ ^Endo^ (mesenchymal end) included: smooth muscle actin/ACTA2 (#155 on the list), and Vimentin/VIM (#192). Loading factors at the negative end of PC3 (perivascular end) included: smooth muscle actin/ACTA2 (#7 on the list). Loading factors at the positive end of PC3 (endothelial end) included: CD49f/ITGA6 (#344). Loading factors at the positive end of PC4 (myoepithelial end) included: smooth muscle actin/ACTA2 (#7 on the list), P63/TP63 (#25) and CD49f/ITGA6 (#68).

Using the epithelial gene set, we performed a two-sided hypergeometric (enrichment/depletion) test to determine if particular molecular functions or biological pathways were disproportionately represented (Gene Ontology^20^, Reactome Pathway database^18^). We found many were, and we aggregated the results to form a functionally organized pathway network, which grouped similar biological processes based on their shared genes (ClueGO^21^). The 124 enriched gene sets aggregated into twenty-four distinct groups related to cell adhesion, peptidase activity, keratinization, estrogen signaling, AP-2 transcription factor regulation, mucins, and innate immune system factors (Figure 10a, Figure 3—table supplement 4: PC1 Epi tab). The most significant group (*p*-value =1.4x10^-8^) contained the *cell adhesion molecule binding* and *cadherin binding* Gene Ontology terms. These gene sets represent different levels of the same ontology hierarchy and are highly correlated. For example, they both contain components of adherens junctions, i.e., epithelial- and placental-cadherins (CDH1 and CDH3), cingulin (CGN), and 348 other genes. By comparison, the *cell adhesion* gene set contains an additional 196 genes related to other aspects of cellular adhesion, such as desmosome-associated desmoplakin (DSP) and desmoglein 2 (DSG2, Figure 3—table supplement 3, sorted by PC1). The algorithm’s ability to cluster overlapping gene sets like these, and assemble them into related clusters, allowed us to extract the common biological themes obscured within the volume and complexity of the data.

**Figure 10.**
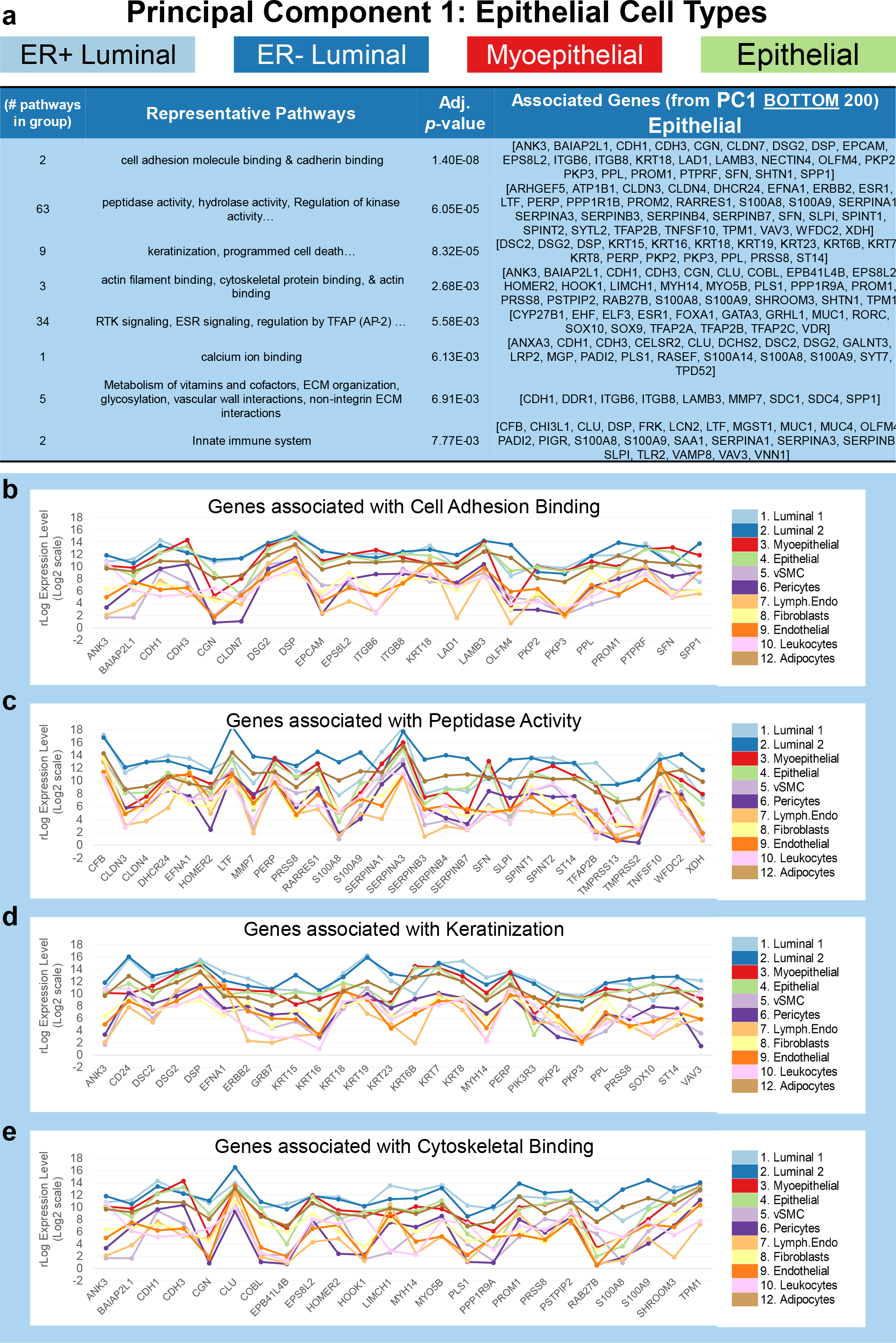
Pathway analysis of epithelial genes (PC1). (a) Grouped GO/Reactome terms overrepresented within the gene list 200 genes contributing to the negative/epithelial end of PC1 (bottom of the list) are ordered by adjusted p-value. Grouping was done by the similarity of terms to evaluate broad differences in cell class. Associated genes for each group are listed in the 4^th^ column. Median mRNA expression (rlog, log2 scale) for PC1 genes associated with select pathways including **(b)** c*ell adhesion and binding*, **(c)** *peptidase activity*, **(d)** *keratinization*, and **(e)** *cytoskeletal binding*.

Reassuringly, when we examined the expression of the genes in these pathways, we found they were indeed elevated in the four epithelial cell types (Figure 10b-e, (Figure 3—table supplement 4: PC1 Epi tab). These enriched pathways and themes describe general features of the epithelial lineage. After exploring the remaining cell lineages’ shared attributes (defined by each PC), we will revisit these cells and explore their individual traits.

#### **Leukocytes** (CD45^Pos^ Cells)

The second principal component separated the leukocytes (CD45^Pos^ FACS population), and analysis of PC2’s loading factors revealed many well-known leukocyte markers (Figure 7a,c). For example, ‘protein tyrosine phosphatase receptor type C’ (PTPRC), the gene encoding the pan-leukocyte marker CD45—which we used to FACS isolate the leukocytes—was notably the first gene on the list. Other top genes included the CD antigens: CD2, CD3E, CD6, CD7, CD8A, and others. Also present was the transcription factor RUNX3, which is critical for T-cell differentiation, as was selectin P ligand (SELPLG), which is crucial for leukocyte trafficking. Using the same technique described above (for PC1 and the epithelial cells), we extracted the top genes from the leukocyte-defining end of PC2 and performed pathway enrichment analysis. We identified 453 enriched gene sets that clustered into 62 distinct thematic groups (Figure 3—table supplement 4: PC2 Leukocyte tab). Representative pathways from the ten most significant groups were: *adaptive immune response*, *activation of immune response*, *leukocyte homeostasis, T-cell activation, leukocyte chemotaxis, phagocytosis, IL-6 production, acute inflammatory response, regulation of GTPase activity, mast cell degranulation, and monocyte differentiation* (*p-*values ranged between 6.07x10^-46^ – 9.53x10^-11^, Figure 10—figure supplement 1a). Although most leukocytes in the breast are macrophages^8^, the sequenced mRNAs from the CD45^Pos^ FACS gate represent a diverse population of white blood cells. Therefore, finding a broad range of enriched pathways attributable to different leukocyte subpopulations was expected. When we compared the expression of enriched genes in each cell type, we found they were indeed highly expressed by the Pop10 leukocytes (Figure 10—figure supplement 1b-d).

#### ***Mesenchymal Lineage*** *(vSMCs, Pericytes, Fibroblasts, Adipocytes)*

Similar gene expression patterns within the four mesenchymal cell types positioned this cell group toward the positive end of PC1’s axis in the 2D projection of PC1 and PC2 (Figure 7a). However, endothelial cells were also located in this region, indicating they and mesenchymal cells share the distinct expression of many genes. The relatively similar projection of endothelial and mesenchymal cells along PC1 prevented extraction of loading factors that could be solely attributable to either cell class. As we were primarily interested in identifying the defining characteristics of mesenchymal cells at this stage, we digitally removed the endothelial cell data and repeated the principal component analysis. The new PCA projection resolved the mesenchymal cells, situating them at the positive end of the first principal component in this ancillary graph (PC1^No^ ^ENDO^, Figure 7—figure supplement 2a). Exploring the loading factors of this new principal component, we confirmed that these genes were indeed highly expressed by the mesenchymal cells. For example, vimentin (VIM), an intermediate filament protein and classical mesenchymal marker (Figure 9f), was near the top of the list, at position 192 (out of 21,560 genes, Figure 7— figure supplement 2b). The top loading factor was the non-selective receptor for endothelins (EDNRB). Endothelins are potent vasoactive peptides, and EDNRB expression was notably highest in the perivascular cell types (vSMC and pericytes) –these cells surround blood vessels and regulate vessel diameter by influencing vasoconstriction and dilation. Genes highly contributing to PC1 included many collagen family members (e.g., COL4A1, COL1A2, COL6A3, COL4A2, COL3A1, COL15A1, COL5A3, and others). Also present were ADAM metalloproteinases, the transcription factors ZEB1and ZEB2, platelet-derived growth factor receptor beta (PDGFRB), insulin-like growth factor II, laminin subunit alpha-4 (LAMA4), interleukin 24 (IL24), chondroitin sulfate proteoglycan 4 (CSPG4), along with many other notable genes (Figure 7— figure supplement 2b, Figure 3—table supplement 3: PC1^NoEndo^). To understand how these genes collectively contribute to mesenchymal cells, we performed pathway analysis using the loading factors from the mesenchymal-defining end of PC1^No^ ^ENDO^.

Pathway enrichment using 200 of PC1’s most critical mesenchymal genes identified 337 enriched gene sets from the Reactome and Gene Ontology databases (*p*-values< 0.05). These pathways assembled into a network of 96 distinct thematic groups (Figure 3—table supplement 4: PC1 Mesenchymal tab). The most significant group (*p*-value =5x10^-33^) contained 94 different aggregated gene sets/pathways that dealt mainly with: cardiac development (chamber, muscle, septum), cartilage morphogenesis, smooth muscle proliferation, bone development, and chondrocyte differentiation—pathways that are consistent with our primary cultures showing the isolated breast fibroblasts have multi-lineage differentiation potential^8^. The defining genes in this group included TBX2 and HEYL transcription factors, the type III TGFβ receptor TGFBR3, fibroblast growth factor FGF2, ADAM metallopeptidase with thrombospondin type 1 motif 1a and 9 (ADAMTS1, ADAMTS9); platelet-derived growth factor receptor beta (PDGFRB), and others (Figure 11a). Representative pathways from the other top significant groups included: *Fibroblast migration, GAG metabolic process, chondrocyte development, bone development, smooth muscle contraction, negative regulation of calcium ion transport, regulation of fatty acid biosynthetic process, and dermatan sulfate biosynthesis,* with *p-*values ranging between 5.00x10^-33^ to 4.01x10^-08^ (Figure 11a, Figure 3—table supplement 4: PC1 Mesenchymal tab).

**Figure 11.**
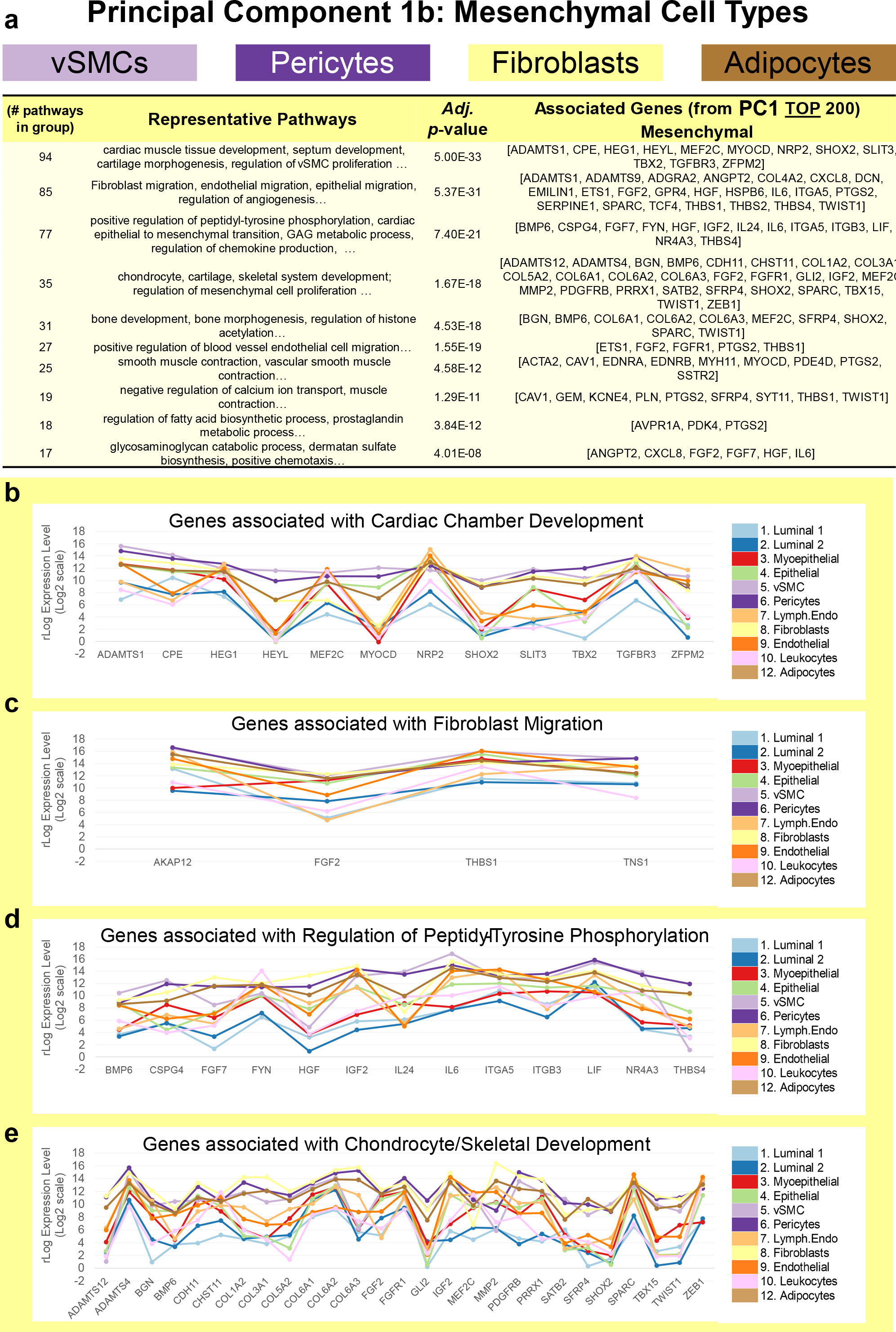
Pathway analysis of mesenchymal genes (PC1^No^ ^Endo^). (a) Grouped GO/Reactome terms overrepresented within the gene list 200 genes contributing to the positive/mesenchymal end of PC1^NoEndo^, ordered by adjusted p-value. Median mRNA expression (rlog, log2 scale) for PC1^NoEndo^ genes associated with select pathways, including **(b)** *cardiac tissue development*, **(c)** *fibroblast migration*, **(d)** *regulation of peptidyl-tyrosine phosphorylation*, and **(e)** *chondrocyte/skeletal development*.

While the four mesenchymal cell types highly expressed genes in the enriched pathways, transcript levels were not always uniform across these cell types (Figure 11b-d). For example, in the *cardiac chamber development* gene set, expression of the pathway’s genes was consistently higher in the perivascular cell types. The heart is composed of mesenchymal cardiac muscle cells and is a highly vascularized tissue that requires perivascular support, which may explain the frequent enrichment of cardiac-related pathways within the mesenchymal gene set (Figure 11— cardiac chamber development). The enrichment with these and other tissue-related gene sets (cartilage, bone, muscle, and adipose) indicates a deeply-rooted bond among mesenchymal cells and common dependence on a core set of genes. Later, we will use specific pairwise comparisons to explore the distinguishing features of the different mesenchymal cell types.

#### ***Endothelial Lineage*** *(Lymphatic and Vascular Endothelial Cells)*

To identify the genes and processes responsible for shaping the unique characteristics of endothelial cells, we explored the third and fourth principal components. These components resolved the vascular and lymphatic endothelial cells as a distinct group near the positive end of PC3 (Figure 8a). Many genes contributing to the endothelial end of PC3 were familiar, such as vascular-endothelial cadherin (CDH5)—a prominent endothelial adhesion molecule (Figure 8b). We also observed von Willebrand factor (VWF); the angiopoietin regulator tie1 (TIE1); the endothelial marker CD31 (PECAM1); endothelial selectin (SELE); angiopoietin 2 (ANGPT2); as well as vascular endothelial growth factor receptors KDR and FLT4, and many other notable genes. The top gene on the list was ‘podocalyxin like’ (PODXL), which binds leukocyte selectin and is vital to lymphocyte-endothelial adhesion. Also present was Claudin 5, which is crucial for endothelial cell junctions. Finding so many genes with known importance to endothelial cells corroborated these genes’ notable contributions to endothelial cells and showed they are characteristics of both vascular and lymphatic cell types. At the same time, the analysis identified other influential genes that are perhaps less appreciated or whose importance to endothelial cell function are not as well characterized (listed in Figure 9, Figure 3—table supplement 3). To explore the biological contribution of these genes to endothelial cell biology, we performed pathway enrichment analysis.

Hypergeometric analysis of the endothelial genes from PC3 identified 650 enriched biological pathways (*p-*values< 0.05, Figure 3—table supplement 4: PC3 Endothelial tab). These pathways assembled into 61 groups based on the genes they shared. Not surprisingly, most groups pertained to blood vessel and circulatory system biology (Figure 12a). The most significant group (*p-*value =6.27x10^-19^) involved cell motility and chemotaxis and contained *ameboidal-type cell migration*, *angiogenesis*, and *signaling by VEGF* gene sets (Figure 12a,b). As expected, PC3-identified genes within these enriched pathways were highly and differentially expressed by the lymphatic and endothelial cell types (Figure 12b-d). Representative pathways from these groups included: *cell-matrix adhesion, endothelial- and muscle-cell differentiation, lymph vessel morphogenesis, regulation of establishment of endothelial barrier, regulation of coagulation, vasculogenesis, and monocyte chemotaxis.* Many notable genes were present in these pathways, including podoplanin (PDPN) —a marker we used to sort lymphatic endothelial cells— ephrins A1 and B2 (EFNA1 and EFNB2), activin a receptor like type 1 (ACVRL1), platelet-derived growth factor subunit b (PDGFB), along with the aforementioned cadherin 5, claudin 5 and the VEGF receptors (KDR and FLT4). Finding vascular-related pathways enriched in the endothelial data set was not surprising, but identifying these pathways was reassuring, as it validated the approach and indicated these processes were occurring in both lymphatic and vascular endothelial cell types found in the breast.

**Figure 12.**
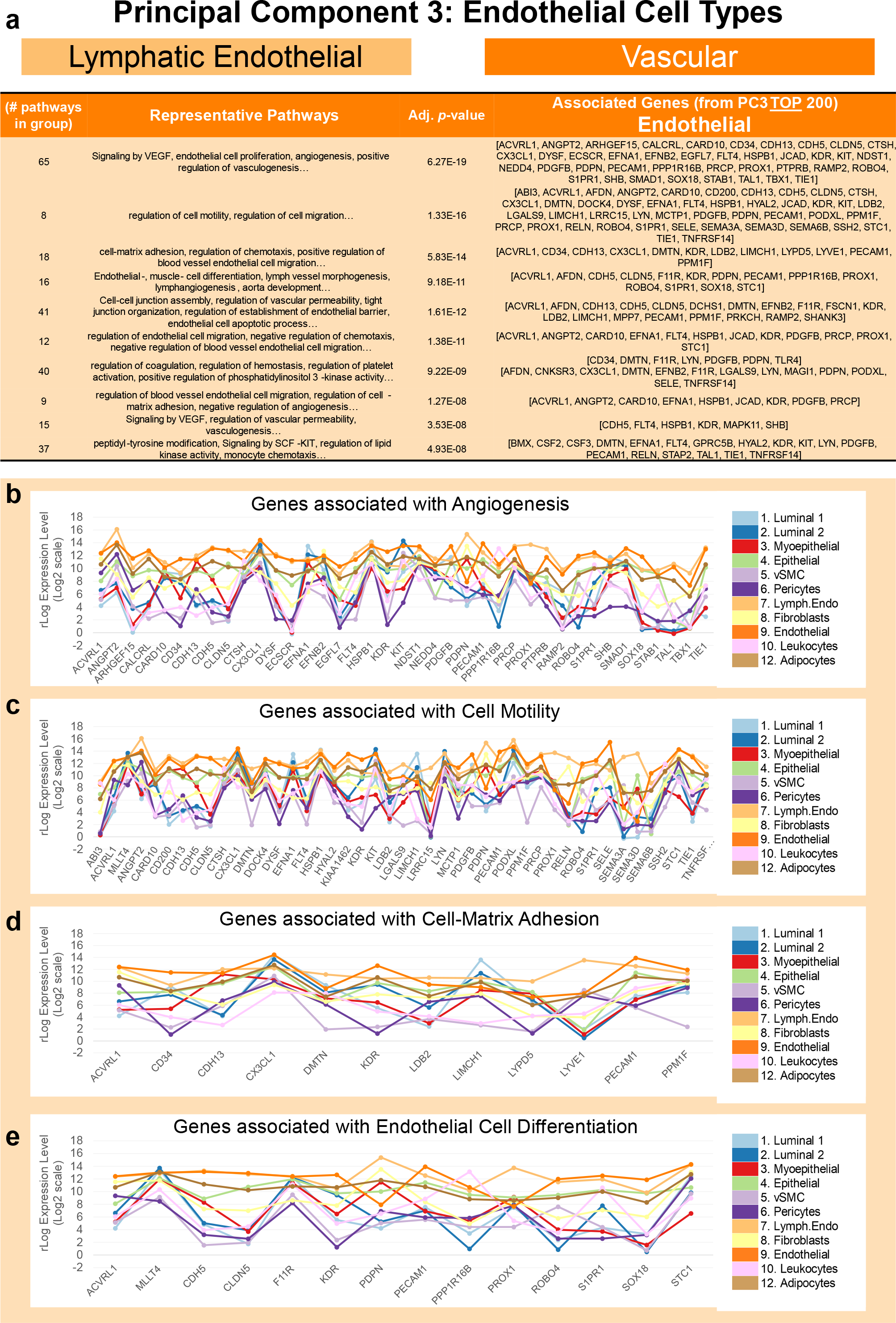
Pathway analysis of Endothelial genes (PC3). (a) Grouped GO/Reactome terms associated with the 200 genes contributing to the positive end of PC3 (top of list), ordered by adjusted p-value. Median mRNA expression (rlog, log2 scale) for PC3 genes associated with select pathways including **(b)** *angiogenesis*, **(c)** *cell motility*, **(d)** c*ell-matrix adhesion*, and **(e)** *endothelial cell differentiation*.

Outside of the anticipated pathways, enrichment analysis of the endothelial genes also uncovered: *Signaling by SCF-KIT* (stem cell factor/c-kit)*, adherens junction organization, regulation of peptidyl-tyrosine phosphorylation* (JAK/STAT)*, semaphorin receptor binding, interleukin-6 secretion* (and IL-10, IL-12 cytokines)*, aminoglycan biosynthetic process, hyaluronic acid binding,* along with roughly 500 other enriched processes and pathways (*p-* values ranging between 1.64E-06 -2.08E-02, Figure 3—table supplement 4: PC3 Endothelial tab). Enrichment of ‘*Signaling by SCF-KIT’* pathway was driven by elevated endothelial expression of the kit tyrosine kinase receptor (KIT), the GRB2-Related Adapter Protein (GRAP), and LYN Proto-Oncogene (LYN). Notably, Kit’s ligand, stem cell factor (KITLG), was differentially expressed between endothelial cell types, as vascular endothelial cells expressed it at a much higher level (40-fold higher) than lymphatic endothelial cells (Figure 12—figure supplement 1).

Endothelial semaphorin expression was intriguing. Semaphorins are a large family of signaling ligands that exist as either secreted or membrane-associated proteins which provide movement and guidance cues^22^. The *Semaphorin receptor binding* pathway was enriched predominantly because of endothelial expression of semaphorin 3a, 3d and 6b (SEMA3A, SEMA3D, SEMA6B, *p-*value=1.47x10^-3^; (Figure 12—figure supplement 2, Figure 3—table supplement 4: PC3 Endothelial tab). The class 3 semaphorins are the only semaphorins in humans that are secreted, and we found that both SEMA3A and SEMA3D were expressed rather exclusively by lymphatic endothelial cells (Figure 12—figure supplement 2a). Interestingly, the co-receptor for sema3 proteins, neuropilin1 and 2 (NRP1 and NRP2), which bind VEGF family members, were highly expressed by the endothelial cell types (*p-*values of 3.13x10^-27^ and 1.23x10^-53^, respectively), with vascular endothelial cells (and pericytes) differentially expressing the highest levels of NRP1. In contrast, the highest levels of NRP2 occurred in lymphatic endothelial cells, which was 2-fold higher than in their vascular counterparts (Figure 12—figure supplement 2b,c). Others have implicated NRP2 as a critical factor in lymphangiogenesis^23, 24^, and our results are consistent with it fulfilling such a role for breast lymphatic cells.

The above analysis has identified many features shared between lymphatic and vascular endothelial cells. Later, we will directly compare these two endothelial cell types and explore their individual traits.

#### Perivascular sub-lineage (vSMC and Pericytes)

After analyzing the endothelial cells, we were left with two other cell groups to explore that were resolved by PC3 and PC4. Perivascular cells were the first of these (Figure 8a). This mesenchymal cell subclass includes pericytes and vascular smooth muscle cells (vSMCs), and the group projected toward the negative end of the third principal component, opposite to the endothelial cells. Like fibroblasts and adipocytes, perivascular cells are mesenchymal in nature; but perform critical functions to the blood vessels they envelop. PC3’s loading factors (at the perivascular end) included genes with known pericyte and vSMC significance (Figure 8c), such as smooth muscle actin (ACTA2, Figure 9g), smooth muscle calponin (CNN1), glutamyl aminopeptidase (ENPEP), regulator of G-protein signaling 5 (RGS5), and platelet-derived growth factor receptor beta (PDGFRB). At the top of the list was the atypical cadherin FAT1, along with phosphodiesterase 3A (PDE3A), peptidase inhibitor 15 (PI15), and carboxypeptidase E (CPE, Figure 8c). Both perivascular cell types differentially expressed AXL receptor tyrosine kinase, which was sixth on the list and, in addition to its other physiological functions, serves as a receptor/co-receptor for many viruses, including Ebola, Lassa, Marburg^25^, SARS-CoV-2^26^, and Zika viruses^27^ –which may help explain the vessel involvement often observed in infected individuals. The perivascular cells also notably and differentially expressed members of the Notch and Hedgehog signaling pathways, such as NOTCH3, HEYL, GLI2, and GLI3, suggesting these cell types fulfill fundamental roles during tissue morphogenesis and maintenance. As before, we used the top 200 genes from the defining principal component for pathway enrichment analysis.

Hypergeometric analysis of the perivascular genes identified 456 enriched processes and pathways from the Gene Ontology and Reactome databases (*p-*values< 0.05, Figure 3—table supplement 4: PC3 Perivascular tab). These aggregated into 24 distinct groups (Figure 13a). Many groups reflected vascular-related biology in line with known perivascular cell functions. Among these were *regulation of blood vessel diameter, regulation of systemic arterial blood pressure by hormone,* and *smooth muscle contraction.* Enriched pathways from other significant groups included: *Epithelial tube morphogenesis, blood vessel- and vascular-development, ECM proteoglycans, artery-* and *cardiac tissue-development, cell fate commitment, syndecan interactions,* and *mesenchymal cell differentiation* (Figure 13 b-e). These analyses identify characteristic processes of both perivascular cell types.

**Figure 13.**
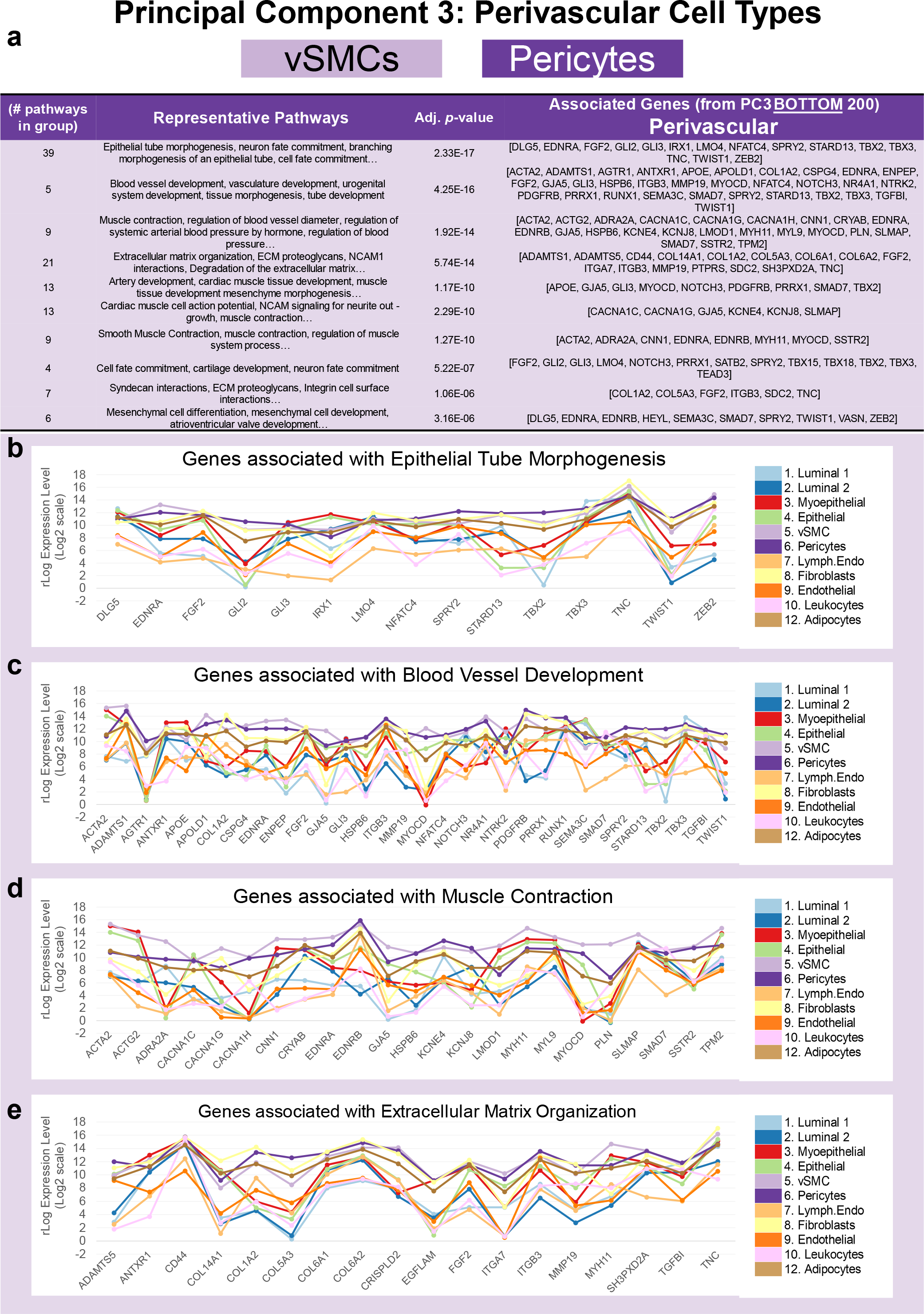
Pathway analysis of perivascular genes (PC3). (**a)** Grouped GO/Reactome terms associated with the 200 genes contributing to the negative end of PC3 (bottom of the list), ordered by adjusted p-value. Median mRNA expression (rlog, log2 scale) for PC3 genes associated with select pathways, including **(b)** *epithelial tube morphogenesis*, **(c)** *blood vessel development*, **(d)** *muscle contraction*, and **(e)** *extracellular matrix organization*.

#### ***Myoepithelial sub-lineage*** *(Myoepithelial Cells and Pop4 Epithelial Cells)*

The final cell group separated by PC3 and PC4 was an epithelial subclass comprised of myoepithelial cells and related Pop4 epithelial cells (a rare cell type we identified during FACS panel development^8^). The fourth principal component resolved these cells, and when we investigated PC4’s loading factors, we found that the most significant gene was none other than keratin 5 (KRT5). Finding keratin 5 was reassuring because it has a long and storied history as a marker of basal epithelial cells^28–30^. Another top gene was alpha-6 integrin (ITGA6/CD49f), which we anticipated because CD49f formed the backbone of our FACS panel –and myoepithelial cells consistently exhibited the highest CD49f staining intensities (Figure 1a, Figure 9f,h). Among the many genes contributing significantly to PC4 (Figure 8d), included the well-known myoepithelial markers: keratins 14 (KRT14, Figure 8d) and 17 (KRT17), laminin subunit alpha-3 (LAMA3), the alpha-1 chain of collagen type XVII (COL17A1), smooth muscle actin (ACTA2, Figure 9g), p63 (TP63, Figure 9i), E-cadherin (CDH1, Figure 9c,h), P-cadherin (CDH3), the transcriptional regulatory factor TRIM29, and integrin subunit beta 4 (ITGB4, Figure 8d, Figure 3—table supplement 3).

Many of these myoepithelial genes notably contribute to the molecular cluster defining the basal-like breast cancer subtype—which provided the rationale for its ‘basal-like’ designation^31^.

As with the preceding cell lineages, we extracted the top 200 loading factors from the defining end of the principal component (PC4, in this case) and used the gene list to explore potentially enriched pathways and processes. Pathway enrichment identified 333 gene sets, which aggregated into 24 groups of related biological functions (Figure 3—table supplement 4: PC4 Myoepithelial (MEP) tab). Processes related to the basement membrane, extracellular matrix organization, and laminin interactions led the way, forming the most significantly enriched group (*p*-value =3.46x10^-16^, Figure 14a). These gene sets were dominated by integrins (ITGA3, ITGA6, ITGB4, ITGB6, ITGB8) and laminin chains (LAMA3, LAMB3, LAMC2), which were highly expressed by myoepithelial cells (Figure 14b-c). Notably, these three laminin genes encode the a3, b3, and g2 chains that form the heterotrimeric laminin 332 molecule—an essential epithelial basement-membrane-specific variant that reinforces the cellular adhesion to the basal lamina^32^.

**Figure 14.**
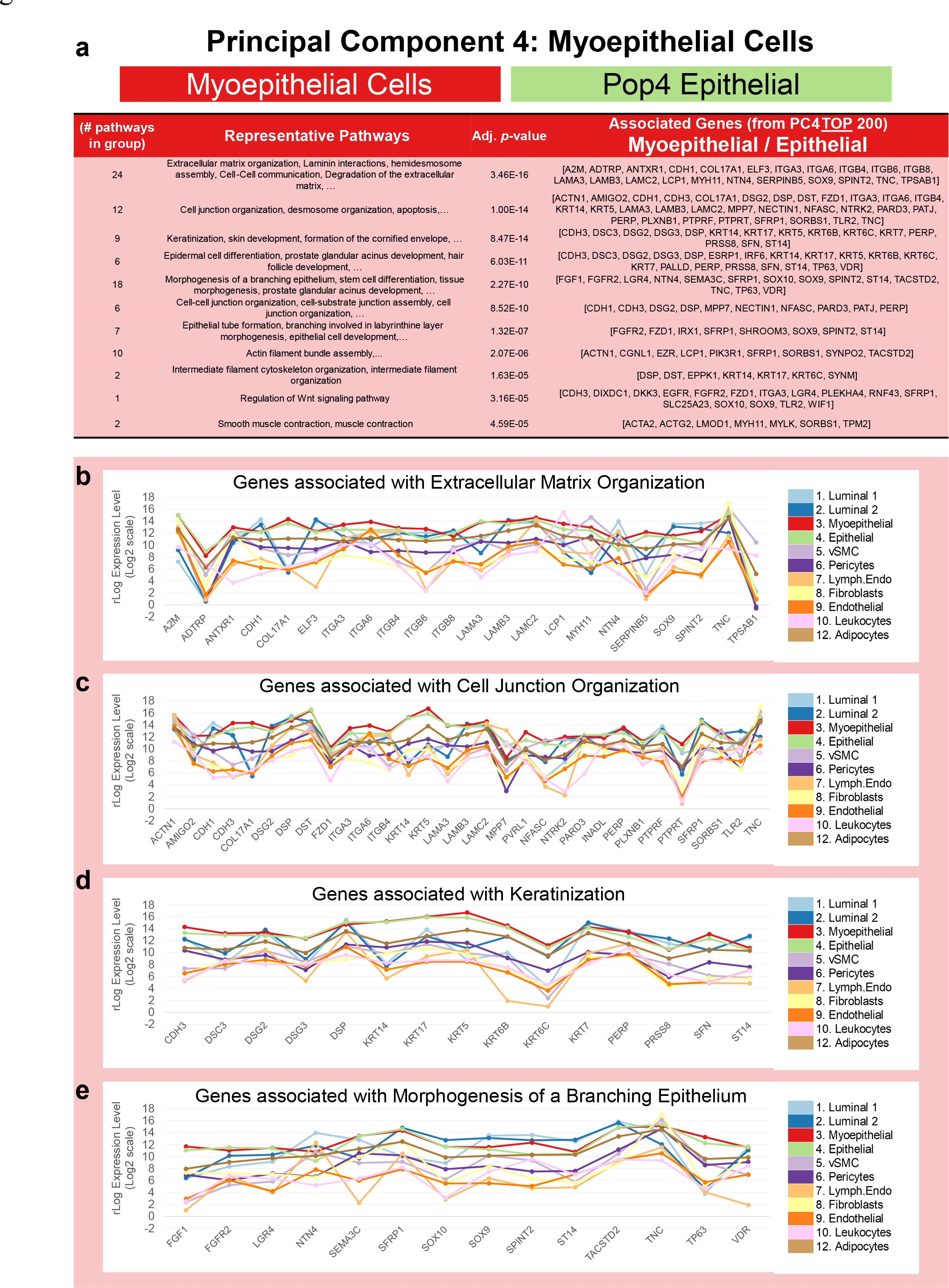
Pathway analysis of myoepithelial-related genes (PC4). (a) Grouped GO/Reactome terms associated with the 200 genes contributing to the positive end of PC4 (top of the list), ordered by adjusted p-value. Median mRNA expression (rlog, log2 scale) for PC4 genes associated with select pathways, including **(b)** *extracellular matrix organization*, **(c)** *cell junction organization*, **(d)** *keratinization*, and **(e)** *morphogenesis of branching epithelium*.

Representative pathways from the other significantly enriched groups included: *cell junction organization, keratinization and skin development, prostate glandular acinus development, morphogenesis of a branching epithelium, epithelial tube formation, actin filament bundle assembly, smooth muscle contraction, intermediate filament organization, regulation of Wnt signaling pathway, and others.* (Figure 14a-e, Figure 3—table supplement 4: PC4 Myoepithelial tab). Among the many integrin, laminin, keratin, and collagen genes represented in these enriched pathways were cadherins CDH1 and CDH3, desmosome-related desmocollin 3 (DSC3) and desmogleins DSG2 and DSG3, stratafin (SFN, 14-3-3 sigma); wnt-modulators ’secreted frizzled-related protein 1’ (SFPR1) and ‘Dickkopf wnt signaling pathway inhibitor 3’ (DKK3); epidermal growth factor receptor (EGFR), fibroblast growth factor 1 (FGF1), and others (Figure 3—table supplement 4: PC4 Myoepithelial tab).

Myoepithelial cells have historically been the most studied breast cell type, likely because of their prevalence in the tissue and ease of culture. Our results are consistent with prior knowledge of myoepithelial function: that these cells are basally located, help shape the basement membrane to which they attach via hemidesmosomes and are rich with smooth muscle actin and myosin that provide them with contractile ability. Furthermore, the mammary gland is formed through invagination of the skin, and the myoepithelial gene list reflects this connection, as it contains genes associated with epidermal development. This list includes basal keratins (KRT-14, -17, -5, -6b, -6c, and -7), the laminin 332 chains, desmosomal genes, vitamin D receptor (VDR), and transcription factors SOX9 and p63. These attributes distinguish myoepithelial cells from all others in the breast. Our results pinpoint the specific factors produced by myoepithelial cells, affirming their pivotal role in ECM-mediated cellular functions.

This section highlights the broad characteristics of cell lineages composing the breast. These characteristics emerged from pathway enrichment analyses using gene lists defined by the principal components aligning and separating each cell lineage. Hundreds of pathways were revealed. Some we anticipated, whereas many others were not. Although we have here highlighted the most statistically significant pathways emerging from this analysis (Figures 10-14), many more exist and are listed in the supplementary tables (Figure 3—table supplement 4). A full review of these results will undoubtedly provide a more comprehensive perspective.

### Gene families contributing to cellular identity

Above, we explored the distinguishing qualities of different cell lineages by determining the pathways enriched in PCA-defined gene lists. These analyses relied on curated gene sets from the Gene Ontology (GO) and Reactome databases—but not every human gene is represented in these archives. For example, the Reactome knowledgebase currently contains 11,291 human proteins (including known isoforms) that, by our count, are encoded by just over 9,300 genes. Because we measured over 20,000 genes in each cell type, many are not captured in the above analysis. Therefore, we have complemented the prior results by shifting to a more ‘gene-centric’ approach. This method entailed identifying all differentially expressed (DE) genes in the breast cell dataset (DESeq2^13^, adj. *p*-value ≤ 0.1, Figure 3—table supplement 5). We then used this information to identify gene families with the most variably expressed genes. Due to their differential expression, these gene types must contribute to cell-specific processes and the unique features of each cell type. Their identification and patterns of expression provided countless insights.

The HUGO Gene Nomenclature Committee has organized 25,320 human genes into 1,519 manually curated gene families (HGNC gene group index^33, 34^). When we explored the breast cells’ transcriptomes, we identified 1,421 gene families in our dataset (Figure 3—table supplement 6). These included *vacuolar-type ATPases*, *matrix metallopeptidases*, *mucins*, *collagens*, *glutathione transferases*, *erb-B2 tyrosine kinases*, *histones*, *interleukins*, and more (Figure 3— Table supplement 6). The sizes of the families varied widely, from 1 to 831 genes (x̄ = 14, sd = 42.2) with a median family size of six genes per family (Fig.15—figure supplement 1).

**Figure 15.**
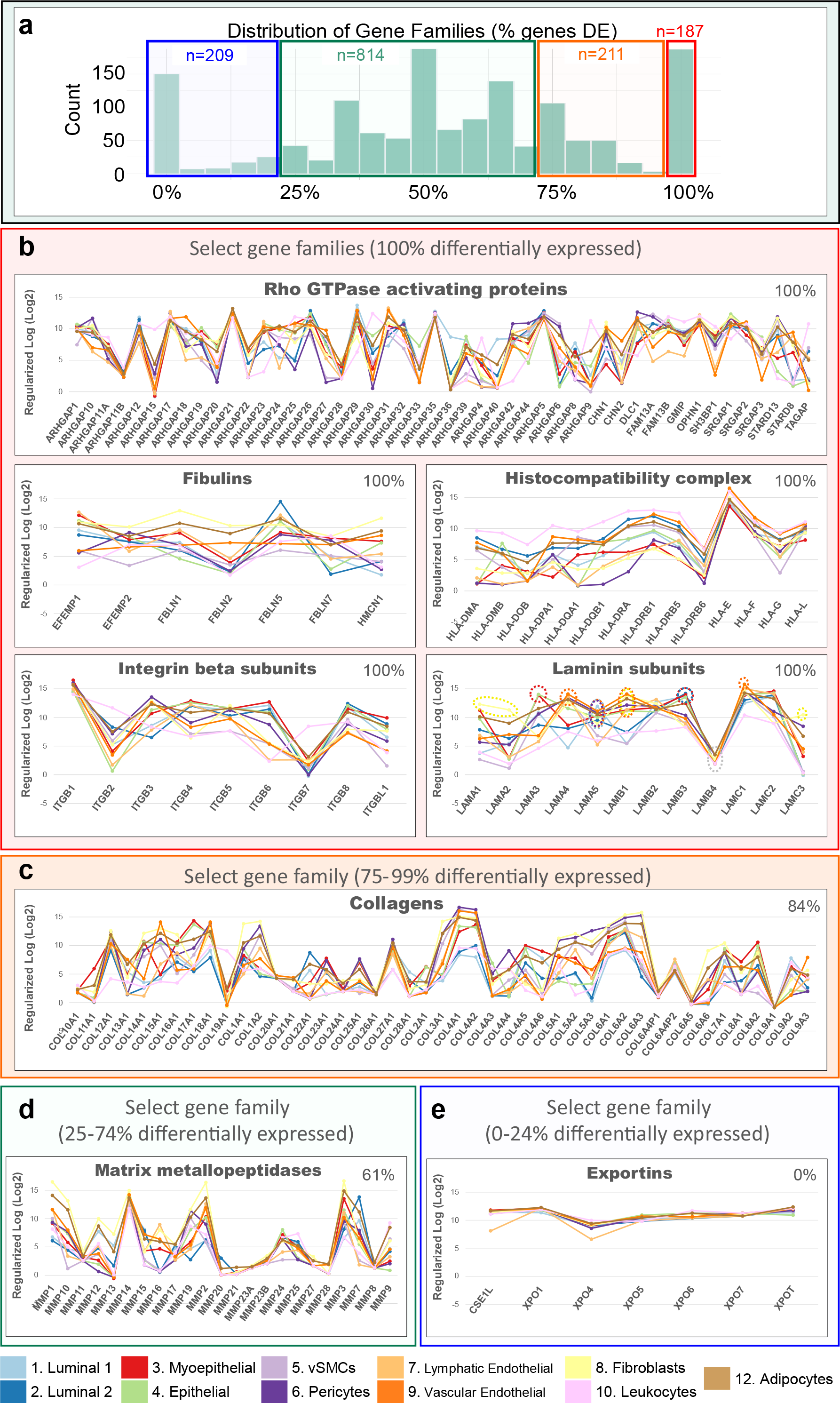
Gene types contributing to cell-specific features. Of the 1,519 types of genes (gene families) defined by HGNC, we found 1,421 gene families (of all sizes) expressed within the breast cells transcriptomes. **(a)** Distribution of the proportions of differentially expressed (DE) genes in each gene family (BH Adj. *p-*value ≤ 0.1). The proportion of DE genes in these families ranged between 0-100%, which we grouped into different bins: 0-24% (blue, n=209), 25%-74% (green, n=814), 75% to 99% (orange, n=211), and 100% (red, n=187). **(b-e)** Median transcript levels (rlog, log2 scale) across all cell types of genes composing select families with **(b)** 100% DE genes, **(c)** 75%-99% DE genes, **(d)** 25-74% DE genes, and **(e)** 0-24% DE genes.

To determine the families with the most variably expressed genes, we examined the expression of each gene, from each family, within every breast cell type—and focused on families that had six or more genes (≥ median size). We discovered 187 gene families composed entirely of differentially expressed genes (Figure 15a). Twenty of these families remained once we excluded those containing fewer than six genes. These included *Rho GTPase activating proteins*, *fibulins*, *histocompatibility complex genes*, *laminin subunits, integrin beta subunits, and more* (Figure 15b, Figure 3—Table supplement 6). The cellular processes affected by genes in these twenty families are wide-ranging, as they are involved in Rho signaling, cAMP regulation, interactions with the immune system, and cellular adhesion—between other cells, the extracellular matrix, and basement membranes. Below, we highlight nine families that we believe are most intriguing, but we have also provided a full accounting of each gene family in a table supplement— indicating the number and percentage of genes differentially expressed among them (Figure 3— Table supplement 6)

#### Laminins

Each gene family told a story. The family of laminins, for example, is composed of 12 genes that code for laminin a, b, and g chains. These chains aggregate to form an array of heterotrimeric laminin molecules with distinct properties that polymerize in the extracellular matrix and help shape basement membranes. Above, we found that myoepithelial cells highly expressed the chains composing laminin 332 (Figure 14). Inspection of the entire laminin family revealed that fibroblasts highly express four laminins (LAMA1, LAMA2, LAMB1, and LAMC3); endothelial cells express the highest levels of three (LAMA4, LAMB1, and LAMC1); whereas ER^Pos^ luminal, vascular endothelial, and pericytes significantly expressed LAMA5 (Figure 15b-laminin subunits). There were also several intriguing instances where closely related cell types expressed widely disparate levels of a particular laminin chain. Notable examples were the differential expression of LAMA1 between ER^Pos^ and ER^Neg^ luminal cells, LAMB1 between the two endothelial cell types, and LAMC3 between pericytes and vSMCs.

Furthermore, no cell type meaningfully expressed LAMB4, consistent with its consideration as a pseudogene^35^.

The relationship between cells and their basement membranes is dynamic and reciprocal. Each cell type contributes to the basal lamina’s structural diversity by secreting basement membrane components like the laminins and other extracellular molecules. These integral basement membrane proteins anchor the cells via membrane-tethered receptors and transmit signals to the cells, where they become integrated and influence multiple cellular processes.

#### Fibulins

One such related and intriguing ECM-related gene family was the fibulins. Fibulins are RGD-containing extracellular matrix glycoproteins that bind basement membrane proteins, such as tropoelastin, fibrillin, and fibronectin^36^. Whereas we found that fibroblasts highly expressed most fibulin family members, fibulin 5 (FBLN5) levels were highest in the ER^Neg^ luminal epithelial cells (Figure 15b-fibulins). FBLN5 was also notably expressed by lymphatic endothelial cells at substantially higher levels than their vascular counterparts. Fibulin 5 is one of the smallest fibulins, is secreted, has unique calcium and integrin binding properties, and is essential for elastic fiber formation^36^—and thus likely contributes to the elasticity of the lymphatic endothelium. We also found fibulin 3 (EFEMP1) was also differentially expressed by lymphatic endothelial cells, presumably providing similar contributions.

The fibulins and laminins are just two examples of the 187 gene families whose members were entirely differentially expressed. Of the remaining families (having fewer than 100% DE genes), we identified 211 containing between 75 to 99% DE genes (Figure 15a). Over half of these families were composed of six or more genes. These 126 families included *integrin alpha subunits*, *toll-like receptors, kruppel like factors, adenylate cyclases, low-density lipoprotein receptors, glypicans, interleukin receptors, ephrins,* and others (Figure 3—Table supplement 6). One of the larger and more familiar families was the *collagens*.

#### Collagens

The family of collagens consists of 45 individual genes, and, as expected, we found mesenchymal and myoepithelial cell types highly expressed most of them. There were some notable exceptions, however. For example, the vascular endothelial cells highly expressed COL15A1 and COL9A3, whereas COL22A1 was expressed uniquely by luminal epithelial cells and adipocytes (Figure 15c). Particularly enlightening were the cell types expressing the collagen IV genes.

Collagen IV is a sheet forming collagen and the principal constituent of basement membranes. It provides cellular integrin and non-integrin attachment sites through its triple-helical and non-collagenous domains^37^. There are six collagen IV isoforms, which assemble to form one of three triple-helical heterotrimers, α1α1α2, α3α4α5, or α5α5α6^37, 38^. We found the expression of collagen IV chains in the breast cells quite revealing, as the most robust levels occurred in cells associated with the basal lamina, but patterns of isoform expression differed by cell type. For example, pericytes and vSMCs, which are embedded within, and help maintain the vascular basement membrane, expressed the highest levels of COL4A1, COL4A2, and COL4A4 (Figure 15c, Figure 15—figure supplement 2a-c). In contrast, myoepithelial cells, which contribute significantly to epithelial basement membranes, exhibited the highest levels of COL4A5 and COL4A6. From these results, we would predict that the dominant collagen IV heterotrimer in epithelial basement membranes to be α5α5α6— whereas our data suggests it would be α1α1α2 within the basal lamina of the vasculature.

#### Integrins

The family of integrins was highly differentially expressed across the breast cell types. Integrins connect the extracellular matrix (ECM) to the cytoskeleton and act as receptors that integrate extracellular signals, relaying information to the cell’s interior. Each integrin heterodimer forms from one α and one β subunit, from families of 18 α and 8 β isoforms, generating the 24 observed αβ combinations. When we investigated the family of integrins in the breast cell types, we found they were all differentially expressed—except for ITGAD, which was essentially absent in every cell type (Figure 15b-integrin beta subunits, Figure 15—figure supplement 2d). Due to their diversity and cell-surface expression, sixteen integrins serve as ‘cluster of differentiation’ (CD) antigens. Above, we introduced that a critical marker in our FACS was CD49f, which is integrin α6 (ITGA6) —and that myoepithelial and endothelial cell types highly expressed it (Figure 4—figure supplement 2a, Figure 14b). Integrin a6 binds laminins through heterodimers formed with either b1 or b4 subunits. The a6b1 and a6b4 integrins, along with a3b1 and a7b1, are the four main laminin-binding integrins. Thus, along with α6 integrin, the main laminin-binding integrin heterodimers use a3 and a7 integrins.

Investigating these transcripts’ expression in breast cells, we found that myoepithelial cells and ER^Pos^ luminal epithelial cells expressed the highest levels of ITGA3. In contrast, pericytes and vSMCs expressed the highest levels of ITGA7, and fairly specifically, which is consistent with prior reports of this subunit’s expression in vascular smooth muscle cells^39^ (Fig.15—figure supplement 2d-e). These data suggest that myoepithelial and perivascular cells use distinct integrin heterodimers to bind laminins within their respective basement membranes.

Another CD antigen is ITGAM (CD11b), which is frequently used to identify myeloid cells. We indeed found that within the breast cell populations, ITGAM was highly expressed by leukocytes—although it was also strongly expressed by adipocytes and luminal epithelial cells (Fig.15—figure supplement 2d). Moreover, leukocytes also highly expressed six other integrins: ITGAE, ITGAL, ITGAX, ITGA4, ITGB7, and ITGB2 (Fig.15—figure supplement 2d). ITGB7 and ITGB2 were exclusively expressed by leukocytes, consistent with their recognized role in trafficking leukocytes to sites of injury and infection^40^. Our data reflect this cell-type specificity, and transcript levels within the different breast cell types provide a detailed accounting of these and other expressed integrin isoforms, offering valuable insights into potential dimer combinations used by the different cell types.

Along with the collagens, alpha integrins, and 206 additional gene families, families containing between 75 to 99% DE genes included the *plakins* (88% genes DE), *spectrins* (86%), and *receptor tyrosine kinases* (80%). By comparison, we identified 209 families whose members were expressed relatively equally across all breast cell types (0-24%), such as the *exportins* (0%, Figure 15e). Furthermore, we also investigated families formed by a more moderate makeup of DE genes, i.e., between 25 to 74% (Figure 3—table supplement 6). Among these 814 families were the *semaphorins* (70%), *proteoglycans* (71%), *mucins* (71%), *prostaglandin receptors* (67%), *nuclear hormone receptors* (66%), and *claudins* (64%, Figure 3—table supplement 6).

Also included were the *matrix metalloproteinases* (61% DE), which are a family of proteinases that degrade all extracellular matrix components—and thus help regulate the homeostatic balance of ECM molecules in the breast microenvironment (Figure 15d).

#### Matrix Metalloproteinases (MMPs)

Of the 23 MMP isoforms encoded within the human genome, we identified 14 differentially expressed among the breast cell types. Fibroblasts highly expressed over half of these MMPs, which included gelatinase A (MMP2), stromelysin 1 and 2 (MMP3, MMP10), macrophage elastase (MMP12), membrane type 1,3, and 4 (MMP14, MMP16, MMP17), and MMP19. Predominant substrates for these MMPs include fibronectin, laminins, type III, IV, and V collagens, elastin, fibrin, pro-MMP2, pro-gelatinase A, aggrecan, and cartilage oligomeric matrix protein^41^. In addition, we found leukocytes (and adipocytes) expressed gelatinase B (MMP9) at its highest levels, consistent with reports of it being absent in fibroblasts and playing an essential role in ECM proteolysis and leukocyte migration^42, 43^.

Arguably the most notable deviation from the near-universal fibroblast expression of MMPs was the differential expression of matrilysin (MMP7) by ER^Neg^ luminal epithelial cells, where we found levels to be 18-fold higher than in their ER^Pos^ counterparts –and 237-fold higher than in fibroblasts (Figure 15d).

#### Tight Junctions

Essential to multicellular life are the mechanisms in which cells join to form the fabric of tissues and guide their shape and functions. Above, we introduced the family of integrins that serve as central receptors binding cells to the ECM. Different cell lineages have distinct requirements. For example, breast fibroblasts are embedded within a collagen-rich matrix, whereas perivascular cells, which are also mesenchymal in nature, encircle vessels, help generate the endothelial basement membrane, and collaborate with endothelial cells to form the vasculature. The endothelial cells, in turn, must create tight junctions to generate a patent and sealed network for transporting liquid blood—especially important for forming the blood-brain barrier of the central nervous system. Epithelial cells are also unique because their glandular role requires them to develop functional sheets while maintaining a semi-permeable barrier to the luminal space, where they may secrete and collect milk. The lumen is also essentially connected to the outside environment through the ductal network and nipple opening, so epithelial cells must also establish a barrier to protect against bacteria and other pathogens—working in concert with adjacent leukocytes that closely associate with the epithelium. Thus, the formation of tight junctions is essential to the gland.

Tight junctions are the prominent cell-to-cell junctions of epithelia and endothelia, establishing the selectively permeable barrier between neighboring cells^44^. Claudin and occludin proteins are vital components of this cell adhesion complex. Investigating their expression in the breast cells, we found the vast majority of claudins (11 of the 19 expressed) were predominantly limited to the two luminal epithelial cell types, although there were some notable exceptions (Figure 16a,b). One was the specific expression of claudins 5 and 14 (CLDN5, CLDN14) by endothelial cells—and to a lesser extent, by adipocytes. Whereas the vascular and lymphatic endothelial cells both highly expressed CLDN5, CLDN14-expression was limited to the vascular endothelial cells. Another exception was myoepithelial CLDN11 expression. Also, of the eleven claudins expressed by the luminal epithelial cells, five were differentially expressed between the two luminal types, with CLDN9, 22, and 23 transcripts being more highly expressed by the ER^Pos^ cells and CLDN16 and 10 being more highly expressed by the ER^Neg^ cells (Figure 16a,b).

**Figure 16.**
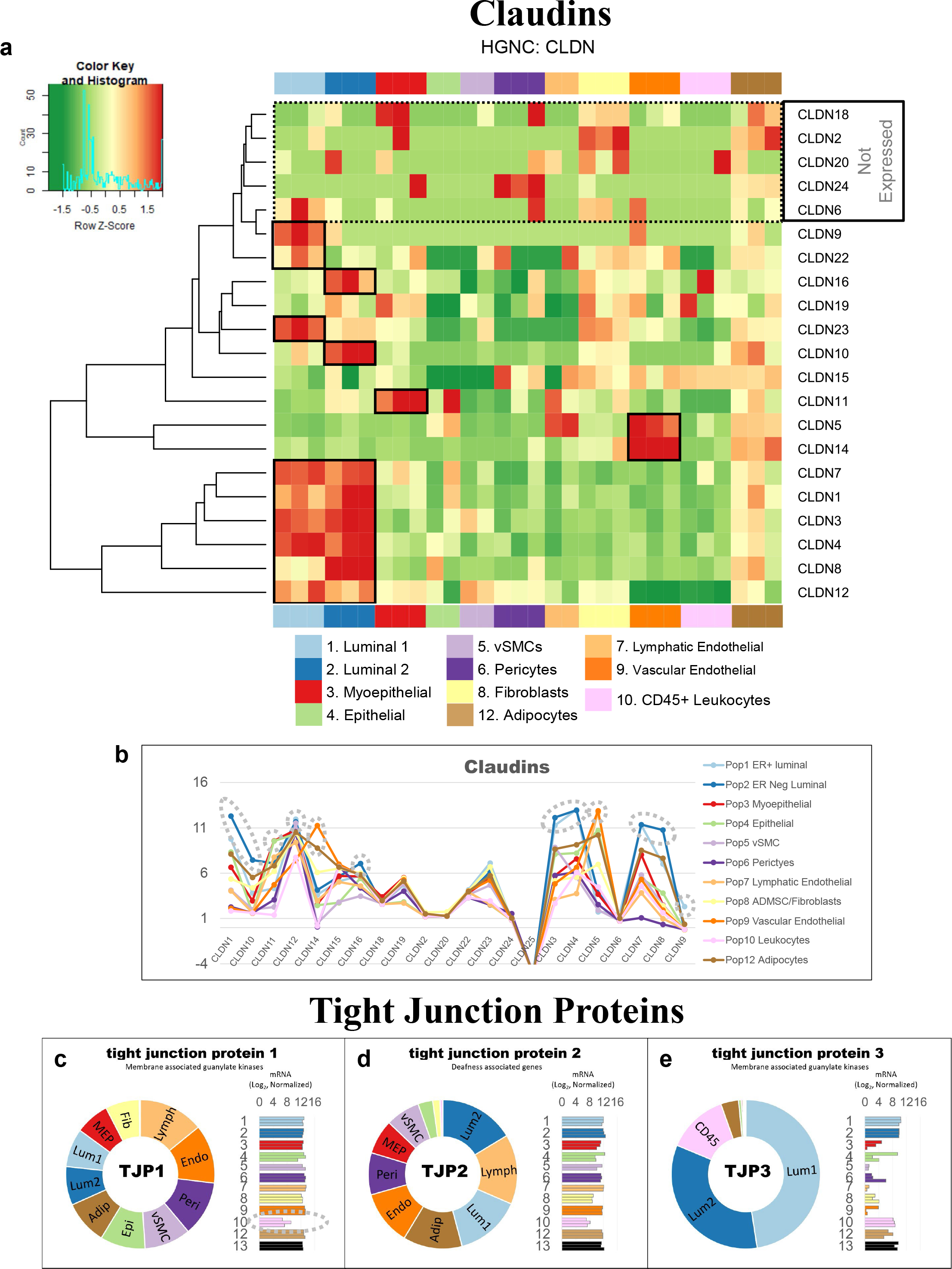
Tight Junctions. (a) Heatmap displaying transcript levels of genes belonging to the claudin gene family (VST, normalized by row). Dashed box indicates claudins that were not appreciably expressed by any cell type. Dark boxes highlight genes differentially expressed by luminal epithelial cells, endothelial cells, and myoepithelial cell types. **(b)** median rlog levels of the claudin family genes. Dashed circles identify genes outlined in the heat map. **(c-e)** Transcript levels for the zona occludens tight junction proteins: **(c)** TJP1, **(d)** TGP2, and **(e)** TJP3 (rlog values on log2 (bar graph) and linear (donut graph) scales).

Furthermore, claudin 1 was one of the six claudins expressed by the luminal epithelial cells, and tissue staining confirmed luminal cell expression, as the staining was confined to the apical surface of the paracellular space, circumscribing the luminal surface (Figure 16—figure supplement 1). These data are consistent with the epithelial and endothelial use of tight junctions, and results pinpoint the differential isoform specificity among the five claudin-expressing cell types.

The other critical components of tight junctions are the occludins (tight junction proteins encoded by TJP1, TJP2, and TJP3 genes). When we explored their expression among the breast cell types, we found that TJP1 and TJP2 (encoding zona occludens -1 and -2) shared a similar pattern, as they were both expressed relatively equally among all cell types. The only exceptions were that for TJP1, leukocyte levels were roughly 50-fold lower than the other cell types, and for TJP2, levels were lower in both fibroblasts and leukocytes. In contrast, TJP3 expression (ZO-3) was predominantly limited to the luminal epithelial cells (Figure16c-e).

#### Gap Junctions

In contrast to the occluding tight junctions that seal the paracellular space between neighboring cells, gap junctions create channels between them, allowing the exchange of ions and low molecular weight metabolites between neighboring cells^45^. We found that the breast cell types expressed eleven of the 20 genes in the gap junction family (Figure 17 a,b).

**Figure 17.**
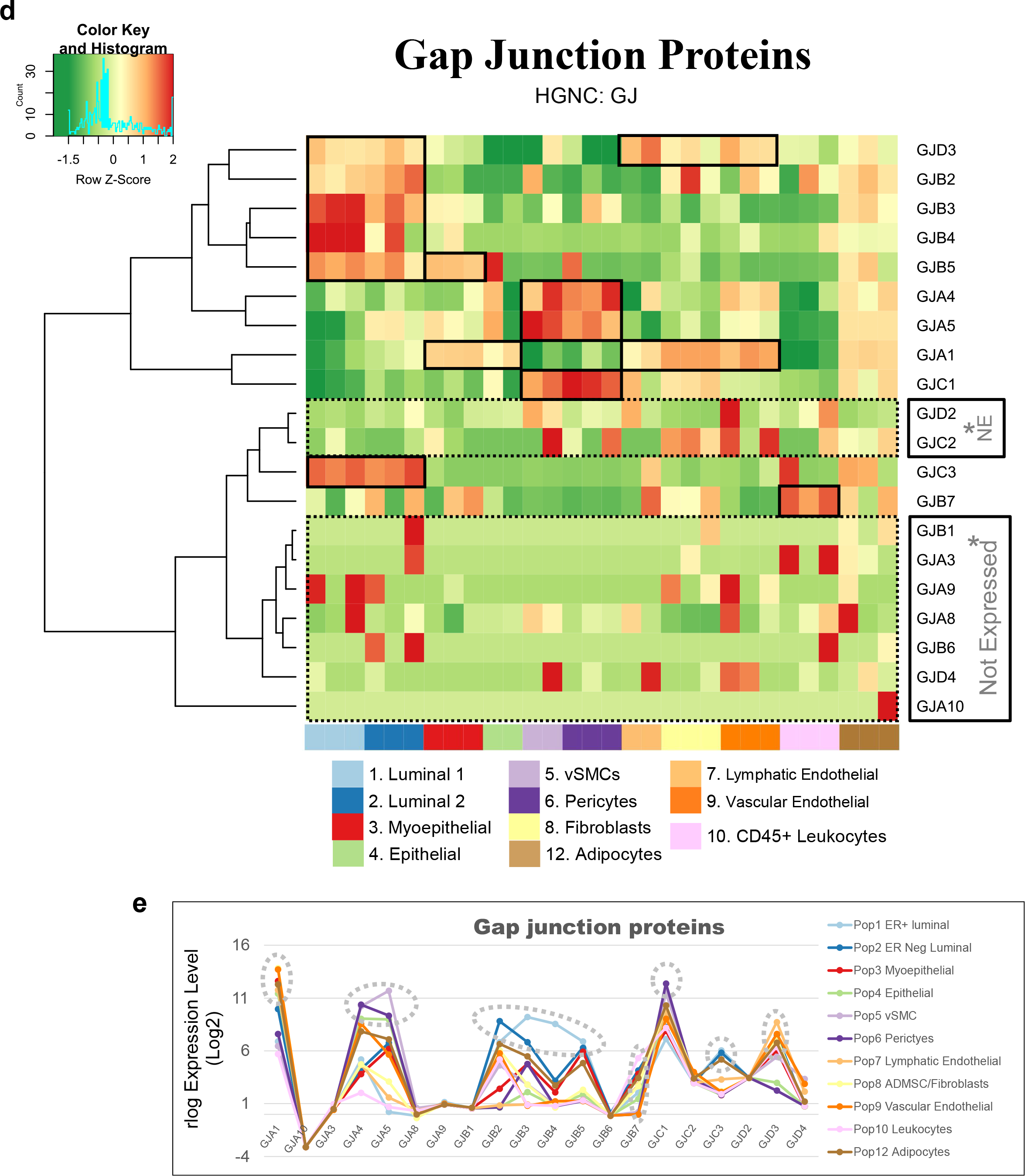
Gap junction proteins. (a) Heatmap displaying transcript levels of genes encoding gap junction proteins (VST, normalized by row). Boxed areas highlight the cell types expressing select gap junction proteins. Dashed boxes identify genes not appreciably expressed by any cell type, as evidenced by the rlog values shown in b. **(b)** Median rlog transcript levels of gap junction genes on a log2 scale; differentially expressed genes are encircled.

Myoepithelial cells expressed two of these genes (GJA1 and GJB5, i.e., connexin 43 and 31.1), whereas luminal epithelial cells expressed six (GJB2, B3, B4, B5, C3, and D3), sharing only with myoepithelial cells the expression of GJB5. Furthermore, ER^Pos^ and ER^Neg^ luminal cells differentially expressed three gap junction genes, with GJB3 and GJB4 being higher in ER^Pos^ cells—and GJB2 being highest in the ER^Neg^ luminal cells. Another intriguing feature was the fibroblast and endothelial expression of GJA1, which they shared with Pop4 epithelial and myoepithelial cells. We also found that leukocytes uniquely expressed the highest levels of GJB7, although we are uncertain what role it would fulfill in these cells. One final observation was the perivascular expression of GJA4, GJA5, and GJC1; and the differential expression of GJA5 by vSMCs, at a 5.5-fold higher level than pericytes (Figure 17 a,b). From these data, we can identify the potential homomeric and heteromeric gap junction complexes used by the perivascular and other cell types.

Whereas gap junctions facilitate the direct connection of cytoplasm between cell types and are essential mediators of cell-to-cell communication, other forms of communication occur through autocrine, paracrine, and endocrine signaling mechanisms. Multicellular organisms—and tissues— rely on efficient and responsive communication between different cells, sometimes separated by great distances. One of the most evolutionary-protected mechanisms facilitating cell communication is the production of proteins, peptides, fatty acids, glycans, cholesterol derivatives, gasses, and other metabolites by one cell, which are detected by another and elicit a response^46^. Unfortunately, the vast number of ligand and receptor families preclude us from exploring each of them here. Instead, we chose to inspect two different receptor families that exemplify the heterogeneous expression patterns observed among breast cell types: *receptor tyrosine kinases* and *nuclear hormone receptors*. The most studied and best-understood family of receptors possessing inherent kinase activity are the receptor tyrosine kinases (RTKs).

#### Receptor Tyrosine Kinases

The human genome encodes 58 different receptor tyrosine kinases (RTK) whose ligands span across twenty subfamilies. These ligands are wide-ranging and include the ephrins, glial-cell derived neurotrophic factor, and: epidermal-, nerve-, hepatocyte-, fibroblast-, vascular endothelial-, and insulin-like growth factors, among others. RTKs are vital regulators of critical cellular processes and are frequently involved in many cancer types. Among the breast cell types, most RTKs displayed a cell-type-specific expression pattern (Figure 18a-m). Notably among these was the robust mesenchymal expression of platelet-derived growth factors alpha and beta (PDGFRA, PDGFRB), fibroblast growth factor receptor 1 (FGFR1), AXL tyrosine kinase receptor (AXL), and orphan receptor 2 (ROR2, Figure 18a-m). In contrast, most cells equally expressed insulin receptor (INSR), insulin-like growth factor 1 receptor (IGF1R), receptor-like tyrosine kinase (RYK), and lemur tyrosine kinase 2 (LMTK2). As expected, the endothelial cells differentially expressed angiopoietin receptors tie2 and tie1 (TEK and TIE1) and VEGF receptors 1 and 3 (FLT1 and FLT4), which are known to regulate many facets of vascular biology (Figure 18a,i). We have previously described, consistent with prior reports^47–49^, how stem cell growth factor receptor (KIT) was highly expressed by ER^Neg^ luminal cells^8^, but we also found it to be appreciably expressed by vascular endothelial cells and vSMCs (Figure 18a, Figure 12—figure supplement 1). Her2 (ERBB2) is amplified in a subset of breast cancers and has been successfully targeted for therapeutic intervention (Herceptin/trastuzumab). We found that Her2, along with two other members of the ErbB family (ERBB3 and ERBB4), were highly expressed by luminal epithelial cells. In contrast, the remaining family member, epidermal growth factor receptor (EGFR), was expressed across a broader range of cell types, with fibroblasts and myoepithelial cells displaying the highest transcript levels (Figure 18j-m).

**Figure 18.**
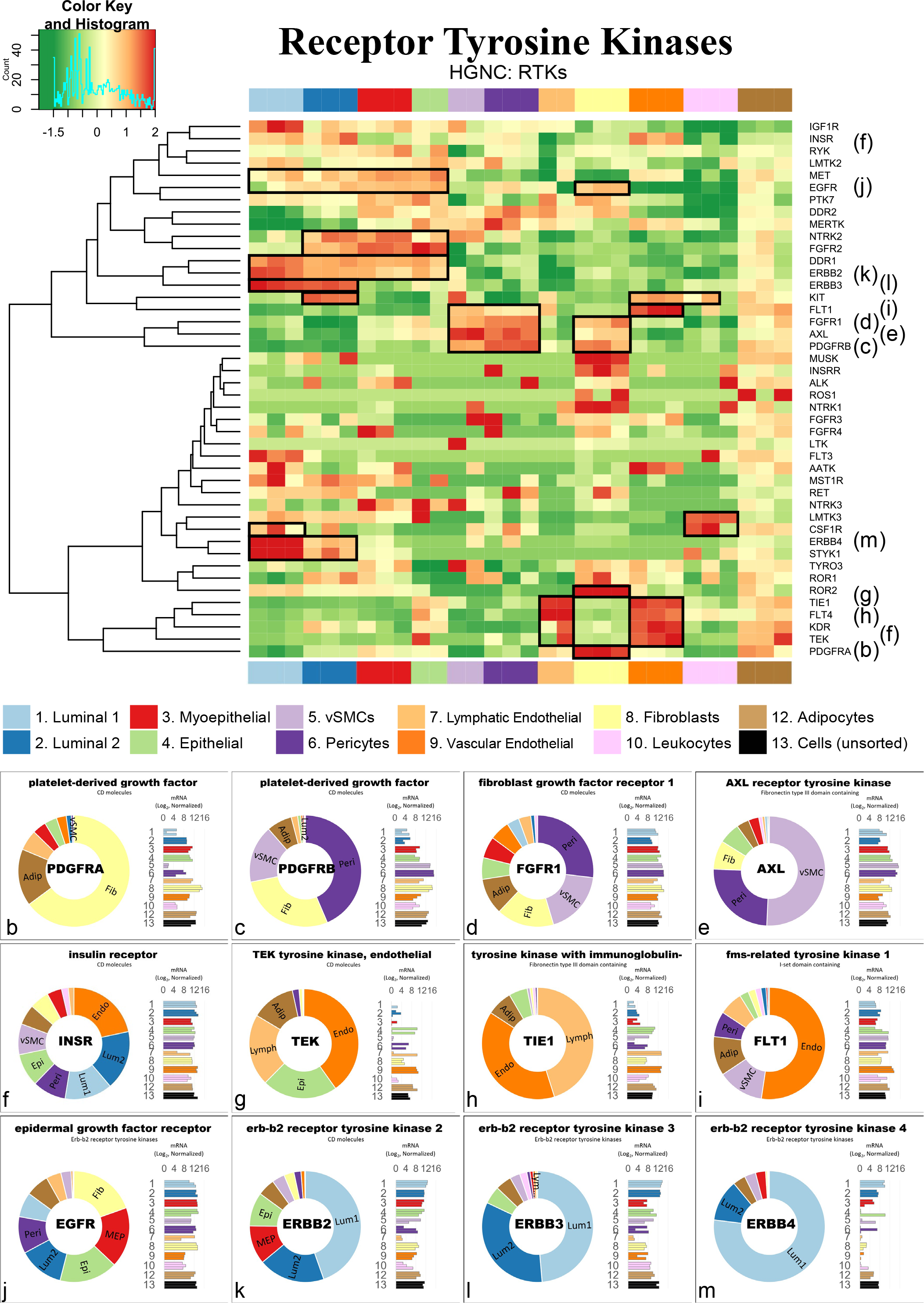
Receptor tyrosine kinases. (a) Heatmap displaying transcript levels of genes belonging to the tyrosine kinase receptor gene family (VST, normalize by row). Boxed areas indicate cell types expressing highest levels of select kinases. **(b-m)** Transcript levels of the select RTKs (rlog values on log2 (bar graph) and linear (donut graph) scales).

Overall, RTKs display a complex pattern of expression. Most, 80%, were expressed differentially across breast cell types, sometimes even within the same lineage, e.g., PDGFRB, KIT, ERBB3.

#### Nuclear Hormone Receptors

Nuclear hormone receptors were another receptor family demonstrating high cell type specificity. These receptors act as ligand-induced transcription factors and are activated upon binding to small lipophilic molecules, such as thyroid or steroid hormones, vitamin D3, retinoic acids (vitamin A), ecdysone, oxysterols, bile acids, fatty acids, leukotrienes, and prostaglandins^50^. The human genome has 49 nuclear hormone receptors, and their regulatory effects are wide-ranging. Of this family’s many prominent members, the most notable for the breast field is estrogen receptor (ERα).

Interestingly, much of what we know about nuclear hormone receptors is rooted in the early studies of estrogen and how it regulates female reproductive organs, including the breast^51^. We found 44 nuclear hormone receptors expressed in breast cells—31 differentially. The androgen, progesterone, and estrogen α receptors were all strongly expressed by luminal epithelial cells— more so in ER^Pos^ cells than their ER^Neg^ counterparts (Figure 19a-d). Both thyroid hormone receptors were similarly expressed among the cell types, except perivascular cells, which expressed 5.3-fold lower levels of THRA—and endothelial cells and leukocytes, which expressed 8.5-fold lower levels of THRB (Figure 19e). The cell-type specific expression extended to many other nuclear hormone receptors, such as vitamin D receptor (epithelial cell types), estrogen-related receptor beta (ER^Pos^ luminal cells), hepatocyte nuclear factor 4 gamma (both luminal cell types), estrogen receptor beta (Pop4 epithelial and vascular endothelial cells), and nuclear receptor subfamily members –5A2 (both endothelial cell types), –4A1(mesenchymal cells), –4A2 and –4A3 (perivascular cells, Figure 19f-m). Nuclear hormone receptors possess broad influence over many cell processes. The cell-specific expression patterns detailed here identify the cell types in which these receptors likely play a prominent role.

**Figure 19.**
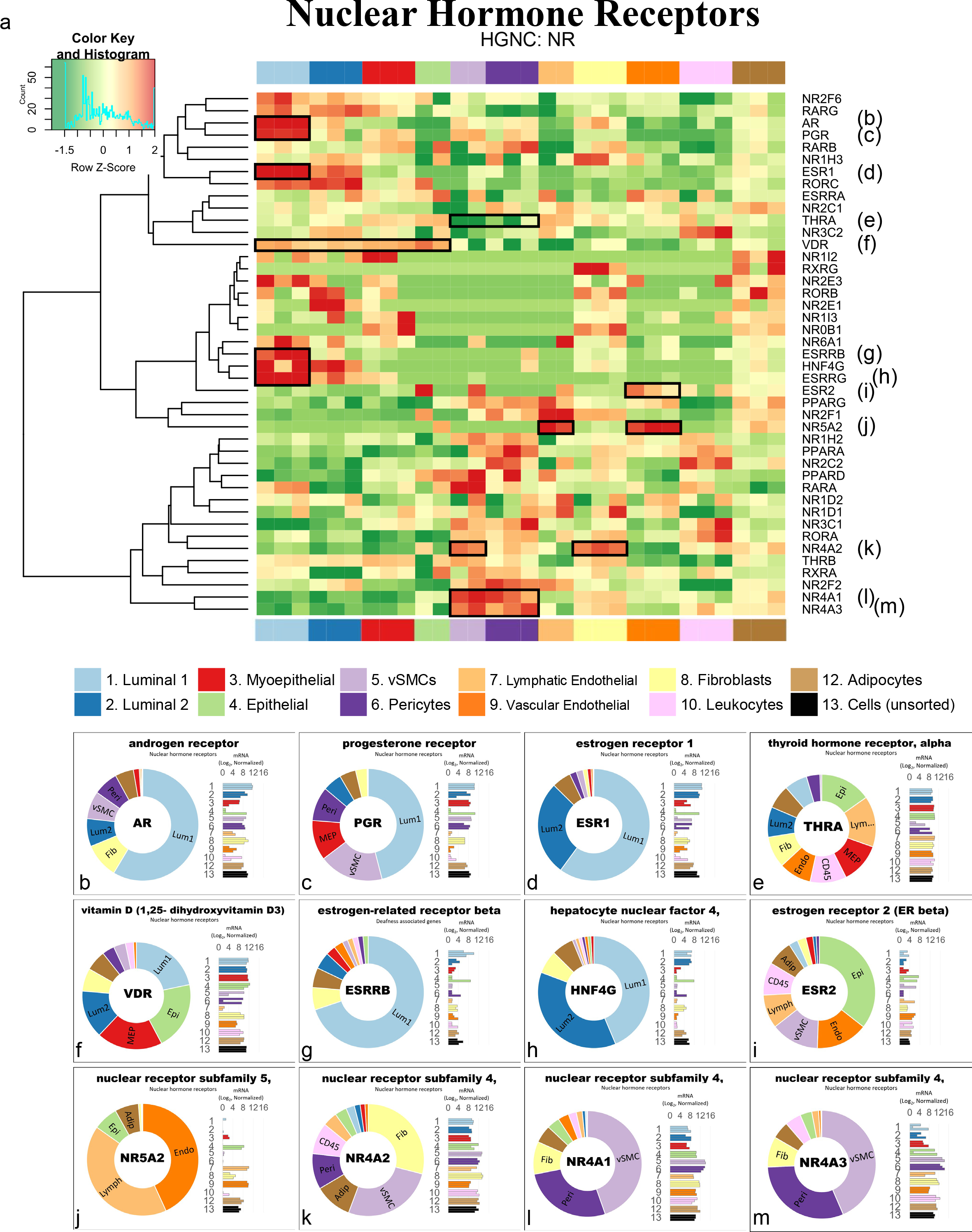
Transcript levels of nuclear hormone receptors. (a) Heatmap displaying transcript levels of genes belonging to the nuclear hormone receptor gene family (VST, normalized by row). Boxed areas indicate high/low gene expression in different cell types, which are highlighted below by the donut plots. Select genes include **(b-m)** mRNA values (rlog) are provided on log2 scale (bar graph of each biological replicate) and linear scale (donut graph of median value) which are both color-coded by cell type.

Analyzing gene families has provided us with an unconventional look at how different types of genes contribute to the cell types in the breast, and the results identified the kinds of genes that vary most across the spectrum of cell types. Above, we meticulously described nine gene families whose functions extend from the nucleus (hormone receptors) to the exterior cell microenvironment (laminins). However, these represent only a limited percentage of families we identified as having a high proportion of highly variable genes. Many more are listed in the supplementary tables (Figure 15—Table supplement 1, Figure 3—Table supplement 6). To further understand how these different genes collectively contribute to the individuality of each breast cell type, we needed to identify and evaluate the cellular pathways in which these genes are collectively involved—and the higher-order processes they bestow.

### Defining the nature of each breast cell type

To acquire a broad perspective of each breast cell type’s fundamental properties and individual functions, we performed a wholesale assessment of the biological pathways and processes disproportionately employed by these different cell types. Gene set enrichment analysis (GSEA)^52–54^ accounted for each of the 20,000+ expressed genes in these cells’ transcriptomes. The algorithm determined whether a particular pathway’s genes were enriched within a bidirectionally ranked list of genes constructed from two contrasted cell types. One advantage of GSEA is that it doesn’t require large transcriptional variations to pinpoint enriched pathways, as it can identify situations where many genes in a pathway differ in a small but coordinated fashion.

Several databases organize functionally related genes into distinct pathways. Here, we present our GSEA results that used gene sets defined by the Reactome Knowledgebase^18^. This database contains a manually curated list of 2,585 pathways cross-referenced to other scholarly resources, such as NCBI^55^, ENSEMBL^56^, KEGG^57^, and GO^17, 20^. Each Reactome pathway represents a collection of genes whose products coordinate to perform a specialized biological process. To determine which of these pathways are potentially active—or differentially enhanced—in a particular cell type relative to another, we analyzed every transcript, in each breast cell type, across every cell-cell comparison, which required 55 separate differential gene expression contrasts (Figure 20, Figure 3—Table supplement 7 [1v2]).). For each pairwise comparison, we calculated: a) the log fold change for each measured transcript, b) the *p*-value for the calculated difference (Wald test), and c) the Benjamini-Hochberg adjusted *p*-value. Using these values, we ranked all expressed genes bidirectionally, placing the most statistically significant genes at the extreme ends of the list, with one cell type represented at the top (most positive) —and the other at the bottom (most negative). After ranking the genes, we performed GSEA, which extracted pathways from the Reactome database, identified the position of each pathway’s genes within our ranked list, and used a running sum algorithm to calculate a normalized enrichment score for each pathway. For each pairwise comparison, hundreds of enriched pathways were typically identified, each consisting of 15 to 200 genes.

**Figure 20.**
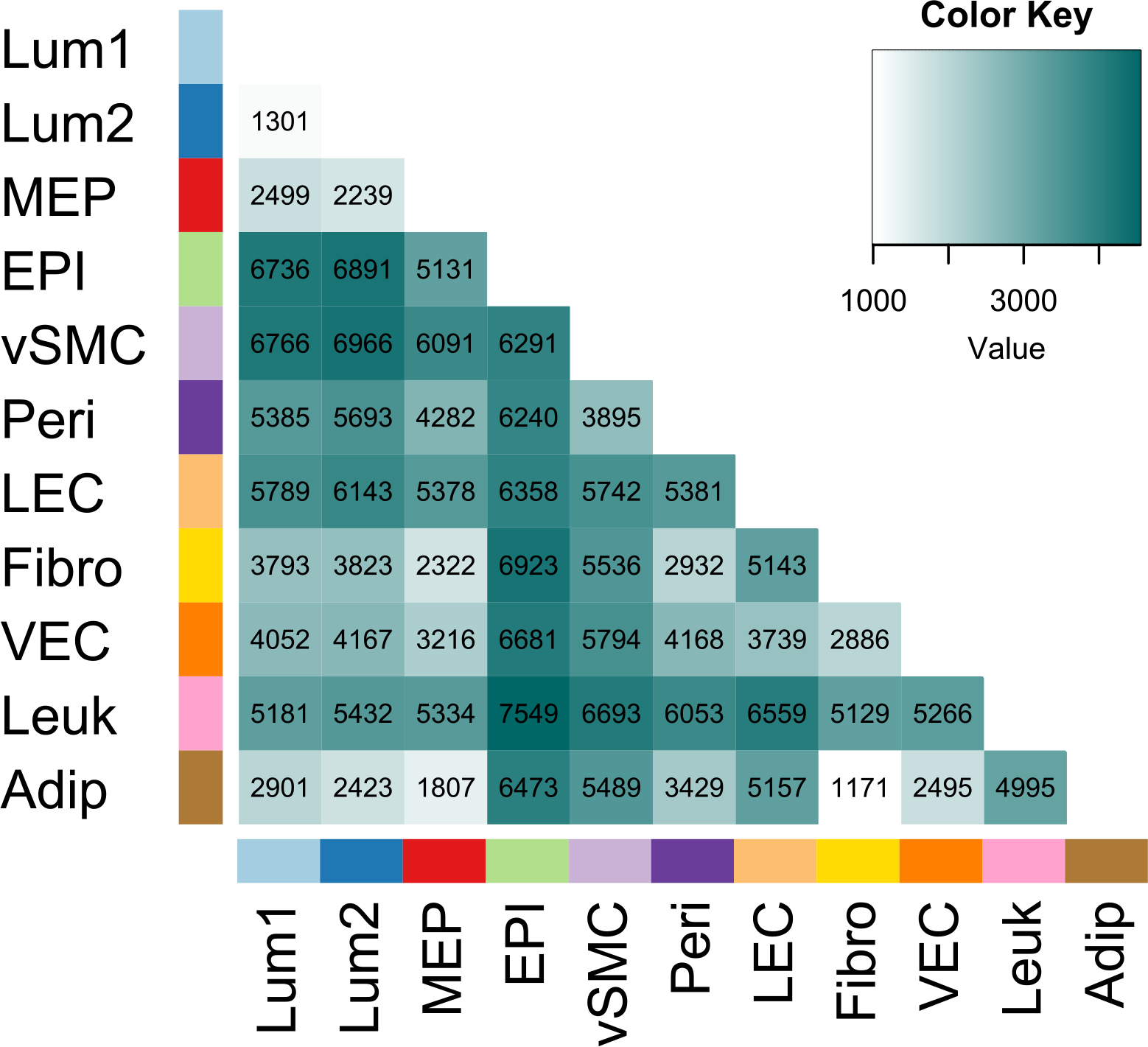
Differentially expressed genes across all cell type pairs. The number of differentially expressed genes between cell-type pairs, as reported by DESeq2; Benjamini-Hochberg adjusted *p*-values ≤ 0.1.

To help us navigate the overwhelming amount of data generated from these combined analyses, we integrated our GSEA results and gene expression values into a single pathway analysis tool that we programmed within Microsoft Excel (Pathway Analyzer^9^). This facile tool efficiently summarizes results according to user inputs (e.g., cell types to contrast, FDR cutoffs to use, biological themes to investigate, specific pathways to examine, and genes to explore).

With the aid of the Pathway Analyzer tool, we found that the number of pathways enriched in each cell type varied widely, ranging between 7 to 462 (x̄ = 194, sd = 120). On average, fibroblasts had the fewest enriched pathways across all comparisons (x̄ = 85 ± 59), whereas those having the most were the vSMCs (x̄ = 315 ± 86), pericytes (x̄ = 272 ± 121), vascular endothelial cells (x̄ = 287 ± 126), and lymphatic endothelial cells (x̄ = 229 ± 114). When we compared cell types across different lineages, akin to the PCA analysis performed above, many of the same enriched pathways emerged, e.g., keratinization within the epithelial cell types, angiogenesis by endothelial cells, and more. Unfortunately, the magnitude and comprehensive nature of the GSEA results preclude us from presenting here every identified aspect. Instead, we will focus on the biology of closely related cell types since the broad features of different cell lineages were examined above. However, the results from all analyses are accessible, and one can thoroughly explore any of the 55 possible comparisons using the Pathway Analyzer tool.

Below, we present the most thought-provoking findings from six cell-type comparisons; these are: a) the two luminal epithelial cell types, b) ER^Neg^ luminal epithelial cells and myoepithelial cells, c) ER^Neg^ luminal epithelial cells and vascular endothelial cells, d) the two endothelial cell types, e) fibroblasts and adipocytes, and finally f) fibroblasts and pericytes. Some enriched pathways we anticipated, such as finding the enrichment of ‘estrogen signaling’ in ER^Pos^ luminal cells. These results validated the analytic approach and helped us establish confidence in the pathway enrichments.

### Estrogen signaling in luminal epithelial cells: a validating example

Luminal epithelial cells help shape the breast’s glandular structure by forming the lumen of mammary ducts. The breast’s two major luminal cell types are commonly classified by their divergent staining for estrogen receptor alpha (ERα), and we were able to isolate each of these cell types based on their mutually exclusive staining for α6 integrin (ITGA6/CD49f; Figure1, Figure 1—figure supplement 1)^8^. To determine the distinguishing characteristics of these two cell types, we contrasted their expressed transcripts, exposing 1,301 differentially expressed genes (BH/FDR adj. p-value < 0.1, Figure 21). ER^Pos^ luminal cells expressed higher levels of 546 of these, including progesterone receptor (PGR), epiregulin (EREG), fibronectin (FN1), and FOXA1. In contrast, transcripts differentially expressed by the ER^Neg^ cells included epidermal growth factor (EGF), matrix metallopeptidase 7 (MMP7), KIT, and 752 other genes (Figure 21, Figure 3—table supplement 7 [1v2]). We ranked all expressed genes by their differential expression (based on their fold-change and statistical significance) and performed pathway enrichment analysis to identify the cellular functions impacted by these cell types’ transcriptional differences.

**Figure 21.**
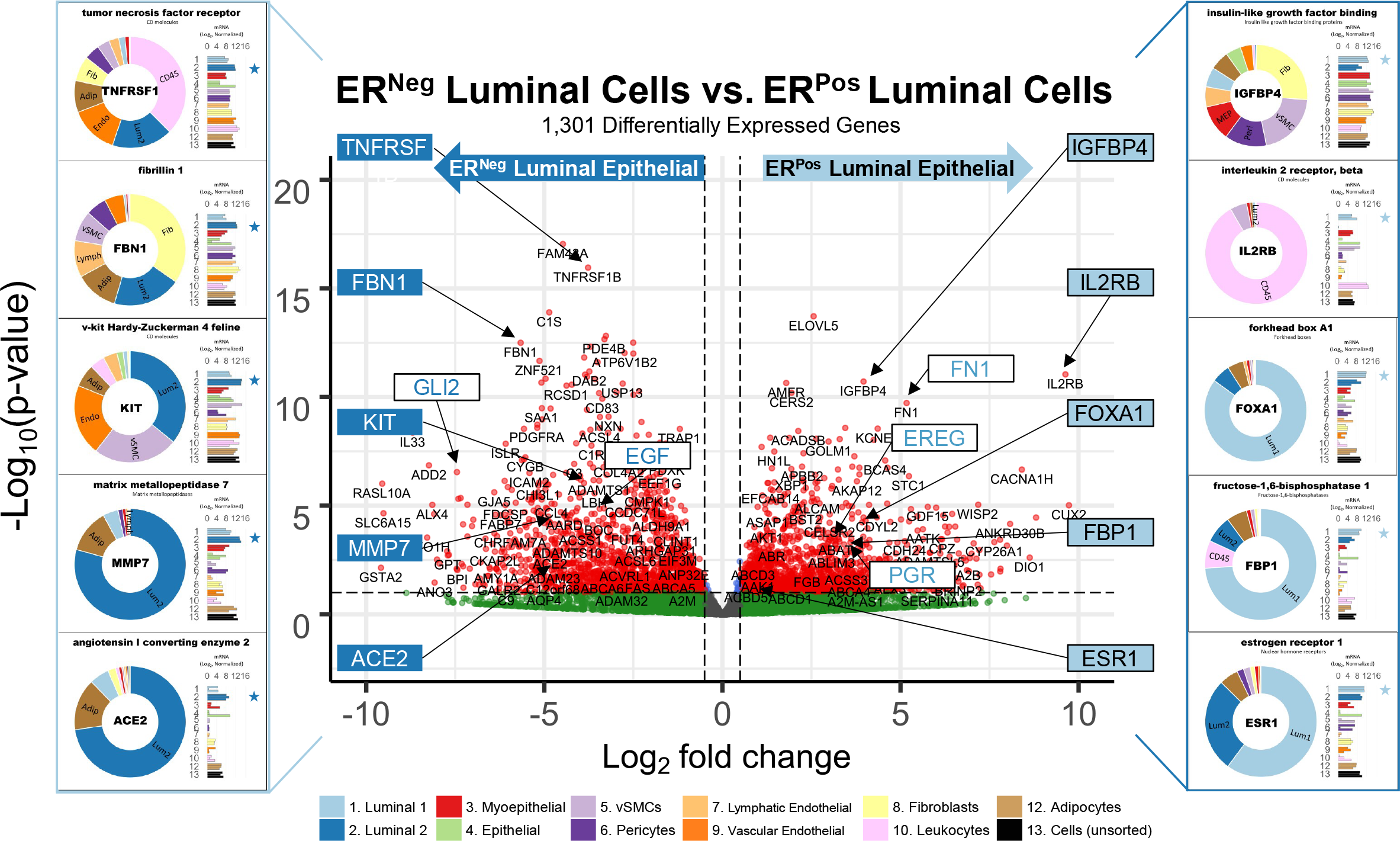
Differences between ER^Pos^ and ER^Neg^ luminal epithelial cell types. A contrast of expressed transcripts between the two luminal epithelial cell types revealed 1,301 differentially expressed genes (DeSeq2/Benjamini-Hochberg adjusted p-value ≤ 0.1). All expressed genes are displayed. The horizontal dashed lines (separating red and green dots) represents an Adj. *p*-value of 0.1. Established cell type markers (of each cell type) are denoted. mRNA expression values for ten genes (indicated by filled, color-coordinated boxes) are shown at left and right of the volcano plot. Color-coded stars in the bar graphs indicate the cell type being contrasted that had the higher level of that gene; i.e., ER^Pos^(light blue) and ER^Neg^ luminal cells (dark blue).

Gene set enrichment analysis (GSEA) assessed where each pathway’s genes fell along our ranked gene list, and the algorithm calculated a normalized enrichment score, nominal *p*-value, and false discovery rate (FDR) for each Reactome pathway. The *estrogen-dependent gene expression* set was among the top pathways enriched in the ER^Pos^ luminal cells (*p*-value <0.001, *q-*value =4.84x10^-4^). Identifying this estrogen signaling pathway was validating and served as a fine example of the enrichment algorithm (Pathway Analyzer [1v2]). Differentially expressed genes contributing to this pathway’s enrichment included progesterone receptor (PGR), FOXA1, erb-b2 receptor tyrosine kinase 4 (ERBB4), GATA binding protein 3 (GATA3), estrogen receptor a (ESR1), MYC, and others (Figure 22a). These genes were among the 65% in the pathway that fell within the leading edge of the analysis (the leading-edge genes are the core genes that contributed most to the GSEA enrichment score, Figure 22a). Furthermore, when we analyzed transcripts from this estrogen-related pathway, we confirmed that the leading-edge genes were indeed differentially expressed between the ER^Pos^ and ER^Neg^ luminal epithelial cells (Figure 22b, Pathway Analyzer).

**Figure 22.**
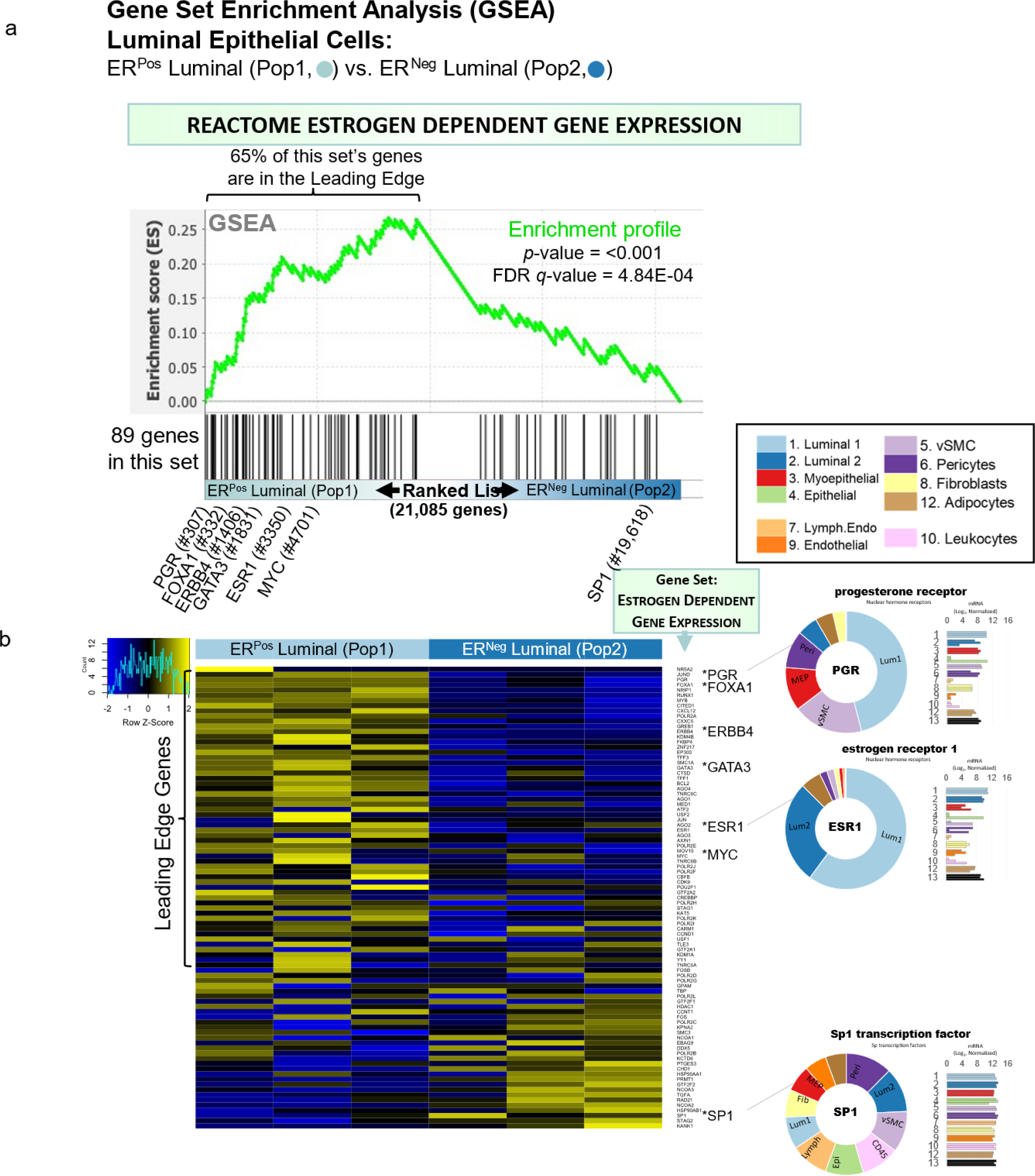
Inspection of GSEA results and the validating example of the *estrogen dependent gene expression* pathway. (a) Gene Set Enrichment Analysis (GSEA) was performed using a list of genes expressed by ER^Pos^ and ER^Neg^ luminal epithelial cells, ranked by the statistical significance of their differential expression, *-log10 {p-value} * sign([Fold Change]).* The ranked list, depicted by the shaded bar, has the most statistically significant genes located at its extreme ends, with the ER^Pos^ cells’ most differentially expressed genes represented at the left end and the ER^Neg^ cells’ most differentially expressed genes on the right. GSEA identified the Reactome pathway *Estrogen dependent gene expression* as one of the the most statistically significant enriched pathways identified (FDR<0.001). The position of the 89 genes composing the *estrogen dependent gene expression* pathway within the ranked list of expressed genes are depicted by vertical black lines. Names and positions of several familiar genes in the pathway are indicated, e.g., progesterone receptor (PGR) is found at position 337 (out of 21,085 expressed genes). The GSEA algorithm assessed the distribution of the pathway’s 89 genes within the ranked list using a running sum score (green trace), calculating statistics by comparing results to analysis of a randomized ranked list (1000 permutations). **(b)** Heat map depicting transcript levels within ER^Pos^ and ER^Neg^ luminal epithelial cells of the 89 genes belonging to *estrogen dependent gene expression* pathway, ordered by their relative position in the ranked list (VST values normalized by row). Positions of the familiar genes are indicated, including SP1 (which was near the bottom of the list and was not differentially expressed). Transcript levels of PGR, ESR1, and SP1 in each cell type (rlog values on log2 (bar graph) and linear (donut graph) scales).

Apart from *estrogen-dependent gene expression,* we identified an additional 324 enriched pathways between the two luminal cell types, 211 within the ER^Pos^ cells and 113 within the ER^Neg^ cells (FDR ≤ 10%, Pathway Analyzer). We used a pathway enrichment and visualization method to extract information rooted within these GSEA results (Enrichment Map^58^). This tool analyzed the pathway’s gene sets and, based on the proportion of genes they shared, formed a complex network of interconnected pathways, placing redundant and biologically related pathways close to each other. We connected pathways when they shared 50% or more genes, and this caused distinct clusters, or ‘biological themes,’ to emerge in the network (Figure 23, Supplemental Cytoscape file [Enrichment Map 1v2]). We circled these grouped pathways and named their overarching theme using Reactome’s hierarchy nomenclature as a guide. For example, the *estrogen-dependent gene expression* gene set clustered with the two other sets related to estrogen signaling: *estrogen-receptor mediated signaling* and *extra-nuclear estrogen signaling.* We encircled all three on the map and labeled this overall theme ‘Estrogen Signaling.’ (Figure 23). We adhered to this strategy for each theme in the network and often arranged related themes together. For example, we placed the enriched *‘*FOXO mediated transcription’ theme adjacent to ‘Estrogen Signaling’ on the enrichment map because FOXO3 is an established ERα regulator ^59, 60^.

**Figure 23.**
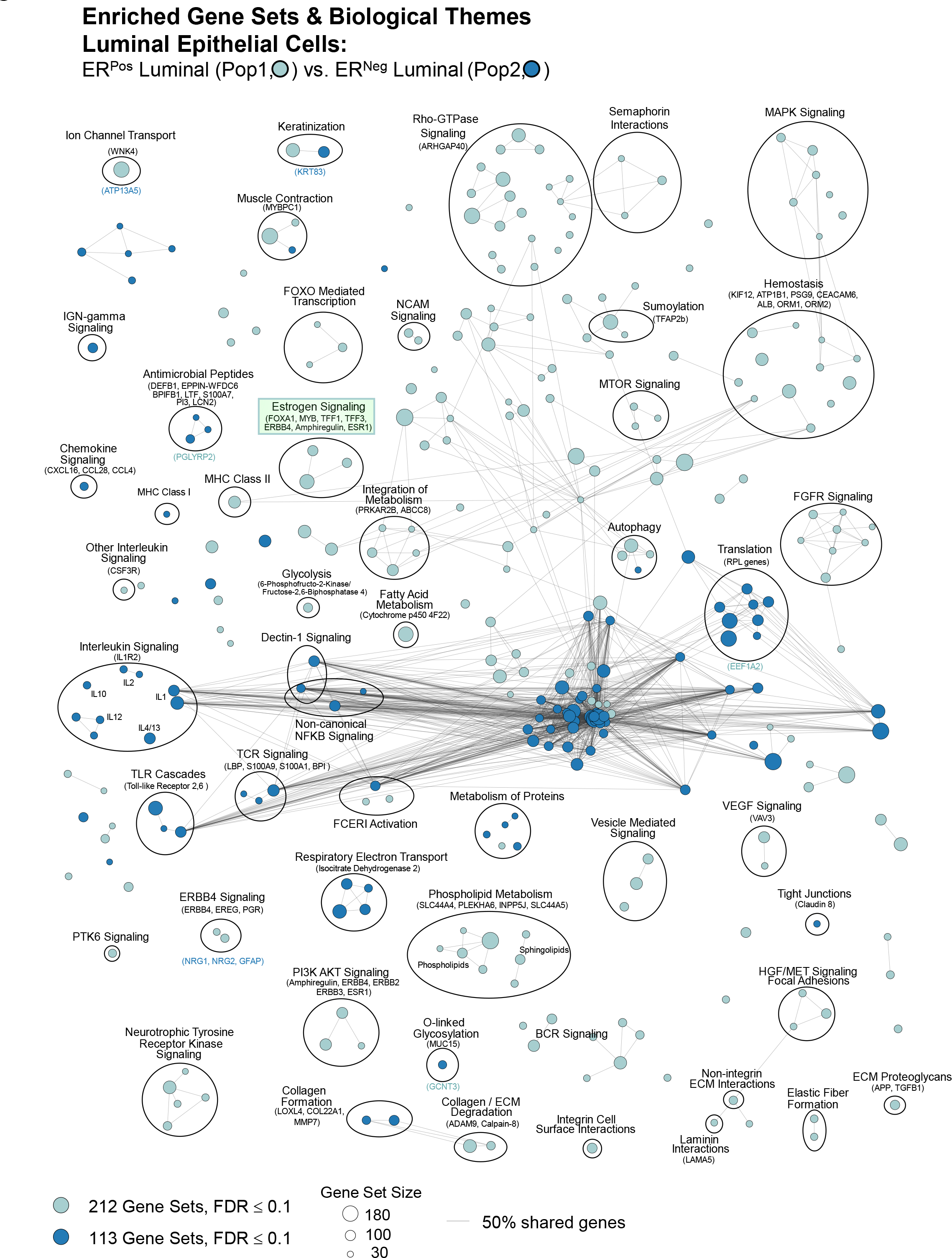
ER^Pos^ and ER^Neg^ luminal cells’ enriched pathways. Visualization of pathways enriched among ER^Pos^ and ER^Neg^ luminal cells (Enrichment Map/Cytoscape). All enriched pathways with a FDR≤0.1 appear on the map (nodes, color coded by cell type). Pathways(nodes) are connected when they share 50% of their genes (or greater). Node size corresponds to the number of genes in the pathway, as indicated (bottom left). Biologically related pathways sharing genes formed clusters in the network (biological themes), and different themes with a common biological function (e.g., metabolism-, immune -, and ECM-related, etc.) were typically placed adjacent to each other (when possible). Select themes were encircled and annotated. Highly differentially expressed genes are indicated (above the theme when it coincided with the attributed cell type—and below if it was expressed by the contrasted cell type). The names of each pathway are available in the supplemental Cytoscape file and Pathway Analyzer tool.

To evaluate the GSEA results and biological rationale for each pathway’s enrichment, we studied the genes from each pathway—and their expression levels in the different breast cell types.

These details often provided valuable insights (Pathway Analyzer). For example, high-scoring genes within the *ER mediated signaling* pathway included: HSBP1, which binds to ERα and promotes membrane localization; JUND, which interacts with ERα and occupies estrogen response element sites within the promoters of target genes; progesterone receptor, which is a direct downstream target of ERα; and others, like FOXA1, AKT1, epiregulin (EREG), amphiregulin (AREG), and SRC proto-oncogene. These genes were all highly expressed by ER^Pos^ luminal cells, and their transcript levels in these cells stood in stark contrast to those we observed within ER^Neg^ luminal cells –and the same was observed for FOXO mediated transcription (Figure 23—figure supplement 1a,b). In some cases, a pathway’s genes were uniquely expressed (or nearly so) by one of the contrasted cell types. We indicated these genes on the enrichment map under each theme’s name. For example, the genes encoding estrogen receptor a (ESR1), amphiregulin growth factor (AREG), transcriptional factors FOXA1 and MYB; tyrosine kinase receptor ErbB4, and the trefoil factor mucosal proteins TFF1 and TFF3 were all highly and differentially expressed by ER^Pos^ luminal cells (Figure 22b, Figure 23).

Collectively, these data provided strong evidence that estrogen signaling is active in this cell type—as one would anticipate in cells expressing estrogen receptor.

Identifying the association between estrogen signaling pathways and ER^Pos^ cells was validating—as was later finding additional anticipated themes in other cell-type contrasts. Moreover, these correlations assured us that the pathway analysis had indeed identified processes used disproportionately between cell types. Below, we move on from the detailed example of estrogen signaling in these luminal cells and sift through the complexity and sheer volume of remaining results.

### ER^Pos^ vs. ER^Neg^ luminal epithelial cells (Pop1 vs. Pop2)

When we compared transcripts expressed by ER^Pos^ and ER^Neg^ luminal cell types (FACS populations #1 and #2, Figure 1a), we identified 325 enriched pathways at a false discovery rate of 10%. The majority of these pathways were associated with cell signaling (83 pathways), metabolism (55 pathways), and immune-related functions (42 pathways). Others involved extracellular matrix organization, developmental biology, nervous system development, cell signaling, muscle contraction, and vesicle-mediated transport (Figure 23, Pathway Analyzer).

Further exploration of expressed transcripts defined by each pathway showed that the GSEA algorithm functioned well at identifying significant, concordant differences in the gene levels between cell types—even if individual transcript levels only modestly diverged. There were, however, pathways that contained a significant share of highly differentially expressed genes— some of these being principal players in the pathway. For example, fibronectin (FN1) and TGFB1 are members of the *ECM proteoglycans* gene set and were expressed at over 35- and 10- fold higher levels in ER^Pos^ luminal cells than their ER^Neg^ cell counterparts (Adj. *p*-value = 3.2x10^-8^ and 3.8x10^-7^, Figure 23, Figure 3—table supplement 7 [1v2]). Because implications of these stark differences were often patently evident, we primarily focus on pathways that contain these, but details of all enriched pathways can be thoroughly examined with the provided analysis tool (Pathway Analyzer).

#### Cell Signaling & Signal Transduction (1v2)

The PI3K/Akt, ErbB4, VEGF, and HGF/MET pathways are all notable receptor tyrosine kinase pathways that we found enriched within ER^Pos^ cells (Figure 23, Pathway Analyzer, Supplemental Cytoscape file [Enrichment Map 1v2]). Along with p53, Phosphatidylinositol 3-kinase (PIK3CA) is the most mutated gene in breast tumors (36% of tumors), and genes in the phosphoinositide 3-kinase (PI3K/AKT) pathway are the most frequently altered in human cancers^61^. Across the span of normal breast cell types, we found PIK3CA to be evenly expressed (Figure 23—figure supplement 1c). The PI3K/AKT pathway’s enrichment in ER^Pos^ luminal cells was instead primarily driven by the differential expression of the serine-threonine kinase AKT1, the EGFR ligand epiregulin (EREG), and the receptor tyrosine kinases: fibroblast growth factor receptor 1 (FGFR1) and platelet-derived growth factor receptor beta (PDGFRB). Other genes contributing to the pathway’s enrichment included the EGFR receptors ERRB2, ERBB3, and ERBB4 (Figure 23—figure supplement 1c). These data show that the ER^Pos^ luminal cells contain different signaling elements than ER^Neg^ cells and possess the potential for stimulating themselves and neighboring cell types.

Another prominent member of the epidermal growth factor family of receptors is ERBB4. This receptor dimerizes with ERα and is involved in many facets of downstream ER signaling^62^. Not surprisingly, we found ER^Pos^ luminal cells enriched with the ERBB4 pathways: *signaling by ERBB4* and *nuclear signaling by ERBB4* (FDR=3.9% and 2.9%, Figure 23). This enrichment was driven by the cell’s prominent expression of ERBB4 and its ligands betacellulin (BTC) and epiregulin (EREG), although ER^Neg^ luminal cells differentially expressed several pathway members as well, including neuregulins 1 & 2 (NRG1, NRG2). In addition, glial fibrillary acidic protein (GFAP), another pathway member, was also expressed—nearly exclusively—by ER^Neg^ luminal cells. In CNS astrocytes (cells that express high levels of GFAP), GFAP promoter activity is inhibited when bound by an erbb4-containing complex^63^. This inhibition perhaps explains the reduced GFAP levels in ER^Pos^ cells, as these cells have high ERBB4 levels (Figure 23—figure supplement 1d).

Another theme emerging from the analysis of the two luminal cell types was VEGF signaling, as we found the pathways *signaling by VEGF* and *VEGFR2 mediated vascular permeability* enriched within ER^Pos^ luminal cells (FDR=0.17% and 4.2%, respectively). This result is, however, an excellent example of how comprehensive information on all the breast cell types provided substantial perspective. Between the two luminal cell types, the VEGF pathways were indeed enriched in the ER^Pos^ cells—predominantly due to their differential expression of the serine/threonine kinase AKT1, amyloid beta precursor like protein 2 (APLP2), neuropilin 1 (NRP1), and other pathway members. However, when we explored these transcript levels across all cell types, we found their expression in ER^Pos^ cells was not dramatically different from most other breast cell types (Figure 23—figure supplement 1e). It thus became clear that the pathway’s enrichment was not a feature of its overexpression in ER^Pos^ cells but rather its relative repression in ER^Neg^ luminal epithelial cells (Pathway Analyzer). Because of the importance of these comparative insights, we similarly evaluated gene expression across all cell types for each pathway, using heat maps and the Pathway Analyzer tool.

Another pathway significantly enriched in ER^Pos^ luminal cells was HGF/MET. Hepatocyte growth factor (HGF) was first identified as a fibroblast-secreted factor that caused dramatic epithelial cell migration^64^. Its cognate receptor is MET receptor tyrosine kinase^65^. We found three HGF/MET signaling pathways enriched within ER^Pos^ luminal cells, including *signaling by MET* (FDR=0.6%, Figure 23, Pathway Analyzer). Enrichment was mainly driven by the differential expression of fibronectin, integrin alpha 3 (ITGA3), and several laminin subunits (LAMA1, -A5, and -B2). These proteins are involved in focal adhesion signaling that acts through focal adhesion kinase (PTK2), which was also modestly elevated in ER^Pos^ cells (Figure 23—figure supplement 1f). Interestingly, when we examined HGF expression across all cell types, we found, consistent with its initial discovery, that it was indeed highly expressed by breast fibroblasts, although pericytes and adipocytes also expressed it. HGF’s receptor, MET, was expressed by vascular endothelial cells, adipocytes, and the breast’s four epithelial cell types (Figure 22—figure supplement 1f).

#### Metabolism (1v2)

To evaluate metabolism-related pathways, we coded a filter in the Pathway Analyzer tool, allowing us to extract pathways involved in specific biological themes. At a 10% FDR threshold, we identified fifty-five enriched pathways related to metabolism—thirty within ER^Pos^ luminal cells and twenty-five within ER^Neg^ luminal cells. Those related to energy metabolism were enriched in ER^Pos^ luminal cells (Figure 23). Notable among these were: *integration of energy metabolism* (FDR=0.550%), *glycolysis* (FDR=7.94%), *regulation of insulin secretion* (FDR=5.51%)*, PKA activation in glucagon signaling* (FDR=4.87%), and *glucuronidation* (FDR=3.25%, Pathway Analyzer). Genes contributing to these pathways’ enrichment included: the inositol 1,4,5-trisphosphate receptor (ITPR1), which is a PKA-activated calcium channel that acts downstream of glucagon and cAMP signaling; fatty acid synthase (FASN), which catalyzes the synthesis of palmitic acid from acetyl-CoA and malonyl-CoA; and cAMP-dependent protein kinase (PRKAR2B), which transduces cAMP signals (Figure 23— figure supplement 1g). Elevated transcript levels of several glycolytic and regulatory enzymes within ER^Pos^ cells also led to the enrichment of the core glycolysis pathway. These were: hexokinase-1 (HK1), phosphofructokinase (PFKP), the bifunctional enzyme 6-phosphofructo-2-Kinase/Fructose-2,6-Biphosphatase 1 (PFKFB1), enolase gamma (ENO2), and pyruvate kinase M1/2 (PKM). These enzymes catalyze the first, last, and an intermediate step of glycolysis (HK1, PKM, ENO2) –and regulate other key regulatory metabolites in the pathway (PFKFB1),Figure 23, Figure 23—figure supplement 1h, Pathway Analyzer). Thus, significant evidence suggests ER^Pos^ cells have increased bioenergetic demands. As such, it was not surprising that we also found enrichment of pathways involved in lipid metabolism.

Metabolism of phospholipids, sphingolipids, and fatty acids were three themes that emerged from the comparison of the two luminal cell types (Figure 23). The ER^Pos^ cells’ disproportional expression of many of these pathways’ genes led to the enrichment *sphingolipid metabolism* (FDR=0.04%), *phosphoinositol metabolism* (FDR=0.06%), *fatty acid metabolism* (FDR=9.53%), and related pathways (Pathway Analyzer). Analysis of the leading-edge genes from these pathways demonstrated their elevated expression in the ER^Pos^ cells (Figure 23—figure supplement 1i-k). The most notable genes involved in phospholipid metabolism included the solute carriers SLC44A4 and SLC44A5. These membrane-embedded choline transporters provide a critical metabolic precursor for phosphatidylcholine synthesis. ER^Pos^ cells also highly expressed pleckstrin homology domain protein (PLEKHA6) and phosphatidylinositol 4,5-bisphosphate 5-phosphatase A (INPP5J). PLEKHA6 interacts with phosphatidylinositol 3,4,5-trisphosphate (PIP3), whereas INPP5J dephosphorylates PIP3. Fatty acid elongase (ELOVL5) and fatty acid synthase (FASN) are both involved in synthesizing fatty acids, were highly ranked, and contributed to the pathways’ enrichment.

The set of pathways involving protein metabolism covers the range of reactions that proteins encounter over their lifespan, from synthesis to degradation (including protein folding, post-translational modifications, and trafficking). We found both luminal cell types enriched with distinct protein metabolic pathways (Figure 23). In ER^Pos^ cells, two of the eleven significant pathways were *N-glycan antennae elongation in the medial trans golgi* (FDR=0.79%) and *sumoylation of intracellular receptors* (FDR=0.95%, Pathway Analyzer). Differentially expressed genes in the former pathway included carbohydrate (N-acetylgalactosamine 4-0) sulfotransferases (CHST8, CHST10), ST8 alpha-N-acetyl-neuraminide alpha-2,8-sialyltransferases (ST8SIA2, ST8SIA6), and mannosyl (alpha-1,3-)-glycoprotein beta-1,4-N-acetylglucosaminyltransferase isoenzymes (MGAT4A, MGAT4B). These enzymes are active in the medial Golgi, where N-glycan remodeling occurs. When we explored the second pathway, *sumoylation* of *intracellular receptors,* we found that the small ubiquitin-like modifiers (SUMO1, 2, 3) were surprisingly expressed evenly across cell types. The pathway’s enrichment was instead driven by elevated expression of the retinoic acid, retinoid X, progesterone, androgen, and estrogen receptors, indicating the receptors that could be modified by sumoylation (Pathway Analyzer).

The protein metabolism pathways enriched in ER^Neg^ cells included *amyloid fiber formation* (FDR=5.95%) and *O-linked glycosylation of mucins* (FDR=9.28%, Pathway Analyzer). We also observed that many, but not all, large ribosomal protein components were elevated in ER^Neg^ cells, leading to the enrichment of ribosome-involved pathways, *eukaryotic translation initiation* (FDR≤0.001%) and *eukaryotic translation elongation* (FDR≤0.001%). Historically, these genes have been considered invariant, but more recent evidence suggests otherwise, that levels may be linked to cell specification^66^—as we observed among the breast cell types here.

#### Extracellular Matrix Organization (1v2)

The extracellular matrix (ECM) is a dynamic structure consisting of a heterogeneous composition of scaffolding proteins, growth factors, proteoglycans, and remodeling enzymes. It undergoes persistent remodeling, primarily through reciprocal interactions with nearby cells in the immediate microenvironment. We found eleven pathways enriched between the two luminal cell types, with ER^NEG^ cells overexpressing genes related to *collagen synthesis* and assembly (FDR=1.58%), whereas the ER^Pos^ cells were enriched in degradative pathways *degradation of the extracellular matrix* and *collagen degradation* (FDR 4.08% and 5.55%, Figure 23, Pathway Analyzer). Driving enrichment of the synthetic pathway in ER^Neg^ cells were collagens (COL4A2, 6A2, 6A1, 22A1), lysyl oxidases (LOX, LOXL4), and matrix metallopeptidases (MMP3, MMP7). In contrast, the degradative pathway in ER^Pos^ cells was driven by differential expression of tissue inhibitor of metalloproteinase 2 (TIMP2), calpains (CAPN8, CAPN2, CAPN13), ADAM metallopeptidase 10 (ADAM10), cathepsin L (CTSL), elastin (ELN), matrix metalloproteinases (MMP12, MMP13), and collagens (COL7A1, 12A1, 4A5, 4A6). These data suggest that these two related—but distinct— luminal cell types have contrasting roles in the synthesis and degradation of specific ECM proteins (Figure 23—figure supplement 1 l,m).

Additional enriched pathways in ER^Pos^ cell relating to the ECM included: *laminin interactions* (FDR=4.1%), *non-integrin ECM interactions* (FDR=0.18%)*, integrin cell-surface interactions* (FDR=0.47%)*, syndecan interactions* (FDR=6.47%), and *ECM proteoglycans* (FDR=0.31%; Figure 23—figure supplement 1 n-r). Differentially expressed genes within these pathways included collagen IV isoforms A5 and A6 (COL4A5, COL4A6), laminin chains (LAMA5, LAMB2), fibronectin (FN1), transforming growth factor-beta -1 and -2 (TGFB1, TGFB2), platelet-derived growth factor beta (PDGFB), agrin (AGRN), synedecan 3 (SDC3), and integrins α5 and β3 (ITGA5, ITGB3), all of which had higher transcript levels in ER^Pos^ luminal cells compared to their ER^Neg^ counterparts.

#### Tight junctions (1v2)

Claudins are a family of genes we explored in the previous section, where we found, with few exceptions, that their transcription was mainly limited to endothelial and epithelial cells—predominantly the two luminal cell types (Figure 16a,b). Here, using GSEA, we identified significant differences between the ER^Pos^ and ER^Neg^ luminal cells (FDR = 9.1%). Specifically, the ER^Neg^ cells expressed significantly higher levels of CLDN10, CLDN16, CLDN11, CLDN8, and CLDN1, which led to enrichment of the *tight junction interactions* pathway (Figure 23, Figure 16a). This was notably the only Reactome ‘Cell-cell Communication’ pathway enriched in either cell type (other pathways, for example, involve *adherens junctions*, *cell-ECM interactions*, and *hemidesmosome assembly*, etc.). Despite the elevated levels of five claudins in ER^Neg^ cells, CLDNs 1, 7, 3, and 12 were expressed at roughly the same level in both cell types, whereas ER^Pos^ cells had higher levels of CLDNs 23, 6, and 9. Thus, the enrichment indicated that the two luminal cell types possess unique tight junction components to support their individual physiologies. These tight junctions are critical to the cells’ ability to form semi-permeable barriers, which helps retain fluid and protect against microorganisms in the outside environment –to which the ductal system is connected via the nipple opening.

#### Immune System (1v2)

Using GSEA, we identified forty-two enriched pathways related to immune function. The vast majority of these were in the ER^Neg^ luminal cells (33 pathways), which overlapped with the two innate immune system pathways previously identified by the PCA/overrepresentation analysis (Figure 10a). One of the larger themes was interleukin signaling, where we found the enrichment of nine pathways related to: *interleukin-4 and interleukin-13 signaling, interleukin-10 signaling, interleukin-1 family signaling, interleukin-2 family signaling,* and *interleukin-12 family signaling* (FDR= 0.181%, 0.792%, 2.33%, 4.79%, 1.096%, respectively; Figure 23; Pathway Analyzer). Enrichment of these pathways was driven by the ER^Neg^ cell’s differential expression of the interleukins 33, 1a, and 18 (IL33, IL1A, IL18), chemokine ligands 4 and 5 (CCL4, CCL5), tumor necrosis factor receptor TNFRSF1B, serum amyloid-A1 (SAA1), interleukin-2 receptor gamma (IL2RG), interleukin-1 receptor type II (IL1R2), interleukin-27 receptor alpha (IL27RA), colony-stimulated factor 2RB (CSF2RB), and lipocalin 2 (LCN2, Figure 23-figure supplement 1s,t; figure supplement 2a-c). We notably found breast macrophages concentrated near the epithelial structures and within the ducts and epithelial compartment^8^, and thus were not surprised to find luminal cell enrichment of interleukin pathways that involve macrophage activation. This indicates that the ER^Neg^ cells are disproportionately responsible for regulating macrophage physiology –a crosstalk that is subverted in breast tumors^67^.

Among the other enriched immune-related pathways, we discovered ER^Neg^ luminal cells expressed higher levels of genes involved in *toll-like receptor cascades* and *antimicrobial peptides* (FDR=8.208%, 4.1067%, respectively). The ER^Neg^ cells differentially expressed toll-like receptors 1 and 2 (TLR1 and TLR2), defensins B4A and B1 (DEFB4A and DEFB1), peptidase inhibitor 3 (PI3), and clusterin (CLU, Figure 23-figure supplement 2d,e). Furthermore, all ten toll-like receptors were highly expressed by luminal cells, with half being the highest in these cells compared to all other cell types (TLR1, TLR2, TLR2, TLR5, and TLR6, Figure 23— figure supplement 2d ; Figure 23—figure supplement 3a-f). Toll-like receptors play a vital role in the innate immune system by triggering an immune response to microbial agents. For example, TLR1 and TLR2 coordinate to mediate the response to bacterial lipoproteins. Although the toll-like receptors are well studied in the colonic epithelium, their contributions to the homeostasis of the breast epithelium are not as well characterized. Because luminal epithelial cells have the potential for elevated microorganism exposure, it is reasonable to expect TLR and antimicrobial peptide expression, as they are needed to provide innate immune protection.

#### Milk proteins (1v2)

Although not part of the Reactome gene sets, we noticed several genes encoding prominent milk proteins were highly differentially expressed by ER^Neg^ luminal cells, such as lactotransferrin (LTF) and casein kappa (CSN3). Lactotransferrin is the major iron-binding protein in milk, whereas casein kappa is critical for stabilizing casein micelle formation in milk. To explore the cellular expression of other milk constituents, we created a list of genes from prior reports of human milk composition ^68–70^. This list comprised 48 genes and included lactotransferrin, caseins, lactalbumin, and other milk proteins (Figure 23—figure supplement 4a). Most of these were expressed by at least one breast cell type (41/48), and nearly all were differentially expressed (37/41; BH/FDR adj. p-value < 0.05), with one of the two luminal cells most often being the cell type with the highest transcript level (25/37). After comparing expression between the luminal cell types, we found 12 genes whose expression significantly differed between the two (adj.*p-*val <0.05), with ER^Neg^ cells most often having the higher transcript levels (8/12). These eight genes were: lactotransferrin (LTF), secreted phosphoprotein 1 (SPP1), polymeric immunoglobulin receptor (PIGR), xanthine dehydrogenase (XDH), clusterin (CLU), complement component 3 (C3), haptoglobin (HP), and milk fat globule-EGF factor 8 protein (MFGE8, Figure 23—figure supplement 4b-j). These results demonstrate that the luminal cells are primed for milk production despite being derived from the resting breast tissues of non-lactating women and, at rest, transcribe the genes encoding major milk proteins–with the highest levels often being in the ER^Neg^ luminal cells.

#### ER^Neg^ luminal epithelial vs. Myoepithelial cells (Pop2 vs. Pop3)

Whereas the luminal cells shape the lumen of the gland’s ductal structure, myoepithelial cells form the basal layer, anchoring the epithelium to the basal lamina (Figure 9 a,d,e,g,h). The ER^Neg^ luminal and myoepithelial cells (FACS populations #2 and #3) are two of the most abundant cell types in the breast (Figure 1); they share the same epithelial lineage and spatial proximity—and physically interact. Most clinical breast tumors have a luminal phenotype and presumably originate from luminal cells, with the ER^Neg^ luminal cells being the most-likely cell of origin for basal-like breast tumors^49, 71^, despite these tumors’ basal-like features. For these reasons, we were interested in—and explored— the differences between ER^Neg^ luminal and myoepithelial cells, focusing on the biochemical and physiological processes distinguishing these two cell types.

Analysis of transcripts expressed by the myoepithelial and ER^Neg^ luminal cells revealed 2,239 differentially expressed genes (BH/FDR adj. *p*-value < 0.1, Figure 24, Figure 3—table supplement 7 [2v3]). ER^Neg^ luminal cells expressed the higher level of roughly half of these genes (51.6%), which included CD24, keratin 19, prominin 1 (PROM1), epidermal growth factor (EGF), and peptidase inhibitor 3 (PI3). Myoepithelial cells, in contrast, differentially expressed 1,083 genes; among these were genes previously associated with this cell type, including smooth muscle actin (ACTA2), keratin 5 (KRT5), p63 (TP63), and oxytocin receptor (OXTR, Figure 24). Using the *p-*values from this differential expression analysis, we created a ranked list of all expressed genes and used this list to perform pathway enrichment analysis. GSEA revealed enrichment of 426 Reactome pathways (FDR≤10%), which clustered into distinct biological themes (Figure 25). The most significant of these involved cell-cell communication, extracellular matrix organization, and metabolism (Pathway Analyzer).

**Figure 24.**
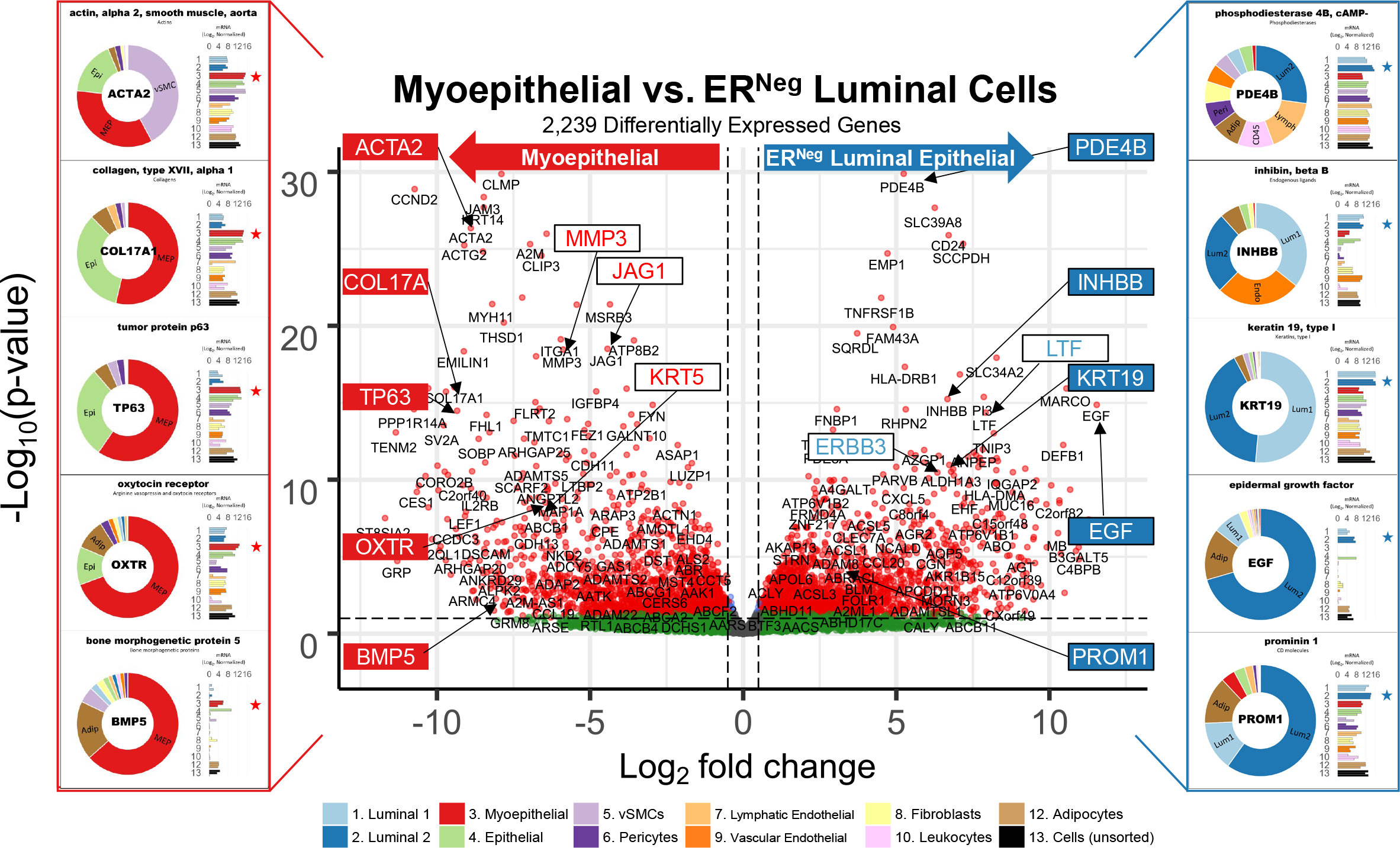
Differences between ER^Neg^ luminal and myoepithelial cell types. A contrast of expressed transcripts between ER^Neg^ luminal and myoepithelial cells revealed 2,239 differentially expressed genes (DeSeq2/Benjamini-Hochberg adjusted p-value ≤ 0.1). All expressed genes are displayed. The horizontal dashed lines (separating red and green dots) represents an Adj. *p*-value of 0.1. Established cell type markers (of each cell type) are denoted. mRNA expression values for ten genes (indicated by filled, color-coordinated boxes) are shown at left and right of the volcano plot. Color-coded stars in the bar graphs indicate the cell type being contrasted that had the higher transcript level, i.e., ER^Neg^(blue) and myoepithelial cells (red).

**Figure 25.**
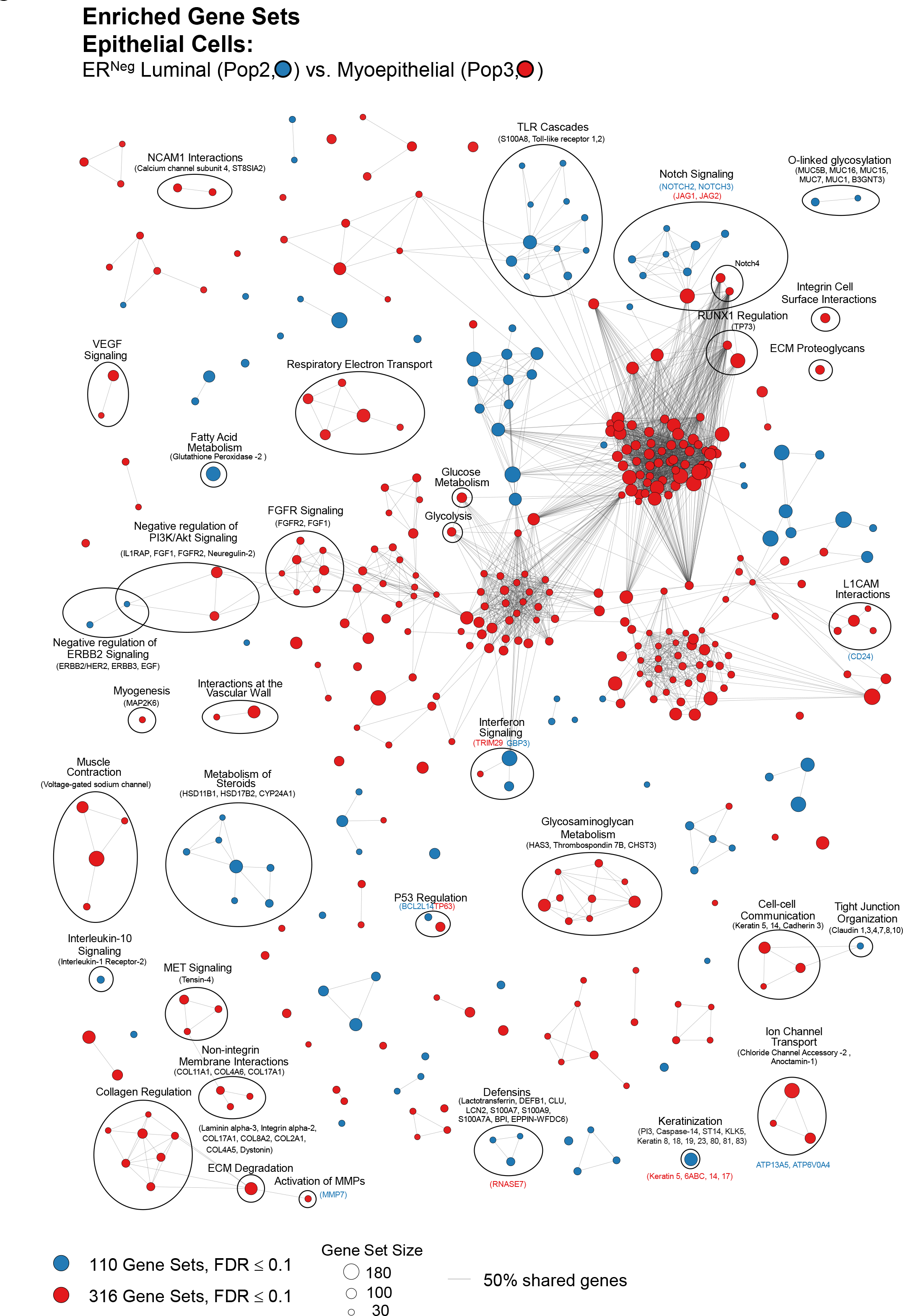
The ER^Neg^ luminal and myoepithelial cells’ enriched pathways. Visualization of pathways enriched among ER^Pos^ and myoepithelial cell types (Enrichment Map/Cytoscape). All enriched pathways with a FDR≤0.1 appear on the map (nodes, color coded by cell type). Pathways(nodes) are connected when they share 50% of their genes (or greater). Node size corresponds to the number of genes in the pathway, as indicated (bottom left). Select biological themes were encircled and annotated. Highly differentially expressed genes are indicated (above the theme when it coincided with the attributed cell type—and below if it was expressed by the contrasted cell type). The names of each pathway are available in the supplemental Cytoscape file and Pathway Analyzer tool.

#### Cell-cell communication (2v3)

Proper intercellular communication is essential for maintaining homeostatic balance and normal tissue function. When we contrasted the ER^Neg^ luminal and myoepithelial cells, we found enrichment of four cell-cell communication pathways: one in the ER^Neg^ luminal cells (tight junctions) and three in the myoepithelial cells (cell-junctions, cell communication, and ECM interactions). Given the previous identification of claudin expression in the luminal epithelial cells, it was not surprising to find the *tight junction interactions* pathway enriched in the ER^Neg^ luminal cells, as they—in comparison to myoepithelial cells— differentially expressed most epithelial claudins, such as CLDN1, 3, 4, 7, 8, 10, 12, and 16 (FDR =5.57%). The only significant exception was the myoepithelial expression of CLDN 11 (Figure 16 a,b). These results show that luminal cells express significantly more claudins than myoepithelial cells and thus rely on the tight junction features that these different claudins provide.

Although luminal cells differentially expressed claudins and were enriched with the *tight junction interactions* pathway, we found other cell communication pathways enriched in myoepithelial cells. These were the *cell-cell communication* pathway (FDR = 0.0919%) and its underlying pathways: *cell junction organization* (FDR =0.0076%) and *cell-extracellular matrix interactions* (FDR =0.1682%). Notable genes shared among these three pathways include actinin and filamin proteins that promote the branching of actin filaments and anchor the filaments to proteins in the cell’s membrane. Shared in the enriched pathways were also the cell adhesion molecule 3 (CADM3), keratins 5 and 14 (KRT5, KRT14), and cadherins 3, 4, and 11 (CDH3, CDH4, CDH11, Figure 25—figure supplement 1a). As we found above with the PCA-derived gene list for myoepithelial cells (myoepithelial and Pop4 epithelial cells), a general feature of myoepithelial cells is their expression of ECM and basement membrane components and formation of desmosomes involved in cell junction organization (Figure 14 a-c). Thus, we were not surprised to find myoepithelial enrichment of the *cell-extracellular matrix interactions* pathway because these cells form and help shape the epithelial basement membrane. For the same reason, we anticipated other pathways involved in extracellular matrix organization would be enriched within myoepithelial cells. We found this was indeed the case.

#### ECM organization (2v3)

As the non-cellular component of tissues, the extracellular matrix is formed by a network of proteins, proteoglycans, growth factors, and minerals that provides structural support, elasticity, and tensile and compressive strength to the tissue^72^. The ECM and basement membrane composition is dynamic and is constantly remodeled by cells in their immediate vicinity. Notably, we found pathways associated with extracellular matrix organization as some of the most significantly enriched pathways in myoepithelial cells, as nine of the ten most statistically significant pathways in the myoepithelial cells involved ECM and basement membrane organization (Pathway analyzer). Myoepithelial cells were enriched in nearly every aspect of ECM organization, including *Elastic fiber formation* (FDR= 0.01%), *Laminin interactions* (FDR<0.001%), *non-integrin membrane ECM interactions* (FDR<0.001%), *ECM Proteoglycans* (FDR<0.001%); *activation of matrix metalloproteinases* (FDR=4.38%), *collagen formation* and *collagen degradation* pathways (FDR<0.001%). These cells differentially expressed many collagen and laminin encoding genes, including COL2A1, 4A5, 4A6, 8A2, 11A1, and 17A1; and laminin chains LAMA1, LAMA3, and LAMB1). Other genes contributing to these pathways’ enrichment included elastin microfibril interfacer 1 (EMILIN1), metalloproteinase 3 (MMP3), tissue inhibitor of metalloproteinases 2 (TIMP2), actinin alpha 1(ACTN1), thrombospondin 1 (THBS1), fibromodulin (FMOD), heparan sulfate proteoglycan 2 (HSPG2), and syndecan 3 (SDC3), among others (Figure 25-figure supplement 1b-e). The myoepithelial cells—which reside on the basement membrane at the edge of the epithelial compartment— are thus a major contributor and remodeler of the basement membrane and ECM. This is a significant role that diverges from that of the luminal epithelial cells. However, the luminal cells were a major producer of matrilysin (MMP7) that, in addition to several ECM proteins, cleaves the milk protein casein, which we also found differentially expressed by ER^Neg^ luminal cells.

#### Muscle Contraction (2v3)

A defining feature of myoepithelial cells has long been their expression of smooth muscle actin^73^, which allows them to contract when stimulated by oxytocin, e.g., during lactation. It was thus reassuring to find *muscle contraction* (FDR = 0.0085%) and associated pathways enriched in myoepithelial cells, including *smooth muscle contraction, phase 0 rapid depolarization,* and *cardiac conduction—*the latter sharing similar genes despite being a pathway in the heart (FDR= 0.0707%, 2.011%, 0.148% respectively, Figure 25, Pathway Analyzer). Myoepithelial cells highly expressed smooth muscle actins alpha-1 and gamma-2 (ACTA1, ACTG2), the voltage-gated sodium channel alpha subunit 3 (SCN3A), the calcium voltage-gated channel subunit L (CACNA1C), myosin heavy chain 11 (MYH11), myosin light chain kinase (MYLK), and leiomodin 1 (LMOD1, Figure 25-figure supplement 1f) that contribute to the contractile ability of this cell type.

#### Ion channel transport (2v3)

Calcium is essential for myoepithelial contractility; Its intracellular availability is regulated by the transport of ions between the extracellular and intracellular spaces. This process was significantly enriched in myoepithelial cells, as we found *ion channel transport* and associated pathways enriched in myoepithelial cells (FDR=5.99%, Figure 25). The enrichment was driven predominantly by these cell’s differential expression of anoctamin-1 (ANO1), chloride channel accessory-2 (CLCA2), and the intracellular calcium channels ryanodine receptors 1 and 3 (RYR1, RYR3, Figure 25-figure supplement 1g).

Furthermore, myoepithelial cells differentially expressed the transient receptor potential (TRP) cation channels TRPV6, TRPC1, TRPC4, TRPM3, TRPC6, and TRPV2 (*TRP channels,* FDR=1.59%). TRP channels are calcium permeable and convert external stimuli into electrical or chemical signals^74^ (Figure 25-figure supplement 1g). These pathways likely contribute to regulating myoepithelial cells’ intracellular hyperpolarization/depolarization and contractility.

#### Cell signaling (2v3)

We identified the enrichment of eighty-one cell signaling pathways between the ER^Neg^ and myoepithelial cells. Most of these (sixty-two) were enriched within the myoepithelial cells. The bulk of these pathways involved receptor tyrosine kinases (23/62), G-protein coupled receptors (13/62), Notch (9/62), Wnt (8/62), and Hedgehog pathways (6/62). Although the ER^Neg^ luminal cells differentially expressed KIT and were enriched with both *signaling by SCF Kit* (FDR=5.7%) and ERBB2 signaling pathways, the myoepithelial cells were enriched with twenty receptor tyrosine kinase pathways, most at a conservative false discovery rate of less than 1% (13/20, Pathway Analyzer). These pathways involved MET, neurotrophic tyrosine receptor kinases, platelet-derived growth factor, vascular endothelial growth factor, fibroblast growth factor, and insulin-like growth factor pathways (Figure 25). For example, the enrichment of the FGFR signaling pathways *signaling by FGFR, signaling by FGFR1,* and *signaling by FGFR2* (FDR= 0.328%, 0.822%, and 4.00%,) was driven by their differential expression of fibroblast growth factors 1, 2, and 7 (FGF1, FGF2, FGF7); FGF receptors 1, 2, and 4 (FGFR1, FGFR2, FGFR4); and the ras proto-oncogene (HRAS, Figure 25- figure supplement 1h).

The enrichment of Notch signaling pathways split between the luminal and myoepithelial cells. *Signaling by notch* and *signaling by notch4* were enriched in myoepithelial cells (FDR= 0.0268%, 0.059%), as these cells differentially expressed smooth muscle actin (ACTA2) and notch ligands, JAG1 and JAG2 (Figure 25- figure supplement 1i). They also had slightly elevated levels of many proteosome subunits that help compose this pathway. In contrast, ER^Neg^ luminal cells were enriched with Notch 1, 2, and 3 signaling pathways (FDR= 3.38%, 7.75%, 1.99%) due to their differential expression of the Notch receptors, NOTCH2 and NOTCH3; the downstream effector HEY2, delta/notch-like EGF repeat containing (DNER) and epidermal growth factor (EGF), and other pathway members (Figure 25- figure supplement 1j).

Other myoepithelial enriched pathways involved G-protein coupled receptor-, Wnt-, and Hedgehog-signaling. Differentially expressed genes from the twelve enriched G-protein coupled receptor pathways included G-protein coupled receptor 176 (GPR176), dopamine receptor (DR), and adenylate cyclase 5 (ADCY5). Enrichment of the Wnt and Hedgehog signaling pathways was driven by elevated transcript levels of these pathways’ downstream effectors, which included transcription factor 7 (TCF7), lymphoid enhancer-binding factor-1 (LEF1), GLI1, and GLI3 (Pathway Analyzer).

#### Metabolism (2v3)

Within the eighty-seven metabolism-related pathways enriched between luminal and myoepithelial cells, one of the larger emergent themes involved the metabolism of glycosaminoglycans (Figure 25). These pathways were enriched in myoepithelial cells and included *glycosaminoglycan metabolism, chondroitin sulfate dermatan sulfate metabolism,* and *heparan sulfate heparin HS-GAG metabolism* pathways (FDR= 3.76%, 5.50%, and 3.04%, Figure 25). Among the genes contributing to their enrichment included key enzymes in GAG metabolism, e.g., a) hyaluronan synthase 3 (HAS3), which catalyzes the addition of acetylglucosamine and glucuronic acid monosaccharides to hyaluronan polymers during hyaluronic acid synthesis; b) chondroitin-6 sulfotransferase-3 (CHST3), which catalyzes the sulfation of chondroitin; and c) the heparan sulfate proteoglycan glypican-3 (GPC3), an ECM glycoprotein that binds to, and modulates, several signaling pathways (Figure 25—figure supplement 1k).

A characteristic feature of epithelial cells is their secretion and release of mucins from their apical surfaces^75^. Not surprisingly, we found the *O-linked glycosylation of mucins* pathway enriched in ER^Neg^ luminal epithelial cells (Figure 25). Luminal cells notably expressed mucin-1 (MUC1), which is a marker in our FACS panel and was used to sort the luminal cells. We found many other mucins expressed by the luminal cells, including MUC15 and MUC16 (Figure 25- figure supplement 2 a-f). Mucins are highly glycosylated proteins and a component of milk, where they provide antimicrobial resistance in the GI tract of breastfed infants^76^. These mucins are derived from the luminal epithelial cells.

The *metabolism of steroids* was another large theme in the network, which was enriched within ER^Neg^ luminal cells (FDR=0.998%, Figure 25, Pathway Analyzer). These cells differentially expressed hydroxysteroid dehydrogenases 17B2 and 17B4 (HSD17B2 and HSD17B4)— essential enzymes involved in the production of the androgen and estrogen sex hormones^77^.

Additionally, luminal cells highly expressed other steroid hormone regulators, HSD11B1 and steroid 5-alpha-reductase (SRD5A3), as well as the cytochrome p450s (CYP24A1 and CYP24B1), which function in Vitamin D metabolism (Figure 25- figure supplement 1l). We found these genes were highly expressed by both luminal cell types, indicating steroid metabolism is a general feature of the luminal lineage.

The ER^Neg^ luminal cells were also enriched with other metabolic pathways, such as *fatty acid metabolism* (FDR=6.04%, Figure 25, Pathway Analyzer), as they differentially expressed acyl synthetase long-chain family members 1, 3, 5, and 6 (ACSL1, ACSL3, ACSL5, ACSL6), aldo- eto reductase family member C3 (AKR1C3), glutathione peroxidase 2 (GPX2), and carnitine O-acetyltransferase (CRAT, Figure 25-figure supplement 1m). The acyl synthetase enzymes catalyze long chain fatty acids into active fatty acyl-CoA esters, which is a critical step in beta-oxidation and lipid biosynthesis. The roughly 10-fold difference in acyl synthetase transcripts between luminal and myoepithelial cells indicates that luminal cells have different metabolic demands requiring the processing of long-chain fatty acids.

In our analysis of the citric acid cycle pathway, we found most genes in this pathway were uniformly expressed across the entire spectrum of cell types. However, there were three notable exceptions. The first, which occurred in both luminal cell types, was their pronounced differential expression of both isoforms of isocitrate dehydrogenase (IDH1, IDH2), which operate in the mitochondria to regulate 2-hydroxyglutarate, alpha-ketoglutarate, and isocitrate (adj. p-values = 5.2e-5 and 2.1e-13). Another differentially expressed TCA cycle gene was pyruvate decarboxylase (PC), which was repressed in both endothelial cell types. Finally, we found phosphoenolpyruvate carboxykinase-1 (PCK1) expression was limited to pericytes and adipocytes (Figure 25-figure supplement 2f).

#### ER^Neg^ luminal epithelial vs. Vascular endothelial cells (Pop2 vs. Pop9)

Histologists consider endothelial cells specialized epithelial cells because they exhibit classical epithelial features^16^. For example, vascular endothelial cells and epithelial cells both form the lumen of their respective structures (e.g., vessels, ducts), use tight junctions for cell-cell adhesion, and rest on a basal lamina. However, when we analyzed the breast cell transcriptomes by PCA (Figure 7a) we found that endothelial cells projected more closely to mesenchymal cell types than to the epithelial cells, which may reflect their mesodermal origin. We found this curious and decided to compare the features of vascular endothelial cells to the abundant ER^Neg^ luminal epithelial cell type (FACS populations #9 and #2).

Comparing transcripts between vascular endothelial cells and ER^Neg^ luminal cells, we identified 4,167 differentially expressed genes (Figure 26). Of those, 1,652 were differentially expressed by the endothelial cells, which included endothelial markers: platelet/endothelial cell adhesion molecule (PECAM1/CD31), CD93, TIE1, and TEK tyrosine kinase. In contrast, luminal cells differentially expressed the transcriptional regulator, SMARCD3, keratin 15 (KRT15), CD24, cadherin 1 (CDH1), and transcriptional repressor GATA binding-1 (TRPS1, Figure 26, Figure 3—table supplement 7 [2v9]). We ranked all expressed genes by their degree of differential expression and used this list for pathway analysis. GSEA revealed five-hundred pathways enriched between ER^Neg^ luminal and vascular endothelial cells (FDR≤10%), which provided insights into the different biological pathways and systems used by each cell type (Pathway Analyzer).

**Figure 26.**
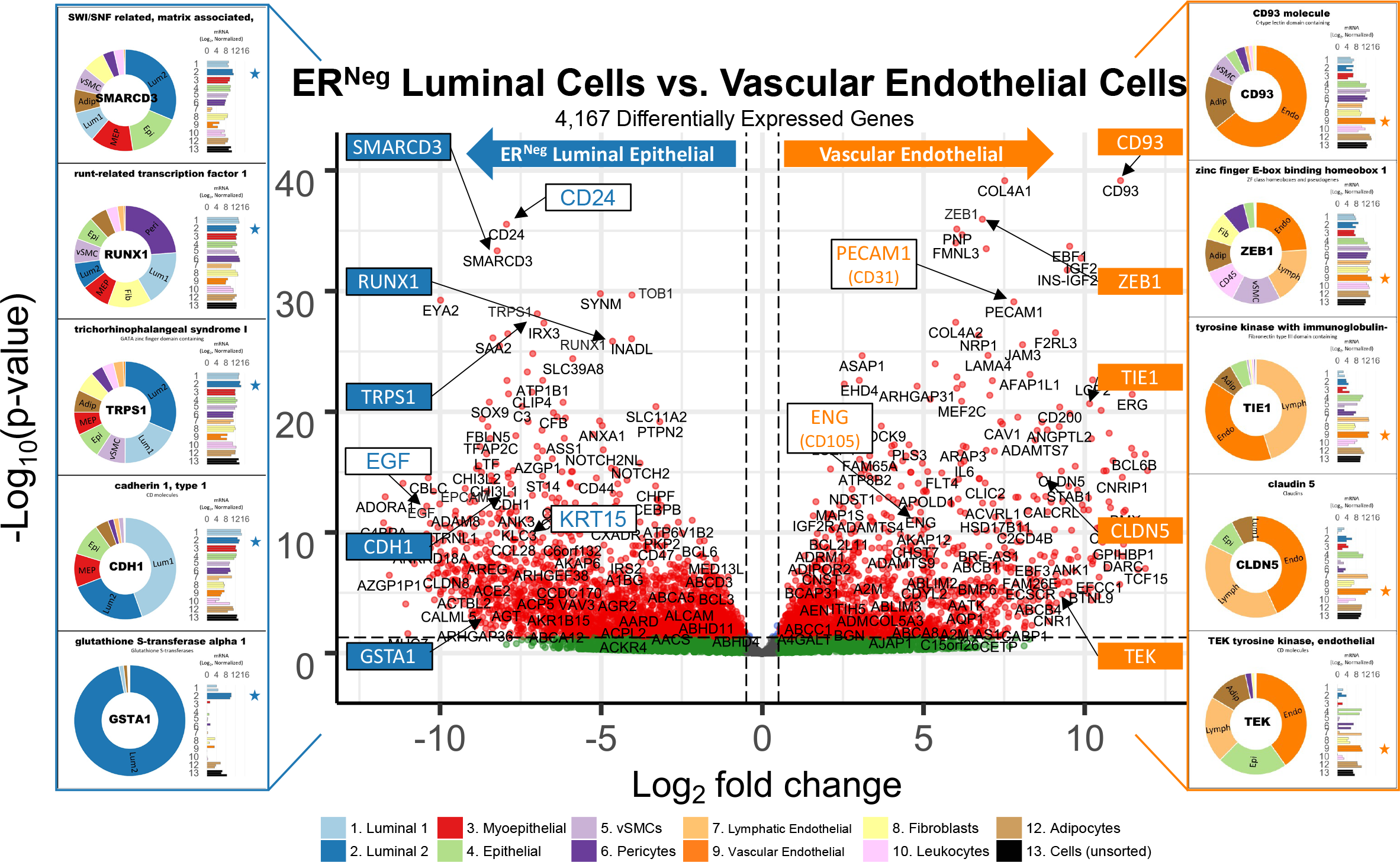
Differences between ER^Neg^ luminal and vascular endothelial cell types. A contrast of expressed transcripts between ER^Neg^ luminal and vascular endothelial cells revealed 4,167 differentially expressed genes (DeSeq2/Benjamini-Hochberg adjusted p-value ≤ 0.1). All expressed genes are displayed. The horizontal dashed lines (separating red and green dots) represent an adj. *p*-value of 0.1. Established cell type markers (of each cell type) are denoted. mRNA expression values for ten genes (indicated by filled, color-coordinated boxes) are shown at the left and right of the volcano plot. Color-coded stars in the bar graphs indicate the cell type being contrasted that had the higher transcript level, i.e., ER^Neg^(blue) and vascular endothelial cells (orange).

#### Hemostasis (2v9)

One of the most significantly enriched pathways, *cell surface interactions at the vascular wall* (FDR<0.001%), was found in vascular endothelial cells—cells that indeed form the vascular wall (Figure 27). Endothelial cells differentially expressed selectins E, L, and P (SELE, SELL, SELP)—which mediate cell-to-cell adhesion; endothelial cell adhesion molecule (ESAM); junctional adhesion molecule 2 (JAM3); platelet/endothelial cell adhesion molecule (PECAM1); and the angiopoietin receptor TEK tyrosine kinase (TEK, Figure 27-figure supplement 1a). These gene products regulate leukocyte adhesion to vessel walls during an inflammatory response and facilitate leukocyte migration. In addition, endothelial cells differentially expressed von Willebrand factor (VWF)— a historical marker of endothelial cells, thrombomodulin (THBD), and coagulation factors F2R and F3 (Figure 27-figure supplement 1b). These contributed to the enrichment of *formation of fibrin clot (clotting cascade,* FDR = 3.04%). This is an essential step for blood clotting after injury, indicating that endothelial cells supply critical factors for this process, thus distinguishing them from other epithelial cell types. Notably, several of these pathway’s genes are shared by mesenchymal perivascular cells (e.g., ITGA5, ESAM, ANGPT); this shared expression contributes to the mesenchymal alignment of endothelial cells in the PCA projection (Figure 7a).

**Figure 27.**
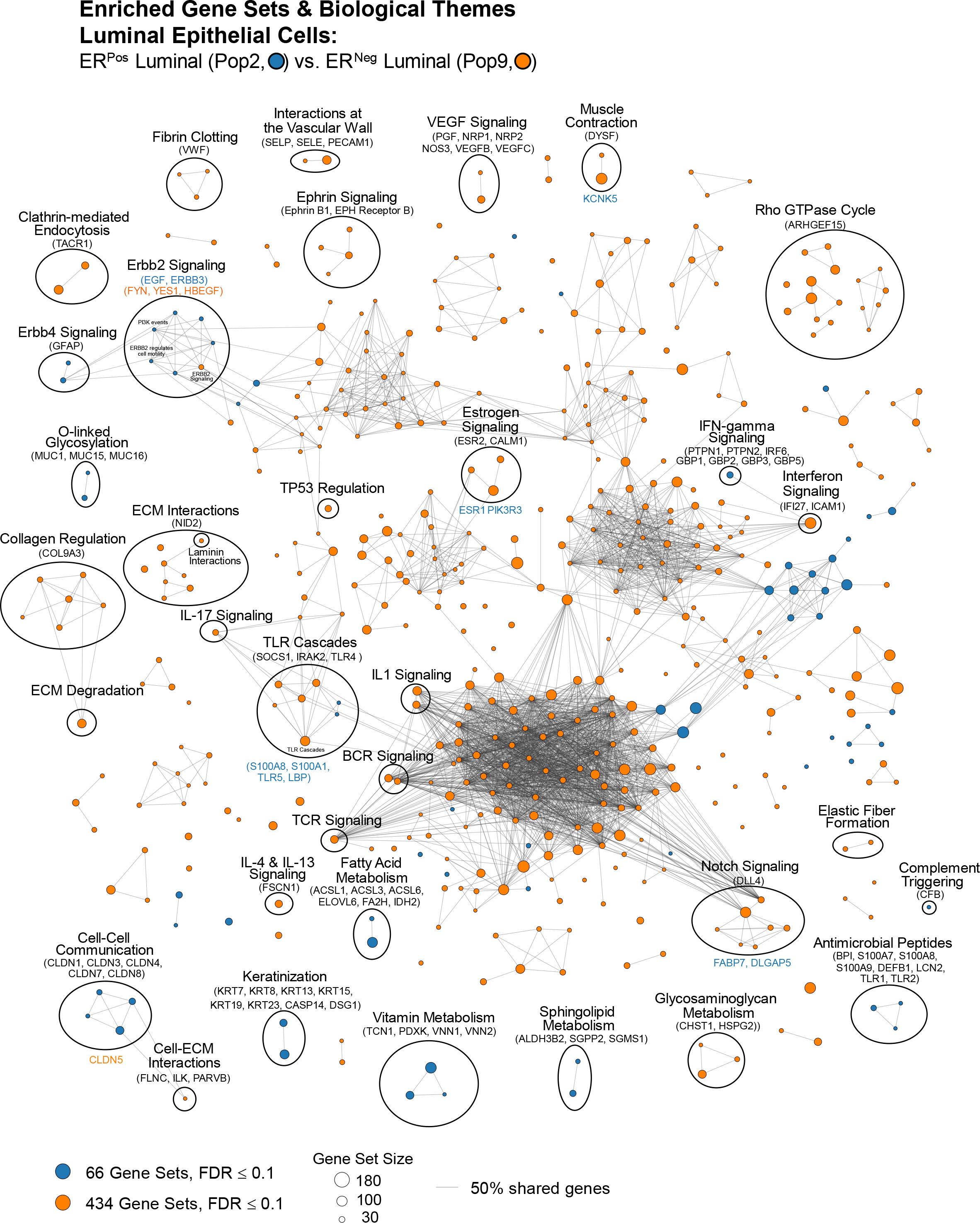
The ER^Neg^ luminal and vascular endothelial cells’ enriched pathways. Visualization of pathways enriched among ER^Pos^ luminal and vascular endothelial cell types (Enrichment Map/Cytoscape). All enriched pathways with a FDR≤0.1 appear on the map (nodes, color-coded by cell type). Pathways(nodes) are connected when they share 50% of their genes (or greater). Node size corresponds to the number of genes in the pathway, as indicated (bottom left). Select biological themes were encircled and annotated. Highly differentially expressed genes are indicated (above the theme when it coincided with the attributed cell type—and below if it was expressed by the contrasted cell type). Samples are color-coded. Due to space constraints and a large number of presented genes, their names (rows of the heatmap) must be inspected digitally (magnified in the PDF). The names of each pathway are also available in the supplemental Cytoscape file and Pathway Analyzer tool.

#### Keratinization (2v9)

Another divergent feature of endothelial and epithelial cells was the keratinization that occurred in ER^Neg^ luminal cells (*keratinization,* FDR<0.001%, Figure 27, Pathway Analyzer). Keratin filaments are heteropolymers of different Type I and II keratins that confer distinct biophysical properties to cells, connecting the cells’ cytoskeleton to other cells and the basement membrane via desmosomes and hemidesmosomes^78^. We found ER^Neg^ luminal cells differentially expressed fifteen keratins, including keratins 7, 8, 19, 6b, 23, 81, 15, 16, 17, and 18. In addition, they also differentially expressed caspase-14 (CASP14)—which is involved in the cornification of epithelia— and desmoglein-1 (DSG1), a component of desmosomes (Figure 27-figure supplement 1c).

#### Cell Communication (2v9)

Among the pathways associated with cell communication, we found that ER^Neg^ luminal cells were, compared to the vascular endothelial cells, enriched with the broad *cell-cell communication* pathway (FDR= 0.954%) and others involving tight junctions (Figure 27-figure supplement 1d). These were *cell junction organization* (FDR=1.8%) and *tight junction interactions* pathways (FDR= 0.835%, Figure 27-figure supplement 1e). The enrichment resulted from ER^Neg^ luminal cells’ differential expression of laminin beta 3 (LAMB3), cadherins 1 and 3 (CDH1, CDH3), as well as the crumbs-cell-polarity complex component 3 (CRB3), which helps form tight junctions. Although the vascular endothelial cell types expressed claudins 5 and 14 rather exclusively, the ER^Neg^ luminal cells differentially expressed an overwhelming number of claudins (CLDN1, -3, -4, -7, -8, -10, -11, and -12), leading to the pathway’s luminal cell enrichment (Figure 27-figure supplement 1e, Figure 16). Another cell-communication pathway is the *cell-extracellular matrix interactions* pathway, which we found enriched in vascular endothelial cells (FDR<0.001%). This pathway includes integrins that link ECM proteins to the cell’s cytoskeleton –with the help of scaffolding and focal adhesion proteins. Genes in this pathway differentially expressed by the vascular endothelial cells included filamin C (FLNC)—which was notably repressed by both luminal cell types (compared to all others), integrin-linked kinase (ILK), and parvin beta (PARVB, Figure 27-figure supplement 1f, Pathway Analyzer [2v9]).

#### Metabolism (2v9)

We identified eighty-two enriched pathways that involve the metabolism of nucleic acids, lipids, proteins, and other metabolites. Some of the more statistically significant pathways within ER^Neg^ luminal cells included the *metabolism of vitamins and cofactors* (FDR= 0.618%), *metabolism of water-soluble vitamins and cofactors* (FDR=0.71%), and *nicotinamide salvaging* (FDR=4.63%, Figure 27, Pathway Analyzer). Genes contributing to their enrichment included a) vanins 1 and 2 (VNN1 and VNN2), which are involved in vitamin B5 metabolism, b) transcobalamin 1 (TCN1)—a vitamin 12 binding protein, and c) pyridoxal kinase (PDXK), which phosphorylates vitamin B6 (Figure 27-figure supplement 1g). We also found *sphingolipid metabolism* enriched in luminal cells (FDR= 4.37%), as they differentially expressed sphingosine-1-phosphate phosphatase 2 (SGPP2), sphingomyelin synthase 1 (SGMS1), and aldehyde dehydrogenase 3 family member B2 (ALDH3B2, Figure 27- figure supplement 1h).

Additionally, *fatty acid metabolism* was enriched in ER^Neg^ luminal cells (FDR= 0.636). Enrichment resulted from the luminal cells’ differential expression of epoxide hydrolase 2 (EPHX2), ELOVL fatty acid elongase 6 (ELOVL6), acyl-CoA synthetase long-chain family members 1, 3, and 6 (ACSL1, ACSL3, ACSL6); fatty acid-2 hydrolase (FA2H); and the mitochondrial isocitrate dehydrogenase IDH2 (Figure 27- figure supplement 1i).

*Glycosaminoglycan metabolism* was another enriched pathway that contained substantial transcriptional differences (FDR=0.95%, Figure 27, Pathway Analyzer). Leading to its enrichment in vascular endothelial cells was their differential expression of carbohydrate sulfotransferase 1 (CHST1), versican (VCAN), biglycan (BGN), decorin (DCN), and heparan sulfate proteoglycan 2 (HSPG2/perlecan, Figure 27- figure supplement 1j). This is notable because endothelial surfaces and their surrounding ECM contain large quantities of heparan sulfate proteoglycans^79^ that modulate vascular functions, such as sequestering growth factors (FGF, VEGF), regulating lipoprotein metabolism, and inhibiting clotting^80^.

#### Immune system (2v9)

Pathways associated with the immune system were enriched in both luminal and endothelial cells (Figure 27). Nine were enriched in luminal cells and 39 in endothelial cells. *Antimicrobial peptides* (FDR=0.058%), *defensins* (FDR=0.349%), and *beta-defensins* (FDR=1.19%) were enriched in luminal cells, which differentially expressed defensin beta 1 (DEFB1), S100 calcium-binding proteins A7A, A7, A8, and A9 (S100A7A, S100A7, S100A8, and S100A9), toll-like receptors 1 and 2 (TLR1 and TLR2), lipocalin 2 (LCN2), and bactericidal/permeability-increasing protein (BPI, Figure 27- figure supplement 1k). Luminal cells also differentially expressed protein tyrosine phosphatase 2 and 6 (PTPN1 and PTPN6), interferon-inducible guanylate binding protein 1, 2, 3, and 5 (GBP1, GBP2, GBP3, and GBP5), and interferon regulatory factor 6 (IRF6)—which define the enrichment of *interferon-gamma signaling* in luminal cells (Figure 27, Figure 27- figure supplement 1l).

*Interferon signaling* was enriched in vascular endothelial cells, which differentially expressed intercellular adhesion molecule 1 (ICAM1) and interferon alpha-inducible protein 27 (IFI27, FDR= 0.512%, Figure 27, Figure 27- figure supplement 1m, Pathway Analyzer). ICAM1 encodes a receptor for leukocyte adhesion protein LFA-1 (ITGAL) and is essential for leukocyte trans-endothelial migration^81^. Additionally*, toll-like receptor cascades* (FDR=1.36%) were enriched in vascular endothelial cells, that differentially expressed suppressor of cytokine signaling (SOCS1), interleukin receptor-associated kinase 2 (IRAK2), and TLR4 (Figure 27, Figure 27- figure supplement 1n). Two pathways associated with TLR cascades were enriched in luminal cells: *IRAK4 deficiency TLR2/4* and *regulation of TLR by endogenous ligand* (FDR=0.926% and 4.94%, respectively, Figure 27). Notably, we discovered that enrichment of many pathways associated immune system was due to the expression of proteasome subunits.

For example, enrichment of *interferon signaling*, *TCR signaling, BCR signaling,* and *Interleukin- 1 signaling* were all affected by overexpression of the proteosome subunits PSMD2, PSMB6, and PSMB5, among others (FDRs < 0.001%, Figure 27, Figure 27- figure supplement 1o-q).

Proteosomes function in peptide processing for many pathways, including those involved in antigenic peptide presentation during immune responses. However, protein modification is required for many other pathways like cell cycle progression, apoptosis, and DNA repair.

#### Cell signaling (2v9)

As expected, *signaling by VEGF* and *VEGFR2 mediated vascular permeability* were enriched in vascular endothelial cells (FDR=0.005% and 6.81%, respectively), with differential expression of placental growth factor (PGF), neuropilin 1 and 2 (NRP1 and NRP2), nitric oxide synthase 3 (NOS3), and vascular endothelial growth factors B and C (VEGFB and VEGFC)—all of which are involved in VEGF-regulated angiogenesis (Figure 27, Figure 27- figure supplement 1r)

Interestingly, *signaling by ERBB2* was enriched in endothelial cells (FDR=8.00%, Figure 27). The differentially expressed genes contributing to the enrichment of this pathway include FYN proto-oncogene (FYN), YES proto-oncogene (YES1), and heparin-binding EGF-like growth factor (HBEGF, Figure 27, figure supplement 1s). However, the luminal cells differentially expressed ERBB2, ERBB3, ERBB4, phosphoinositide-3-kinase regulatory subunit 1 (PIK3R1), epidermal growth factor (EGF), and EGF receptor (EGFR), which define the enrichment of other pathways associated with ERBB2 signaling, like *PI3K events in ERBB2 signaling* and *ERBB2 regulates cell motility* (Figure 27).

One other surprising finding was, despite the higher levels of estrogen receptor α mRNA in ER^Neg^ luminal cells (so named because of their undetectable protein expression and lower mRNA levels compared to the ER^Pos^ luminal cells), we found *estrogen receptor-mediated signaling* and associated pathways enriched in vascular endothelial cells (FDR = 0.008%). After investigating other members of this pathway, we found vascular endothelial cells differentially expressed estrogen receptor β (ESR2), calmodulin-1 (CALM1), and the G protein subunits GNAI2, GNG2, GNG5, GNG11, GNB1, and GNB2, and endothelial nitric oxide synthetase (NOS3) that compose this pathway. (Figure 27, Figure 27, figure supplement 1t). This is noteworthy because endothelial cells respond to estrogen for nitric oxide and prostacyclin production^82^. Our data thus indicate that the bulk of estrogen’s effects on breast endothelium likely results through ERβ signaling.

#### Lymphatic endothelial vs. Vascular endothelial cells (Pop7 vs. Pop9)

Whereas the blood’s vascular system circulates blood throughout the body, the lymphatic system collects and filters interstitial fluid, ultimately returning it to the bloodstream—a process that helps maintain fluid homeostasis. Much like capillaries of the blood system, lymphatic vessels are also formed by endothelial cells, although the two vascular architectures are distinct (Figure 2 d,f). We found lymphatic and vascular endothelial cells expressed similar markers, like CD31 (PECAM1) and von-Willebrand factor (VWF, Figure 8b) but diverge and expressed other genes that contribute to their individual functions. To explore the biological processes shaping these two endothelial cell types, we investigated their transcriptional differences.

Lymphatic endothelial cells are a rare cell type in the breast. Because of this, acquiring enough cells required for mRNA-sequencing proved challenging. However, by scaling up the number of cells stained—and through sheer brute force (∼15 hours of continuous FACS sorting for each sample), we succeeded in purifying a sufficient number of cells from two samples (24,713 cells from N228 and 29,111 cells from N239, Figure 3—table supplement 1). We detected roughly 21 million and 43 million paired end reads, respectively from these purified lymphatic endothelial cells (N228 and N239). During quality control assessment of the RNA-sequencing data, we however found outliers and conflicts between the lymphatic cell replicates (Figure 4c).

Nonetheless, the pair of lymphatic endothelial cell types clustered together well— in the hierarchical clustering dendrogram (Figure 6) and projection of principal components (Figure 7a). We thus retained the lymphatic endothelial samples for further analyses, yet, we did expect to find discordance between replicates for some genes.

When we compared transcript levels between lymphatic and vascular endothelial cell types, we identified 3,739 differentially expressed genes (Figure 28, Figure 3—table supplement 7 [7v9]). Lymphatic endothelial cells expressed the higher level of 1,462 of these, including podoplanin (PDPN), the marker we used to differentiate lymphatic cells in our FACS strategy (adj.*p-*value of 1.14E-7). We also found other lymphatic markers differentially expressed, such as ‘lymphatic vessel endothelial hyaluronan receptor1’ (LYVE1), ‘vascular endothelial growth factor receptor- 3’ (FLT4), and prospero homeobox 1 (PROX1, Figure 28). In contrast, the vascular endothelial cells differentially expressed 2,277 genes. Among these were the vascular endothelial growth factor C (VEGFC), cytokine-like 1 (CYTL1), lipocalin 1 (LCN1), endothelial selectin (SELE), and the endothelial marker CD93 (Figure 28). After ranking the expressed genes by degree of differential expression, we performed GSEA, and discovered 240 pathways enriched between the lymphatic and endothelial cell types (FDR≤10%, Pathway Analyzer). Of these enriched pathways, we’ve concentrated only on those established from genes where there was concordance between lymphatic endothelial cell replicates.

**Figure 28.**
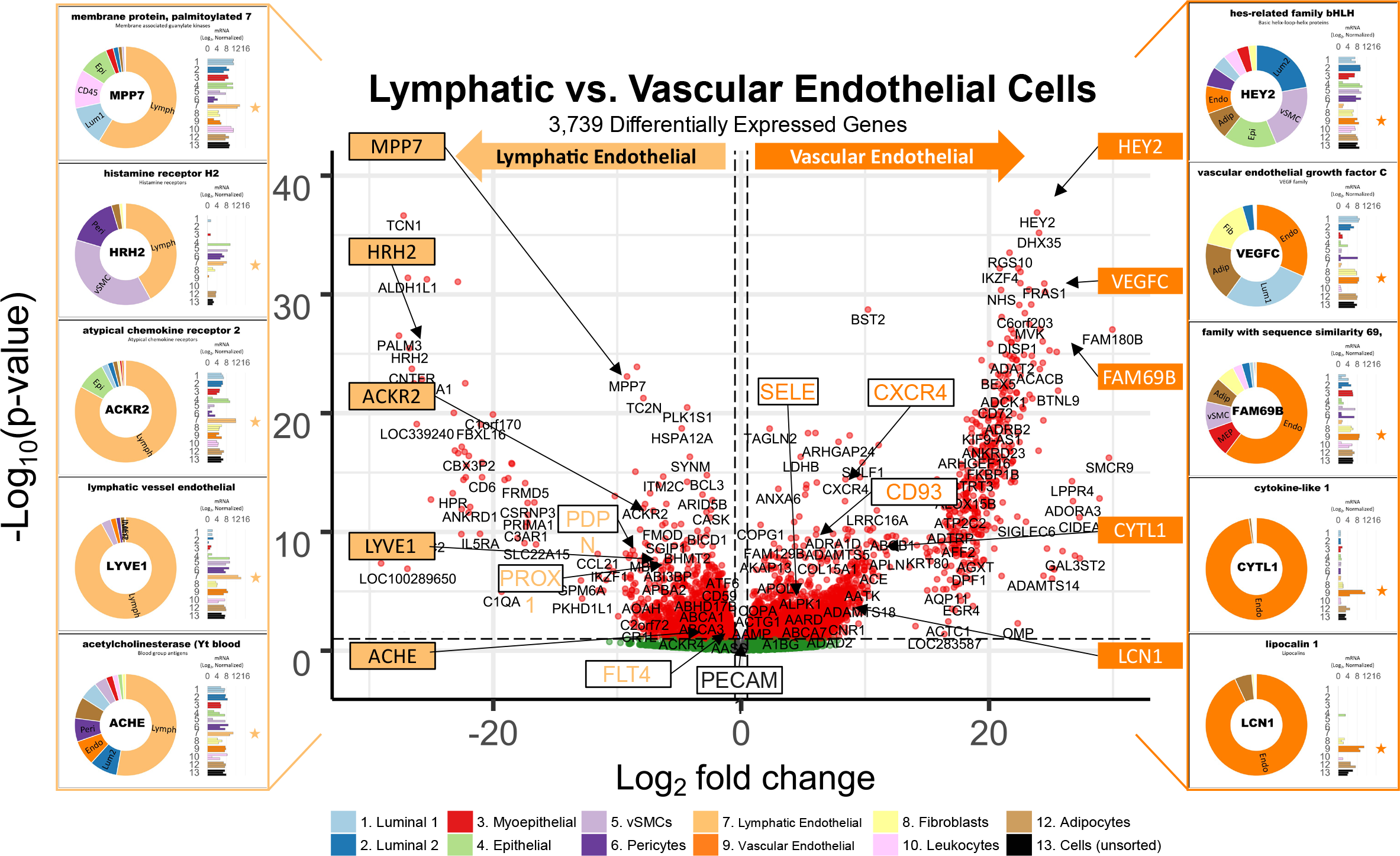
Differences between lymphatic and vascular endothelial cell types. A contrast of expressed transcripts between lymphatic and vascular endothelial cells revealed 3,739 differentially expressed genes (DeSeq2/Benjamini-Hochberg adjusted p-value ≤ 0.1). All expressed genes are displayed. The horizontal dashed lines (separating red and green dots) represent an adj. *p*-value of 0.1. Established cell type markers (of each cell type) are denoted. mRNA expression values for ten genes (indicated by filled, color-coordinated boxes) are shown at left and right of the volcano plot. Color-coded stars in the bar graphs indicate the cell type being contrasted that had the higher transcript level, i.e., lymphatic (light orange) and vascular (dark orange) endothelial cells.

#### Immune system (7v9)

Both lymphatic and blood vascular systems are essential and active components of immune responses. Lymphatic vessels serve as conduits for lymphocyte migration from peripheral tissues to the lymph nodes, making them crucial to adaptive immune response. In contrast, vascular vessels are first responders to pathogen detection and danger-signal-sensing mechanisms in the bloodstream. Indeed, when we inspected the enrichment results, both cell types were enriched with immunity-related pathways. For instance, we found *interleukin-4 and interleukin-13 signaling* pathways enriched in lymphatic endothelial cells (FDR= 1.54%). Contributing to this enrichment was the lymphatic cells’ differential expression of transforming growth factor-beta 1 (TGFB1) and IL13 receptors A1 and A2 (IL13RA1 and IL13RA2, Figure 29a). In contrast, the vascular endothelial cells were enriched with *the TCR signaling* and *interleukin-1 family signaling pathways* (FDR= 0.009% and 0.394%, Pathway Analyzer). The TCR signaling pathway’s most highly differentially expressed genes included major histocompatibility complexes HLA-DRB1 and HLA-DQA1, vasodilatory-stimulated phosphoprotein (VASP), and the TCR subunit CD247 (Figure 29b), whereas differentially expressed genes of the IL-1 signaling pathway included HMGB1, which encodes the ‘high mobility group box 1’ protein that promotes inflammatory responses to infectious signals, and protein tyrosine phosphatase 7 (PTPN7) which regulates TCR signaling (Figure 29c).

**Figure 29.**
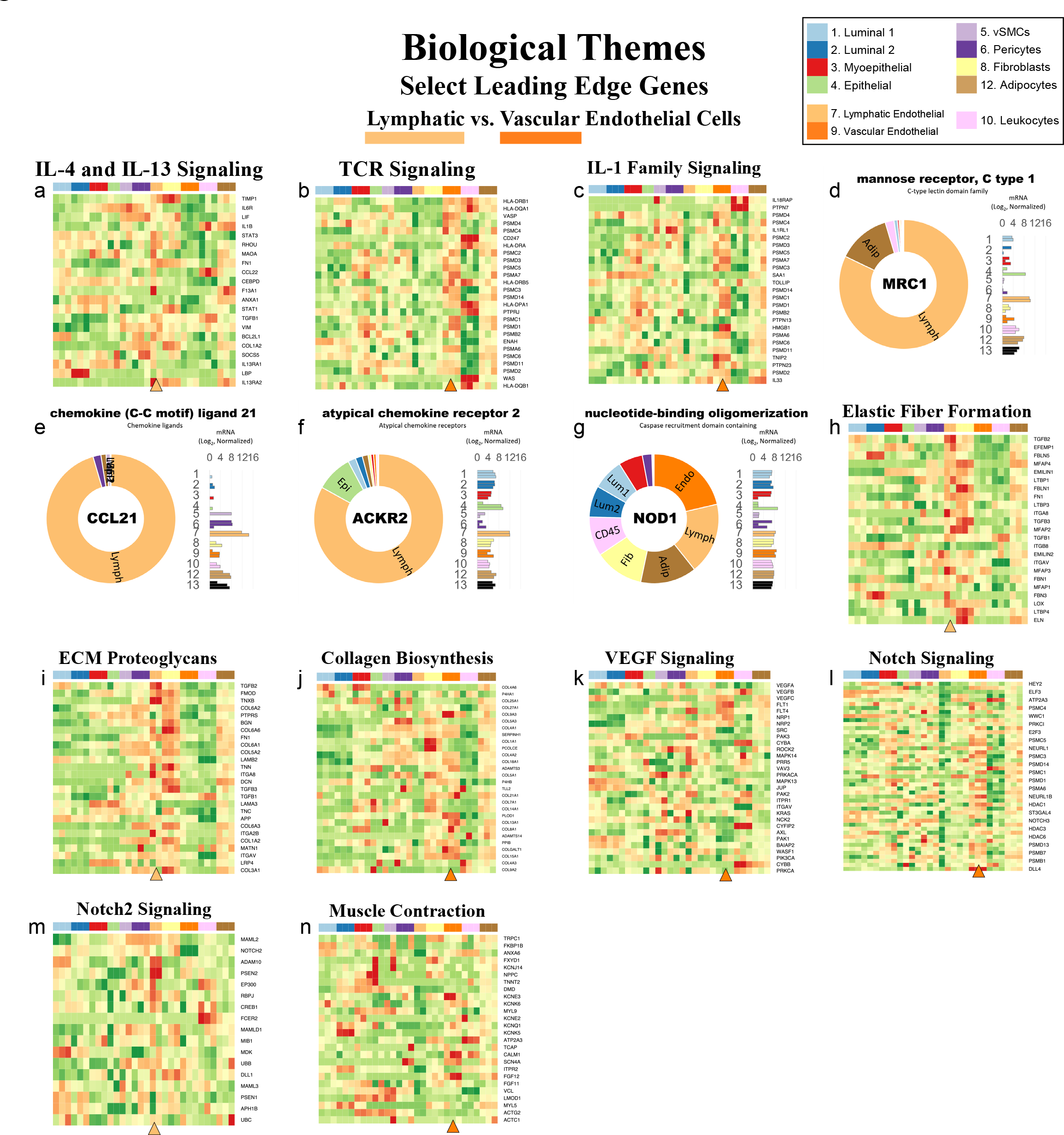
The lymphatic and vascular endothelial cells’ enriched pathways. Visualization of pathways enriched among lymphatic and vascular endothelial cell types (Enrichment Map/Cytoscape). All enriched pathways with a FDR≤0.1 appear on the map (nodes, color-coded by cell type). Pathways(nodes) are connected when they share 50% of their genes (or greater). Node size corresponds to the number of genes in the pathway, as indicated (bottom left). Select biological themes were encircled and annotated. Highly differentially expressed genes are indicated (above the theme when it coincided with the attributed cell type—and below if it was expressed by the contrasted cell type). Samples are color-coded. Due to space constraints and a large number of presented genes, their names (rows of the heatmap) must be inspected digitally (magnified in the PDF). The names of each pathway are also available in the supplemental Cytoscape file and Pathway Analyzer tool.

Other immune-related genes that we identified as being differentially expressed by lymphatic endothelial cells but were not associated with the above pathways included MRC1, TLR4, CCL21, and ACKR2 (Figure 29—figure supplement d-f, Figure 23- figure supplement 3g).

MRC1 encodes mannose receptor 1, a cognate binding partner of L-selectin—that is expressed by lymphocytes and facilitates their binding to endothelial cells. Toll-like receptor 4 (TLR4) plays a crucial role in mediating the innate immune response to bacterial lipopolysaccharide (LPS), where it engages other levers of the immune response upon binding to bacterial LPS^83^. chemokine ligand 21 (CCL21) mediates homing of lymphocytes to lymph organs, whereas the atypical chemokine receptor 2 (ACKR2) regulates cellular interactions with inflammatory leukocytes^84^. Interestingly, we found lymphatic and vascular endothelial cells both expressed the pattern recognition receptor NOD1, which detects bacterial peptidoglycan fragments and initiates immune responses to gram-negative bacteria^85^ (Figure 29g). There is thus ample evidence showing the respective endothelial cell types are interacting with cells of the immune system and actively participating in immune surveillance –in distince ways.

#### ECM Organization (7v9)

While vascular endothelial cells are surrounded by a continuous basement membrane, the basement membrane surrounding lymphatic endothelial cells is thin and discontinuous (Figure 2d, Figure 2—figure supplement 2)^8^. We thus expected to find differences in their respective expression of extracellular matrix components. After performing pathway analysis, we indeed found enrichment of *ECM proteoglycans* in the lymphatic cells (FDR=1.09%). Contributing to the pathway’s enrichment was the strong lymphatic expression of all three TGFβ ligands (TGFB1, -B2 and -B3), fibromodulin (FMOD), biglycan (BGN), fibronectin (FN1), tenascin XB (TNXB), integrin alpha 8 (ITGA8), collagen 6A2 (COL6A2), and laminin beta 2 (LAMB2, Figure 29h). Elastic fibers appear to be strongly associated with lymphatics, as we found the *elastic fiber formation* pathway highly enriched in this cell type (FDR=0.5%). Twenty-four genes in the elastic fiber pathway were elevated in lymphatic endothelial cells, including ‘elastin microfibril interfacer 1’ (EMILIN1) and latent transforming growth factor beta (LTBP1). In a prior section, we introduced the fibulin gene family, where we discussed their importance to elastic fibers and identified several family members (EFEMP1, FBLN1, FBLN5) differentially expressed by lymphatic endothelial cells –these genes are also members of the *elastic fiber formation* pathway (Figure 29i).

The only ECM-related pathway enriched in vascular endothelial cells was c*ollagen biosynthesis and modifying enzymes* (FDR = 6.65%). Differentially expressed genes contributing to its enrichment included the collagens COL9A3, COL4A1, COL4A2, COL18A1, COL21A1, COL13A1, COL8A1, COL15A1; procollagen-lysine 2-oxoglutarate 5-dioxygenase 1 (PLOD1); and collagen beta (1-O) galactosyltransferase 1 (COLGALT1, Figure 29j). The type IV collagens are a major structural component of basement membranes, indicating that endothelial cells collaborate with pericytes to produce the vascular basement membrane.

#### Cell signaling (7v9)

VEGF signaling is a critical regulator of vascular development. Interestingly, we found *signaling by VEGF* enriched in lymphatic endothelial cells (FDR= 0.563%) due to their elevated expression of 55 of the 105 genes in the pathway, despite the vascular endothelial cells’ differential expression of several key pathway members, like vascular endothelial growth factor c (VEGFC), placental growth factor (PGF), and VEGF Receptor-1 (VEGFR1/FLT1). The lymphatic endothelial cells, in contrast, expressed higher levels of other pathway members, like: ‘p21 (Cdc42/Rac)-activating protein 3’ (PAK3), cytochrome b245 (CYBA), the mTORC2 target PRR5, and VEGFR3 (FLT4, Figure 29k).

Distinct Notch signaling pathways were enriched in both endothelial cell types. We found the broader *Signaling by notch* pathway enriched in vascular endothelial cells (FDR=0.434%), whereas the *signaling by notch2* pathway was enriched in lymphatic endothelial cells—at a false discovery rate of 9.97% (near the 10% cut-off). Of the 174 genes in the *signaling by notch* pathway, 103 were differentially expressed by vascular endothelial cells, including genes encoding the Notch ligands jagged 1 (JAG1), jagged 2 (JAG2), and delta-like 4 (DLL4, Figure 29l). Notch2 transcript levels were higher in the lymphatic cells, which, compared to all other breast cell types, appeared to stem from its repression within vascular endothelial cells (Figure 29m). This contributed to *signaling by notch2* pathway’s enrichment in lymphatics, along with robust lymphatic expression of other pathway members, like ADAM metallopeptidase domain 10 (ADAM10) and mastermind-like transcription coactivator (MAML2).

#### Muscle contraction (7v9)

We found the *muscle contraction* pathway enriched in vascular endothelial cells (FDR= 4.30%, Pathway Analyzer). However, both endothelial cell types did not appreciably express skeletal actin (ACTA1) or smooth muscle actins (ACTA2 or ACTG2), so they do not appear to be contractile. Of course, this is consistent with contractile smooth muscle cells being the primary regulator of vessel diameter. Instead, we found the vascular endothelial cells’ enrichment of the *muscle contraction* pathway primarily resulted from the differential expression of a) the calcium-permeable transient receptor potential cation channel (TRPC1), b) the calcium-dependent membrane protein annexin A6 (ANXA6), and c) several potassium channels, including KCNE3, KCNK5, and KCNK6. These intermediate and small conductance channels play an essential role by regulating the release of endothelial cell autacoids, like nitric oxide^86^, and were noticeably absent in lymphatic endothelial cells (Figure 29n).

#### Fibroblasts (ADMSCs) vs. Adipocytes (Pop8 vs. Pop12)

As epithelial cells form the parenchyma of the breast, the bulk of the tissue stroma is formed by the mesenchymal cell types (adipocytes, fibroblasts, vascular smooth muscle cells, and pericytes), with endothelial cells and a spectrum of leukocytes providing key elements^8^. We did not quantify adipocytes by FACS, but we could deduce from the RNA-sequencing data that adipocytes are one of, if not the most, abundant breast cell type. This was inferred from the covariance analysis and hierarchical clustering that grouped adipocytes with the whole tissue control samples, indicating they were disproportionately comprised of adipocytes (Figure 4b and Figure 6). The remaining cell types were assessed by FACS, and it came as no surprise that fibroblasts were also abundant (Figure 1b).

Fibroblasts are spindle-shaped cells that predominantly provides the tissue’s foundation—a collagenous-rich matrix of fibrillar extracellular proteins, which fibroblasts shape through the secretion of growth factors, proteoglycans, and matrix-bound proteases. Fibroblasts are the archetypical mesenchymal cell type, existing within a developmental lineage that is shared with adipocytes. In our adjoining article, we described how breast fibroblasts express mesenchymal stem cell markers (CD10, CD34, CD73, Thy1, CD44, CD105) and displayed a functional multi- lineage potential when cultured under osteogenic or adipogenic conditions^8^. For these reasons, we designated Pop8 fibroblasts as adipocyte-derived mesenchymal stem cells (ADMSCs) but commonly refer to them as fibroblasts. In our cultures, and especially within the tissue, adipocytes appeared as large, spherical cells containing a single large lipid droplet (unilocular adipocytes) that stained for neutral lipids^8^ (Figure 2a,c; Figure 2—figure supplement 1a, 2a). Adipocytes notably serve as the body’s chief storage depot of triglycerides and other acylglycerols and regulate energy metabolism through secretion of adipokines^87^, e.g, leptin hormone (LEP) and adipsin (CFD), which we found highly expressed in the pop12 lipid (adipocyte) samples (Figure 31a,b). To establish the different cellular processes shaping adipocytes and fibroblasts, we examined their expressed transcripts.

In contrast to the other breast cell types, freshly-isolated adipocytes were not FACS purified prior to RNA-sequencing, and therefore results should be interpreted differently. Adipocytes were instead isolated from the phase-separated lipid layer of the collagenase tissues digests, which we had visually inspected for contaminating cellular content (Hoechst staining). The number of paired-end mRNA fragments measured from the RNA isolated from the lipid layers was excellent (34 to 45 million fragments), and the replicates nicely clustered together (Figure 4b; Figure 6; Figure 7—figure supplement 2a, Figure 8a). However, one sample did stray in the projection of PC1 and PC2 (Figure 7a), indicating some discordance between samples. We also noticed, in all three adipocyte replicates, low transcript levels for many genes (e.g., this can be observed in the many heatmaps presented herein, such as those containing the claudins and gap junction proteins, Figures 16,17). These adipocyte samples thus have a higher noise level than that observed in the FACS-purified samples, which is to be expected given the purity and excellent quality of those samples. The noise is likely derived from a fraction of contaminating cells in the samples, such as leukocytes. Therefore, we have prioritized genes and pathways in our analyses that are not heavily associated with immune response or where there is not evidence of robust adipocyte mRNA levels. With the caveat that the adipocyte samples should be interpreted differently, we explored the nature of adipocytes by comparing them with fibroblasts.

Pairwise comparison of RNA transcripts from adipocytes and fibroblasts revealed 1,171 differentially expressed genes. This was notably the lowest number of differentially expressed genes between any cell-type comparison, similar to the number differentially expressed between the two luminal epithelial cell types (1,301, figure 20). We anticipated fewer DE genes given the similarity and shared lineage of adipocytes and fibroblasts. Of the 1,171 differentially expressed genes we identified, 207 were more highly expressed by fibroblasts, and 964 were more highly expressed by adipocytes (Figure 30, Figure 3—table supplement 7 [8v12]).

**Figure 30.**
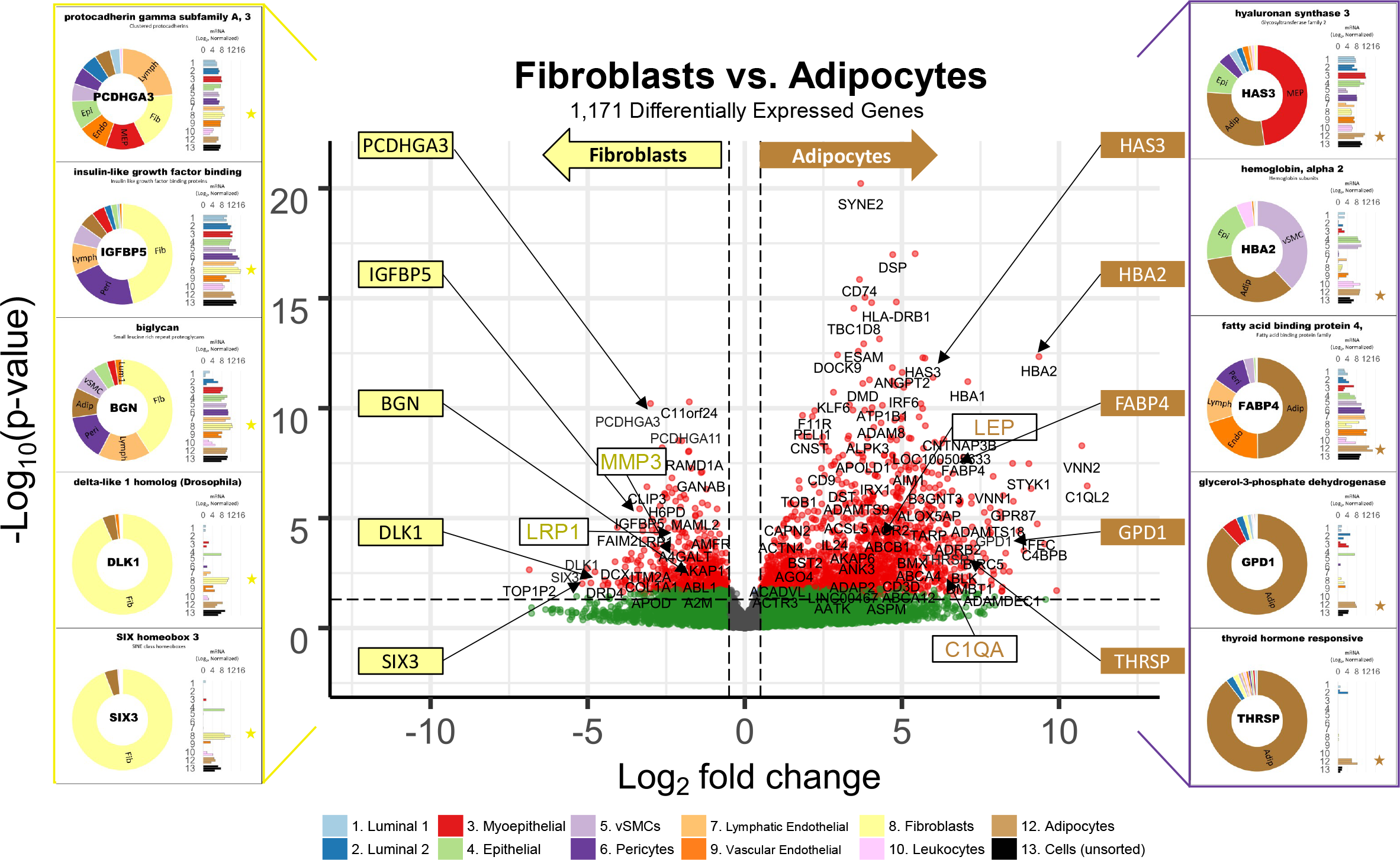
Differences between fibroblasts and adipocytes. A contrast of expressed transcripts between fibroblasts and adipocytes revealed 1,171 differentially expressed genes (DeSeq2/Benjamini-Hochberg adjusted p-value ≤ 0.1). All expressed genes are displayed. The horizontal dashed lines (separating red and green dots) represent an adj. *p*-value of 0.1. Established cell type markers (of each cell type) are denoted. mRNA expression values for ten genes (indicated by filled, color-coordinated boxes) are shown at left and right of the volcano plot. Color-coded stars in the bar graphs identify the position of the fibroblasts (yellow) and adipocytes (brown).

**Figure 31.**
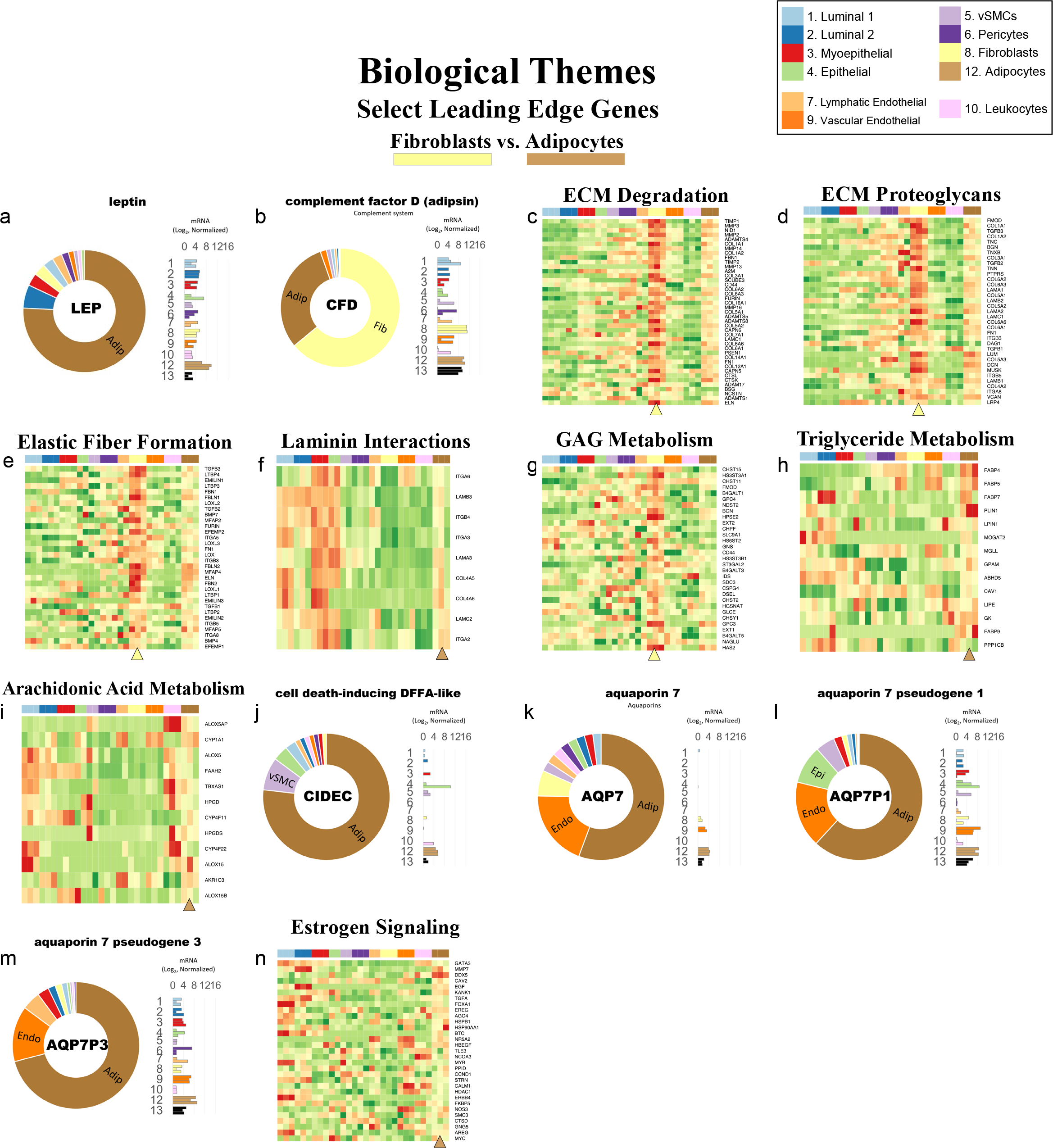
The fibroblasts and adipocytes’ enriched pathways. Visualization of pathways enriched among fibroblasts and adipocytes (Enrichment Map/Cytoscape). All enriched pathways with a FDR≤0.1 appear on the map (nodes, color-coded by cell type). Pathways(nodes) are connected when they share 50% of their genes (or greater). Node size corresponds to the number of genes in the pathway, as indicated (bottom left). Select biological themes were encircled and annotated. Highly differentially expressed genes are indicated (above the theme when it coincided with the attributed cell type—and below if it was expressed by the contrasted cell type). The names of each pathway are available in the supplemental Cytoscape file and Pathway Analyzer tool. Samples are color-coded. Due to space constraints and a large number of presented genes, their names (rows of the heatmap) must be inspected digitally (magnified in the PDF).

When we investigated the identity of the differentially expressed genes, we found fibroblasts expressed the ECM protein biglycan, delta-like non-canonical Notch ligand 1 (DLK1), matrix metalloproteinase 3 (MMP3), and others. In contrast, adipocytes differentially expressed leptin (LEP), thyroid hormone-responsive protein (THRSP), and glycerol-3-phosphate dehydrogenase (GPD1), along with 961 other genes (Figure 30). After performing GSEA, we identified 387 pathways enriched between the two cell types (FDR ≤ 20%), with 69 pathways in fibroblasts and 318 in adipocytes (Pathway Analyzer). Due to the large number of of immune-related genes and pathways in adipocyte samples, we increased the false discovery rate cutoff to 20% for this comparison, which resulted in the enrichment of more pathways related to fat storage and metabolism.

#### Extracellular matrix organization (8v12)

Twelve pathways associated with ECM organization were enriched in fibroblasts, including *degradation of the extracellular matrix*, *ECM proteoglycans*, and *elastic fiber formation* (FDR< 0.001%, < 0.001%, and =0.090%, respectively, Pathway Analyzer). Fibroblasts differentially expressed genes contributing to these pathways, including TIMP metallopeptidase inhibitor 1 and 2 (TIMP1 and TIMP2); metallopeptidases 2, 3, 13, 14, and 16 (MMP2, MMP3, MMP13, MMP14, and MMP16); fibulin- 1 (FN1); fibrillin-1 (FBN1); and elastin (ELN, Figure 31c-e). There were two pathways associated with ECM organization enriched in adipocytes, including *laminin interactions* and *anchoring fibril formation* (FDR=3.23% and 9.97%, Figure 31f). Adipocytes differentially expressed integrins A6, B4, A3, and A2; laminins B3, A3, and C3; and collagens 4A5 and 4A6, which contributed to the enrichment of these pathways (Figure 31f).

#### Metabolism (8v12)

Included in the enriched metabolic pathways was *Glycosaminoglycan metabolism* that was associated with fibroblasts (FDR<0.001%). This pathway’s genes included heparanase-2 (HPSE2), hyaluronan synthase-2 (HAS2), and glypican-3 (GPC3, Figure 31g). In contrast, we found adipocytes enriched in pathways associated with lipid metabolism, that included *triglyceride metabolism, triglyceride catabolism*, and *arachidonic acid metabolism* (FDR= 15.5%, 15.5%, and 4.13%, Pathway Analyzer, Figure 31h). Differential expression of fatty acid binding proteins (FABP4, FABP5, FABP7, and FABP9) were among those genes contributing to enrichment of pathways associated with triglyceride metabolism and catabolism (Figure 31h). These intracellular fatty acid-binding proteins (FABPs) exhibit high affinity for a single long-chain fatty acids and are essential for triglyceride storage; they mediate intracellular fatty acid transport and interact with PPAR nuclear receptors^88^. Enrichment of *arachidonic acid metabolism* in adipocytes was defined by differential expression of arachidonate 5-lipoxygenase (ALOX5) and its activating protein (ALOX5AP, Figure 31i), which catalyzes the first two steps of the biosynthesis of leukotrienes, which are potent mediators of inflammation.

We also found adipocytes uniquely expressed cell death inducing DFFA like effector C (CIDEC, Figure 31j), which binds to lipid droplets and regulates their enlargement, thereby restricting lipolysis and favoring storage^89^. The adipocytes also uniquely expressed aquaporin AQP7 and the aquaporin pseudogenes AQP7P1 and AQP7P3 (Figure 31k-m), which control water and glycerol transport and promote the catabolism of lipids in adipocytes^90^.

#### Cell signaling (8v12)

Accumulation of adipose tissue is regulated in part by estrogen signaling^91^. Therefore, it is unsurprising that *ESR-mediated signaling* and its associated pathways were enriched in adipocytes (FDR= 0.110%), which differentially express the contributing genes GATA binding protein 3 (GATA3), caveolin 2 (CAV2), epidermal growth factor (EGF), and epiregulin (EREG, Figure 31n). Adipocytes also expressed estrogen receptors 1 and 2 (ESR1 and ESR2), albeit at 10x lower levels lower than ER^Pos^ luminal epithelial cells but at a 4.3x higher level than fibroblasts (Figure31n, Pathway Analyzer).

#### Pericytes vs. Fibroblasts (ADMSCs, Pop6 vs. Pop8)

Whereas mesenchymal adipocytes contribute to the storage and mobilization of energy-rich metabolites; and mesenchymal fibroblasts secrete ECM proteins and enzymes that shape the extracellular stromal environment, the mesenchymal perivascular cells influence and modulate blood vessels. In a companion article^8^, we described how we identified vascular smooth muscle cells and pericytes while devising methods to isolate and culture all breast cell types, finding these perivascular cells to be an elusive set of cell types. Pericytes have long been overlooked^92, 93^, largely stemming from their heterogeneity that, in turn, required a combinatoric assessment of markers to identify and purify them. After discovering how to reliably isolate and propagate breast pericytes, we found their microscopic appearance fascinating, as they had a stellate astrocyte-like morphology that was entirely distinct from fibroblasts and adipocytes^8^.

However, it was difficult to tell the pericytes and fibroblasts apart at higher confluency. We were thus interested in exploring the biological processes defining these two similar mesenchymal cell types (i.e., FACS populations #6-pericytes and #8-ADMSC/fibroblasts).

Analysis of transcripts expressed by freshly isolated pericytes and fibroblasts revealed 2,932 differentially expressed genes (BH/FDR adj. *p*-value < 0.1, Figure 32). Interestingly, these were not split evenly between the two cell types, as fibroblasts had roughly three times the number of DE genes than pericytes. Among these 2,187 genes, were platelet-derived growth factor receptor alpha (PDGFRA), matrix metallopeptidase 2 (MMP2), and hydroxyacid oxidase 2 (HAO2). In contrast, pericytes differentially expressed 748 genes, including pyruvate dehydrogenase kinase 4 (PDK4), CD36, and desmin (DES, Figure 32). Along with desmin, we found the pericytes differentially expressed other genes previously associated with pericytes, such as melanoma cell adhesion molecule (MCAM), G protein receptor RGS5, and the potassium channel KCNJ8. We also found pericytes uniquely expressed the serotonin receptor isoform HTR1F, which was distinct from the HTR1B isoform we discovered uniquely expressed by vascular smooth muscle cells that, coincidently, helped us identify the vSMC FACS population^8^ (serotonin is a vasoactive peptide^94^). We ranked all expressed genes by their degree of differential expression and used this list for pathway analysis.

**Figure 32.**
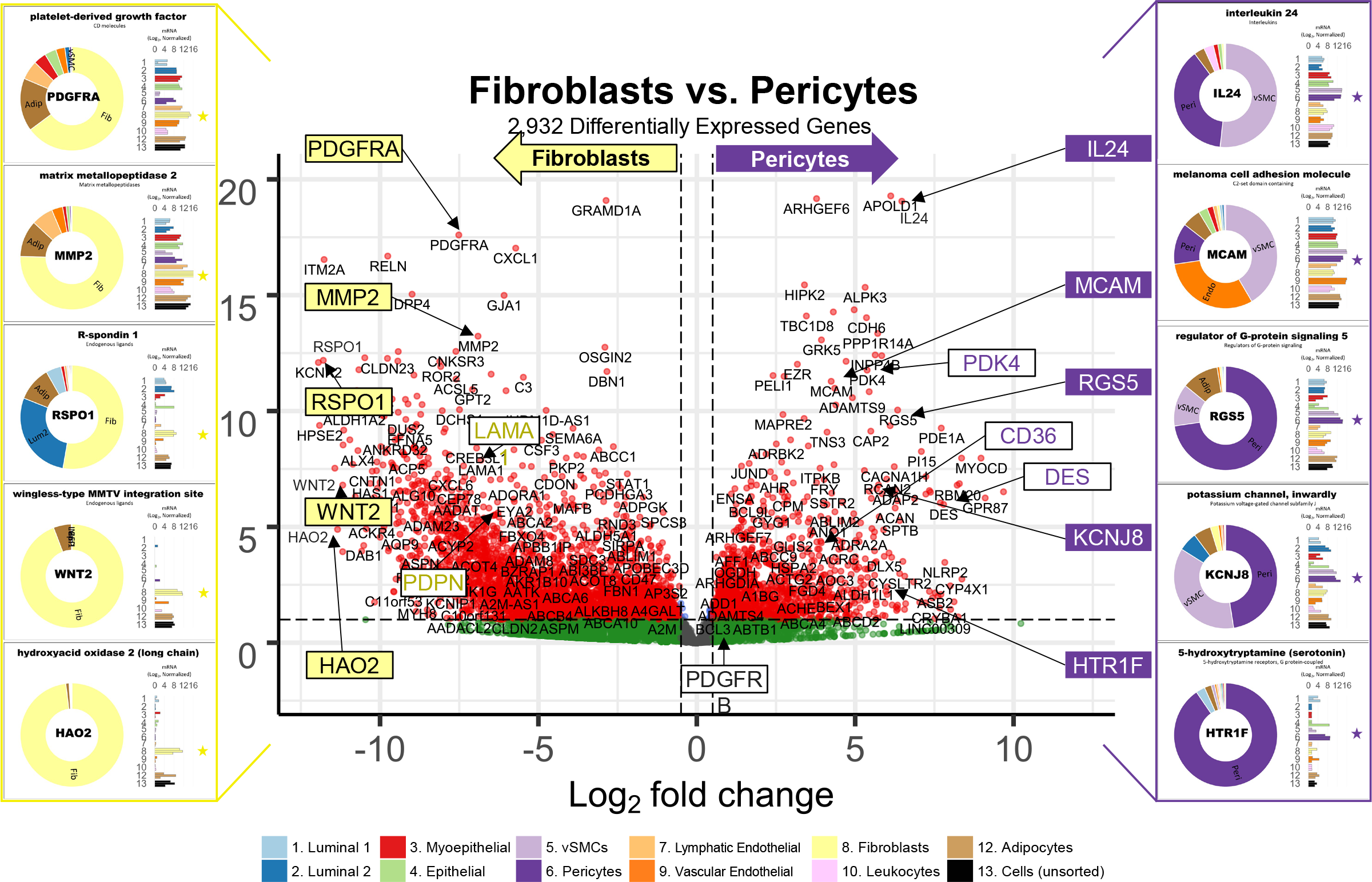
Differences between pericytes and fibroblasts. A contrast of expressed transcripts between pericytes and fibroblasts revealed 2,932 differentially expressed genes (DeSeq2/Benjamini-Hochberg adjusted p-value ≤ 0.1). All expressed genes are displayed. The horizontal dashed lines (separating red and green dots) represent an adj. *p*-value of 0.1. Established cell type markers (of each cell type) are denoted. mRNA expression values for ten genes (indicated by filled, color-coordinated boxes) are shown at the left and right of the volcano plot. Color-coded stars in the bar graphs identify the position of the pericytes (purple) and fibroblasts (yellow).

We discovered 477 pathways enriched between pericytes and fibroblasts at a false discovery rate of 10%. We found the enrichment of many pathways resulted from small concordant expression differences between the cell types, such as those involved in transcriptional regulation, cell cycle, and immunity. Here, we focus on pathways containing significantly differentially expressed genes.

#### Muscle Contraction (6v8)

As one would predict, we found perivascular pericytes enriched with pathways involved with *muscle contraction* and *smooth muscle contraction* (FDR =0.361% and 0.026%, respectively, Figure 33). Pericytes differentially expressed myosin light chain kinase (MYLK), myosin heavy chain 11 (MYH11), actin gamma 2 (ACTG2), and the voltage-gated sodium channel type IV (SCN4A). Also differentially expressed by pericytes was dystrophin (DMD), which anchors ECM proteins to cytoplasmic filamentous actin (and whose defects lead to Duchenne and Becker muscular dystrophies). Tropomodulin 1 (TMOD1) was also highly expressed by pericytes, which is a protein that regulates tropomyosin via blocking the elongation and depolymerization of actin filaments (Figure 33- figure supplement 1a). Interestingly, *cardiac conduction* was enriched in fibroblasts (FDR=5.64%), which differentially expressed T-box 5 (TBX5) transcription factor; potassium channels KCNK1, KCNK2, KCNIP1, and KCNIP4; and the sodium channel SCN1B (Figure 33, Figure 33- figure supplement 1b). Fibroblasts share similar expression of voltage-gated channels to cardiac myocytes, which are responsible for cardiac conduction in the heart. Additionally, the dramatic difference in gene expression could be due to the low expression of many of these channels in pericytes, which may instead rely on endothelial cells for depolarization signals (as we found similar channels were differentially expressed by vascular endothelial cells when compared to their lymphatic counterparts, Figure 29n).

**Figure 33.**
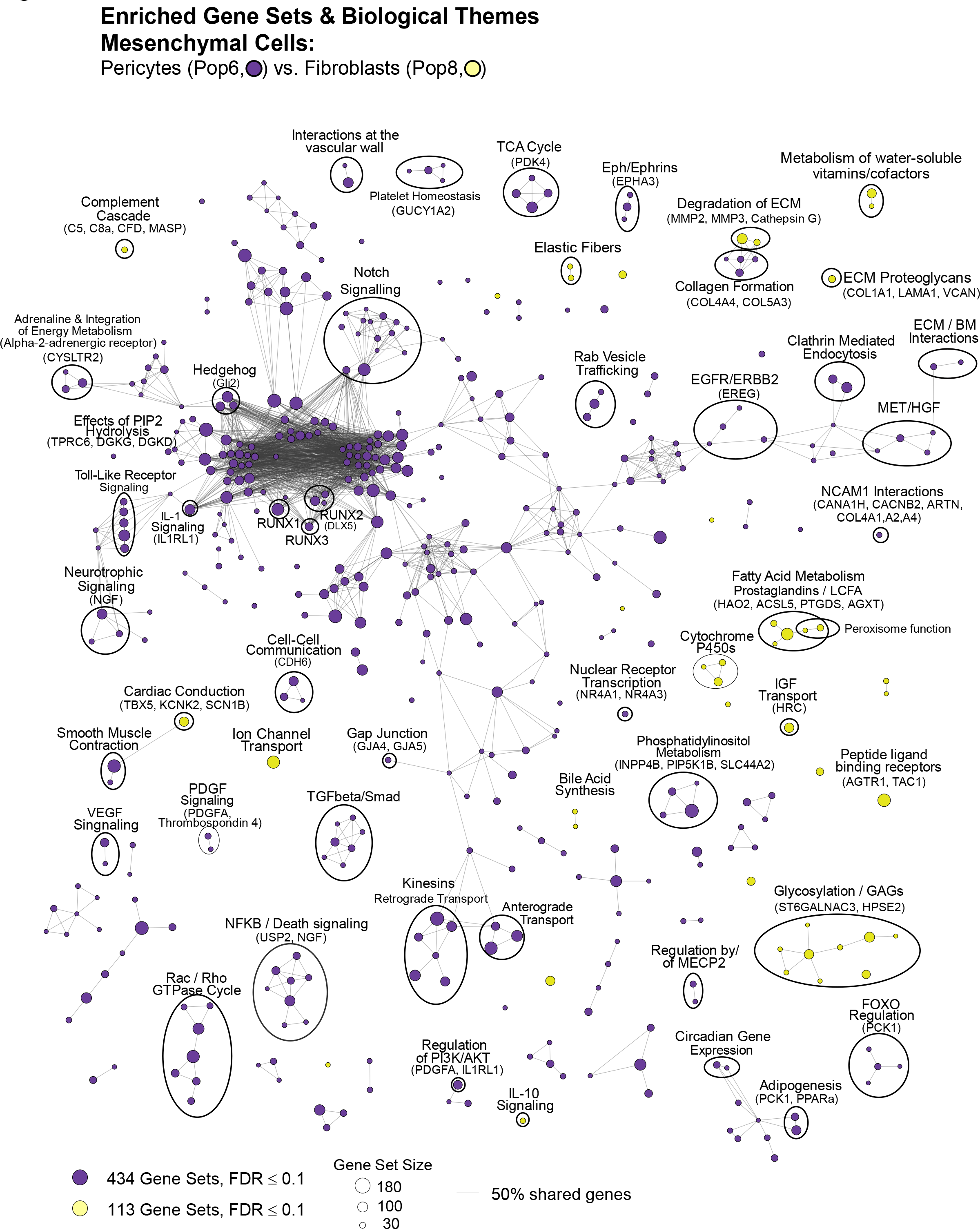
The pericytes and fibroblasts’ enriched pathways. Visualization of pathways enriched among pericytes and fibroblasts (Enrichment Map/Cytoscape). All enriched pathways with a FDR≤0.1 appear on the map (nodes, color-coded by cell type). Pathways(nodes) are connected when they share 50% of their genes (or greater). Node size corresponds to the number of genes in the pathway, as indicated (bottom left). Select biological themes were encircled and annotated. Highly differentially expressed genes are indicated (above the theme when it coincided with the attributed cell type—and below if it was expressed by the contrasted cell type). Samples are color-coded. Due to space constraints and a large number of presented genes, their names (rows of the heatmap) must be inspected digitally (magnified in the PDF). The names of each pathway are also available in the supplemental Cytoscape file and Pathway Analyzer tool.

#### Hemostasis (6v8)

We found fourteen pathways associated with hemostasis enriched in pericytes, including *platelet homeostasis* (FDR= 0.732%, Figure 33). Its enrichment was defined by differential expression of phosphodiesterases 1A, 1B, and 5A (PDE1A, PDE1B, and PDE5A), transient receptor cation channel C6 (TRPC6), cGMP-dependent protein kinase 1 (PRKG1), and the prostaglandin receptor PTGIR (Figure 33- figure supplement 1c). The phosphodiesterases are responsible for hydrolyzing cGMP and modulating the effects of nitric oxide signaling. Also related to the vasculature was the enrichment, in pericytes, of *cell surface interactions at the vascular wall* (FDR=1.14%), resulting from their differential expression of MER proto-oncogene (MER), thrombomodulin (THBD), and angiopoietin 1 (ANGTP1, Figure 33, Figure 33- figure supplement 1d).

#### Intracellular Transport (6v8)

Pericytes differentially expressed kinesins (KIF4A, KIF2A, and KIF26B) and tubulins (TUBB2A, 3, and 4B), leading to their enrichment of *kinesins* pathway (FDR<0.001%, Figure 33, Figure 33- figure supplement 1e). *Clathrin-mediated endocytosis* was also enriched in pericytes (FDR<0.001%), due to their differential expression of the endocytic adapter protein SGIP1, WNT5A ligand, the wnt receptor ‘frizzled class receptor 4’ (FZD4), angiotensin II receptor (AGTR1), and the endocytosis inhibitor synaptotagmin XI (SYT11, Figure 33, Figure 33- figure supplement 1f). This pathway regulates the cell surface expression and signaling of RTKs and GPCRs^95, 96^, and we found that most of the enriched pathways associated with RTKs or GPCRs were enriched in pericytes (Pathway Analyzer).

#### Cell signaling (6v8)

We identified thirty enriched pathways associated with RTK signaling in pericytes, including *signaling by NTRKs*, *signaling by VEGF*, and *signaling by PDGF* (FDR=0.011%, 0.365%, and 6.67%, respectively, Figure 33). Neurotropic tyrosine kinase receptors are activated by neurotrophins like NGF and NTF3—which were highly and differentially expressed by pericytes (Figure 33- figure supplement 1g). Pericytes also differentially express AXL receptor tyrosine kinase (AXL), caveolin-1 (CAV1), and integrin beta 3 (ITGB3), which define the enrichment of *signaling by VEGF* (Figure 33- figure supplement 1h)*. Signaling by PDGF* was enriched in pericytes due to the differential expression of collagens COL4A1, COL4A2, COL4A4, COL5A3; platelet-derived growth factor alpha (PDGFA); and thrombospondin 4 (THBS4, Figure 33- figure supplement 1i).

The enrichment of thirteen Notch signaling pathways in pericytes indicate their importance to pericyte function. The broader *signaling by notch* pathway (FDR<0.001%) was enriched largely because of the differential expression of NOTCH receptors 1 and 3 (NOTCH1, NOTCH3), the downstream transcription factors HEYL and HEY2, and SMAD family member 3 (SMAD3, Figure 33- figure supplement 1s).

#### Cell-Cell Communication (6v8)

Communication between pericytes and endothelial cells is critical to vessel development and function. We found pericytes enriched with the broad *cell-cell communication* and cell junction pathways (FDR=0.10% and 0.06%, respectively, Figure 33). These pathways involve tight junctions, adherins junctions, gap junctions, and signal regulatory proteins, and pericytes differentially expressed a) cadherins 3 and 6 (CDH3 and CDH6), classical calcium adhesion molecules; b) plectin (PLEC), which interacts and organizes connections to intermediate filaments and microtubules and are essential components of desmosomes and hemidesmosomes; and c) laminin subunit alpha-3 (LAMA3), which assembles into laminin heterotrimers that are essential components of the vascular basement membrane. (Figure 33- figure supplement 1j).

#### Extracellular Matrix Organization (6v8)

Pericytes and fibroblasts have different functional roles requiring distinct interactions with different ECM components. This was reflected in the different ECM organization pathways we found in either cell type. For example, acting in opposition, the *collagen formation* pathway was enriched in pericytes, whereas the *collagen degradation* and *degradation of the extracellular matrix* pathways were enriched in fibroblasts. Enrichment of the *Collagen formation* pathway in pericytes (FDR=4.28%) resulted from their differential expression of COL4A1, COL4A2, COL4A3, COL4A4, COL4A5, COL5A3, COL17A1, COL23A1, COL25A1, and COL27A1 (Figure 33, Figure 33- figure supplement 1k). As introduced in the discussion of the collagen family above, the perivascular cells expressed the highest levels of type IV collagens (basement membrane collagens, Figure 15—figure supplement 2a-c), consistent with their spatial location in the tissues—embedded within the vascular basement membranes—and role of maintaining these basement membranes. In contrast, fibroblasts exist in the extracellular matrix, which they are responsible for forming and shaping. We found fibroblasts differentially expressed collagens COL1A1, COL3A1, COL6A6, COL8A1, COL8A2, COL14A1, and COL16A1, contributing to the *degradation of the extracellular matrix* pathway’s enrichment (FDR=0.705%, Figure 33, Figure 33- figure supplement 1l). Additional differentially expressed genes included matrix metalloproteinases MMP2, MMP3, MMP10, MMP12, MMP13, and MMP16; TIMP metallopeptidase inhibitor 1 (TIMP1); calpains 5 and 6 (CAPN 5 and CAPN6); and the bone remodeling protein cathepsin K—which cleaves collagen I fibers (Figure 33- figure supplement 1l). We also found enrichment of *ECM proteoglycans* in fibroblasts (FDR=4.68%), as they differentially expressed versican (VCAN), asporin (ASPN), lumican (LUM), heparan sulfate proteoglycan 2 (HSPG2), decorin (DCN), and biglycan (BGN) proteoglycans (Figure 33, Figure 33- figure supplement 1m). Other genes in this pathway highly expressed by fibroblasts were tenascins N and C (TNN, TNC), TGFB3, and laminin alpha 1 (LAMA1, Figure 33- figure supplement 1m).

#### Elastic Fibers (6v8)

Elastic fibers are essential components of connective tissue and contribute to tissue elasticity. We discovered pathways related to elastic fibers were among the most significantly enriched pathways in fibroblasts, i.e., *molecules associated with elastic fibers* (FDR=0.29%) and *elastic fiber formation* (FDR=0.90%). Leading to these pathways’ enrichment were the fibroblasts’ robust expression of elastin (ELN), elastic microfibril interfacer 1 (EMILIN1), fibrillins 1 and 2 (FBN1 and FBN2), bone morphogenic protein 7 (BMP7), fibronectin (FN1), and lysyl oxidase 1 (LOXL1, Figure 33, Figure 33- figure supplement 1n).

#### Metabolism (6v8)

Pericytes and fibroblasts were enriched with different metabolic pathways. There were fifteen pathways enriched in pericytes and thirteen in fibroblasts (Pathway analyzer).

One of the most significant pathways enriched in pericytes was *Integration of energy metabolism* (FDR=0.016%), stemming from the pericytes’ differential expression of adrenoreceptor alpha 2A and 2C (ADRA2A and ADRA2C), thrombospondin receptor (CD36), adenylate cyclases 6 and 9 (ADCY6 and ADCY9), cysteinyl leukotriene receptor 2 (CYSLTR2), and the voltage-dependent calcium channel CACNB2 (Figure 33, Figure 33- figure supplement 1o). In contrast, fibroblasts were enriched in *fatty acid metabolism* (FDR=0.889%), owing to their differential expression of prostaglandin synthases PTGES, PTGS2, PTGIS, and PTGDS; the C2 domain-containing phospholipase PLA2G4A; and their unique expression of hydroxyacid oxidase 2 (HAO2, Figure 33, Figure 33—figure supplement 1p).

#### White adipocyte differentiation (6v8)

We found forty-five of the eighty-four genes composing the *transcriptional regulation of white adipocyte differentiation* pathway were elevated in pericytes, leading to its enrichment (FDR=0.007% Figure 33). Genes differentially expressed by pericytes included NR2F2, which regulates apolipoprotein A-I transcription; CD36 (fatty acid translocase), which regulates the transport and uptake of long-chain fatty acids, and fatty acid binding protein-4 (FABP4), which mediates intracellular fatty acid transport (Figure 33—figure supplement 1q). One of the most notable genes in this pathway is phosphoenolpyruvate carboxykinase 1 (PCK1), which catalyzes the production of phosphoenol pyruvate from oxaloacetate. The activity of PCK1, which is regulated at the transcriptional level, commits mitochondrial pyruvate to gluconeogenesis or glyceroneogenesis^97^. This step is irreversible under physiological conditions and is the rate-limiting step controlling gluconeogenesis flux. We found PCK1 expression in breast cells was limited to the pericytes and adipocytes (Figure 33— figure supplement 1r), indicating, at the very least, that pericytes have an increased demand for cytoplasmic pyruvate.

## Summary

Resolving the types and properties of cells in a tissue is vital to understanding the underlying symphony of interactions that make life possible. It is essential to overcome diseases such as cancer. Here, and in the adjoining article, we explored the fabric of the breast through a cytometric and molecular dissection of the tissue, which has provided us with insights into its inner workings. From the outset, we had questions regarding the number and types of cells composing the breast, their spatial location, their functions, and how they contributed to the tissue as a whole.

To address these questions, we revisited breast tissues to explore their architecture and cellular composition through an exhaustive cytometric dissection of the tissue, which led to our development of a FACS procedure that reconciled every cell in the tissue (Figure 1). We isolated twelve cell populations from four lineages and developed techniques to culture nearly every one (Figure 2)^8^. Our system of defining breast cell types provides a comprehensive resource that can be used to explore interactions in more complex and physiologically relevant culture systems aimed at mimicking the tissue microenvironment.

Here, we explored the nature of each breast cell type by dissecting their transcriptomes. These were acquired by scaling up our FACS procedure and painstakingly sorting each cell type to purity. RNA was then isolated and sequenced, leading to the acquisition of transcriptomes that were of high depth and quality. Although the rare cell types yielded fewer cells and transcripts, concordance of transcriptomic profiles between all cell replicates illustrated the reproducibility of our methods (Figure 4). Analysis of transcripts identified uniquely expressed genes that have already helped us decipher spatially resolved transcripts from breast tumors. Bioinformatic analyses of transcriptional profiles further helped validate the identity of several cell types by providing information on expressed markers and revealing their functional commonalities to related cell types (Figure 6).

Over 20,000 genes were measured in each cell type, providing a wealth of information. As we explored the transcriptomes from every major cell type in the breast, there were so many exciting features that we had to come to terms with the fact that we couldn’t describe all that we encountered. Our solution was to build resource-sharing tools that aid exploration of our data^9, 10^. To demonstrate the utility of these resources, we addressed three pressing questions: a) What features distinguish each cell lineage? b) What gene types confer cell-specific functions? and c) What biological processes define each cell type?

To identify the cell lineages’ distinguishing features, we investigated the cellular pathways impacted by lineage-specific genes, which we defined through principal component analysis. The pathway enrichment analyses affirmed established attributes of each cell lineage while identifying others we had not previously associated with these lineages (Figures 10-14).

Focusing on the genes families that varied most among the different cell types provided us with an unconventional look at the types of genes contributing to the essential qualities of the breast cell types. We examined the expression of each gene, from each gene family, within every breast cell type, revealing the gene types that vary across cell types. We provide a full accounting of each gene family in a supplemental table and have meticulously described nine gene families whose functions extend from the nucleus (e.g., hormone receptors) to the exterior cell microenvironment (e.g., laminins, Figures 15-19).

To address our last question and develop a broad perspective of each breast cell type’s fundamental properties and individual functions, we performed a comprehensive assessment of the biological pathways and processes that are differentially enriched in each cell population. We performed fifty-five independent analyses identifying differentially expressed genes and enriched pathways between every possible cell-type comparison. These results are available as a downloadable interactive application (Pathway Analyzer^9^). We comprehensively analyzed six comparisons we believed were of the highest interest, as they revealed features of every major cell type and identified those distinguishing closely related cell types (Figures 21-33). As the field moves forward with single-cell and spatial transcriptomic approaches to explore and define the significance of cellular heterogeneity within tissues and tumors^98, 99^, we predict that the abundant number of transcripts measured here, within the full spectrum of FACS-purified breast cell types, will provide a strong foundation for identifying cell populations and interpreting results.

A broad wealth of information touching on many scientific disciplines is presented herein, and we recognize that developing deeper insights requires engagement and a focused exploration of the associated datasets. We offer novel tools to aid this exploration^9, 10^ and look forward to creating and sharing others. This body of work, presented here and in the adjoining article, presents a systematic approach for identifying, isolating, and culturing nearly every major breast cell type of epithelial, endothelial, and mesenchymal origin. We spent many years refining our FACS strategy, used multiple rounds of sorting to ensure purity, exhaustively stained tissues and primary cultures to validate and characterize cell types, and comprehensively and painstakingly analyzed each transcriptome. This rich resource, which we submit here, has provided countless insights and has already changed our perception of the cells composing this elaborate and beautiful tissue.

## Methods

### Tissue acquisition and processing

Breast tissues from reduction mammoplasties were obtained from the Cooperative Human Tissue Network (CHTN), a program funded by the National Cancer Institute. All specimens were collected with patient consent and were reported negative for proliferative breast disease by board-certified pathologists. The University of California at Berkeley Institutional Review Board and University of New Mexico Human Research Protections Office granted use of anonymous samples through exemption status, according to the Code of Federal Regulations 45 CFR 46.101. Upon receipt, several fragments (roughly 2cm^2^) were embedded in optimal cutting temperature compound (OCT) in tissue cassettes, flash-frozen in nitrogen, and archived at -80°C for later cryosectioning and staining. The remaining tissue, typically 20-100g, was thoroughly rinsed with phosphate-buffered saline and processed to organoids as previously described (gentle agitation method)^100^. Briefly, this included manually mincing and scoring tissues with a scalpel and incubating them overnight (12-18 hrs, 37°C) with 0.1% collagenase I (Gibco/Invitrogen) in Dulbecco’s Modified Eagle Medium containing 100 U/ml penicillin, 100 mg/ml streptomycin, and 100 mg/ml Normocin™ (Invivogen, San Diego, CA). The resulting divested tissue fragments (organoids) were collected by centrifugation (100g x 2 min) and either archived in liquid nitrogen (90%FBS+10% DMSO) or immediately processed to cell suspensions for FACS analysis. Tissues used for RNA-sequencing were immediately processed upon receipt and kept at 4°C during processing and FACS sorting, except for the overnight collagenase digestion at 37°C. RNA-sequenced cells were from fresh specimens and were not previously archived in nitrogen.

### Antibodies

A list of antibodies and reagents used in this study is provided in Figure 3—Table Supplement 8. The table includes antibody clone designations, conjugations, isotype, supplier product numbers, dilution factors used.

### Cryosectioning and Immunostaining

Immunofluorescence was performed on 10-80 mm cryosectioned tissue sections as described in the adjoining article^8^. Briefly, specimens were fixed with 4% paraformaldehyde for 5 minutes at 23°C, which was followed by 4% formaldehyde / 0.1% saponin for 5 minutes at 23°C. Tissues were rinsed for 20 minutes in wash buffer (0.1% saponin/10% goat serum in PBS) and then incubated with primary antibodies, typically overnight at 4°C. Following the primary incubation period, samples were washed and incubated with secondary antibodies conjugated with Alexafluor 488, 568, 594, or 647 (Thermo). After 1 hour incubation at 23°C, samples were rinsed in PBS and their nuclei counterstained with 300nM DAPI (4’,6-Diamidino-2-Phenylindole, Dihydrochloride, Thermo). Coverslips were mounted with Fluoromount G (Southern Biotech). Images were captured using a Zeiss LSM710 confocal microscope, Zeiss Axioscope, or EVOS FL Auto imaging station. If needed, image contrast was applied to the entire image using Photoshop and were annotated with the antibodies used to stain the tissue.

### Cell Preparation for Flow Cytometry and FACS analysis

To prepare cell suspensions for FACS and flow cytometry analysis, we thoroughly rinsed the organoids in PBS, then pelleted them via centrifugation at 100g x 2 min. After removing PBS by aspiration, the organoids were suspended in 1 ml of ‘Cell Dissociation Reagent’ (Sigma# C5914; or Thermo# 13150016), incubated for two minutes at room temperature, upon which 3ml trypsin (used for RNA-sequenced samples) was added (0.25%, Thermo 25200072; Note that we have more recently found TrypLE better preserves CD49f staining, although it dampens CD34 resolution). Samples were incubated by hand/body temperature, thus permitting visual inspection and brief (1-3sec) pulse vortexing every 30-60 seconds. After the mixture became cloudy from the dissociating cells (about 8-10 min), the organoids/cells were gently and repeatedly pipetted through a 16- gauge needle until clumps dissipated. Afterward, we filtered cell suspensions through a 100mm cell strainer and added 3ml 0.1% w/v soybean trypsin inhibitor to stop the digest (Sigma# T9128). The suspensions were then filtered through a 40mm cell strainer, rinsed in 10-20ml PBS, and pelleted by centrifugation at high centrifugal force (400g x 5 min.—or longer, until all cells had pelleted, which we confirmed by microscopic examination of the supernatant). The cells were then rinsed in 10 ml Hanks balanced salt solution/1% BSA(w/v), counted, and again pelleted by centrifugation. After centrifugation, nearly all the Hanks/BSA was aspirated, leaving the cell pellet with roughly 60ml of Hanks/BSA. The pellet was resuspended in this small volume, and then we added FACS antibodies to the samples and incubated the samples on ice, covered, for 30 minutes (See Figure 3- table supplement 8 for antibodies). Following the incubation, the cells were rinsed in Hanks/1% BSA, centrifuged (400g x 5 min), and resuspended in Hanks/BSA with To-Pro-3 viability marker (diluted 1:4000, Thermo T3605), then filtered again through a 40mm cell strainer cap (into the FACS tube). We did not use DNase enzyme or hypotonic RBC lysis solutions on the cell preparations because we found they were unnecessary. We also wanted to avoid any potential deleterious effect these treatments may have on the cells, such as reducing cell viability or altering gene expression levels. We, again, also conscientiously chilled samples during cell preparation and the sorting procedure for similar concerns.

### Flow Cytometry

The FACS panel and gating strategy outlined herein was developed using a BD FACS Vantage 8-channel cytometer (FACSDIVA software) and has since been adapted and reproduced using a 14-channel SONY SY3200. PMT voltages were optimized on both instruments by analyzing individually stained compensation beads (AbC™ Anti-Mouse Bead Kit, Thermo # A10344) at increasing voltage intervals for each detector. Because the FACS Vantage has only eight available channels, it was necessary to use two markers (podoplanin and CD45) in a single channel (A488), which we experimentally validated. To-Pro-3 was used as the viability marker, so the corresponding 405nm channel (455/50) could be used for BV421. This channel was dedicated to CD49f to provide a maximum resolution of cell populations, which is essential to this sorting strategy. The considerable compensation typically required between the heavily overlapping dyes PE/Cy5 and To-Pro-3 was not needed, as the To-Pro-3 negative (viable) cells were used for downstream analyses, allowing us to use the PE/Cy5 channel for Thy1. Negative controls consisted of unlabeled beads and cells incubated with isotype control antibodies conjugated to each fluorophore. Sort times typically ranged 14-18 hours (50-120 million cells stained— needed to yield enough cells for RNA-sequencing). Cells were chilled at the sample intake and collection chambers, and all samples were chilled on ice during the entire experiment. All media and PBS were filtered through a 0.2mm filters to prevent the FACS machine from triggering events from crystals and debris. Staining profiles of cells collected at the beginning and end of the sorts (sometimes separated by as many as 18 hours) were identical and did not show any signal loss or appreciable decrease in viability. Because our machines could sort cells into a maximum of 4 tubes, we first sorted cells into four groups, containing 2-4 populations each. These mixtures were: Tube1 (CD34Pos) and contained Pops 7,8,9; Tube 2 (Thy1^Pos^) contained Pops 3,6; Tube 3 (Luminal) contained Pops 1&2; and Tube 4 (Null gate) contained Pops 4,5,10,11. During the first round of sorting, cells were typically sorted at approximately 8,000 cell/sec. We followed this initial enrichment with purity sorts for each cell type, using stringent sort masks, sorting each population into individual tubes at a rate of approximately 2,000 cells/sec. FACS data were analyzed using Flowjo software (version v7.6.3 – v10.8), Tree Star Inc./ Becton, Dickinson & Company).

### RNA Isolation and Sequencing

Following the overnight digestion of tissues with collagenase, we centrifuged the samples at 100g x 2 min. This produced aqueous and lipid phases that separated under centrifugal force. The top lipid phase was collected, visualized microscopically for cellular content, and then processed immediately for RNA extraction using Qiagen RNA Lipid Tissue Mini Kit (Qiagen 74804), according to the manufacturer’s instructions. Total RNA from the FACS-purified cells was isolated immediately after FACS sorting using Qiagen RNA Mini (>100,000 cells, Qiagen 74104) or RNA Micro kits (<100,000 cells, Qiagen 74004) according to the manufacturer’s instructions. Total RNA was treated with DNAse I for 30 minutes at 37°C (Ambion DNA-*free* DNA Removal kit, Thermo AM1906), then treated with DNAse Inactivation beads. RNA was then concentrated with RNA Clean & Concentrator™-5 columns (Zymo R1015), quantified by Nanodrop and Agilent Bioanalyzer assays, and stored at - 80°C.

### RNA Library construction and sequencing

mRNA was captured using poly-T coated dishes (Clontech) and used for library construction using SMARTer technology (SMARTer Stranded RNA-seq Kit, Takara Biotech) following the manufacturer’s guidelines. For low abundance samples, mRNA was captured on biotinylated LNA coated beads, magnetically captured and rinsed 3x in binding and wash buffer (10 mM Tris-HCl (pH 7.5), 1 mM EDTA, 2 M NaCl). mRNA was eluted from Dynabeads in 10ml volume of elution buffer, following by a 5 min. incubation at 68°C. It was then used for first-strand cDNA synthesis (SMARTer Stranded RNA- seq Kit). Sample clean-up and removal of adapter-dimers) was performed using Ampure XP beads (Beckman Coulter), according to the manufacturer’s protocol. Specific attention was given to the samples to prevent under- or over-drying the beads, which we found to reduce PCR efficiency. PCR amplification of the RNA-Seq library was performed using SeqAmp DNA Polymerase (Takara Bio, cat# 638509). Each library was then purified prior to sequencing using SPRI AMPure Beads (Beckman Coulter). Sequencing was performed using Illumina Trueseq v.2 technology, targeting approximately 30 million paired-end reads for each cell population (Illumina HiSeq 2x100bp).

### RNA-Seq pre-processing and quality control assessment

We processed raw count data from aligned fragments (Figure 3—table supplement 2) using R programming language (RStudio) and associated BioConductor packages. Quality control analyses (Cook’s distance box plot, covariance matrix, sample correlation matrix) were performed using the DESeq2 package^13^. For data visualization (donut and heatmaps), the count matrix data was transformed by two methods used in the DESeq2 package: (i) regularized log transformation (rlog), and (ii) variance- stabilizing transformation (vst), both normalized to library size. Quality control assessment of expressed transcripts across sorted cell samples was conducted, and one sample was removed from statistical analyses as it was an extreme outlier (Epithelial Population 4 Sample N239). This was determined by: (i) distance matrix, (ii) Principal Component Analysis, and (iii) Kolmogorov-Smirnov test that compares each sample to the reference probability distribution of the pooled data. We included this sample in downstream visualizations for reference purposes, for example, the bar and donut graphs used throughout the manuscript, which display rlog transformed values on both log2 (bar graphs) and linear (donut graph) scales.

### Differential Gene Expression Analysis

All differential expression analyses were performed using DESeq2^13^. The DESEQ2 likelihood ratio test (LRT) was used to identify genes differentially expressed genes across all cell types (Figure 3—table supplement 5); These genes were then used to calculate the proportion of differentially expressed genes within each HGNC families/group in our expression dataset (https://www.genenames.org/data/genegroup).

Differential expression p-values were corrected for multiple hypothesis testing using Benjamini- Hochberg method across all genes and all 55 possible pairwise contrasts. Volcano plots of the DESeq2 contrasted cell types were generated using the EnhancedVolcano R/Bioconductor package^101^

### Principal component analysis

Principal component analysis was performed on VST transformed data (subsetting without tissue and bulk cell controls) using the pcaExplorer R/Bioconductor package^102^. Loading factors from the ends of principal components 1, 2, 3, and 4 were further analyzed using the ClueGo^21^ Cytoscape package, referencing the Gene Ontology biological processes (GOBP) and Reactome pathway databases^18, 20^. Subsequent enriched pathways were grouped automatically by likeness to obtain a condensed list of defining features for each cell class.

### Gene Set Enrichment and Pathway Analysis

Pathway enrichment analysis was performed using Broad Institute’s GSEA software (version 4.3.2)^52^. We pre-ranked our individual gene lists using the DESeq2 calculated Fold changes and Benjamini-Hochberg adjusted *p*-values from all 55 possible pairwise contrasts (Figure 3—table supplement 7). The rank metric was calculated by: (sign[Fold Change]* -log10{*p*-value}). GSEA analysis was performed using a classic, unweighted enrichment statistic (p=0)^52^ and was used to evaluate the enrichment of Reactome pathways^18^ Pathways were obtained via the Molecular Signatures Database^54^, and we included pathways with sizes between 15-200 genes. The resulting analysis produced a collection of enriched pathways and calculated statistics (1000 permutations to correct for multiple hypotheses testing) for each of the 55 pairwise comparisons. These data became the foundation for the Excel-based Pathway Analyzer tool.

### Enrichment Map/Cytoscape

Visualization of enriched gene sets was performed using the Enrichment Map Cytoscape plugin^52, 58^, with applied edge cut-offs of 0.5 and FDR cutoffs of 0.1 (10%) for most comparisons, with the exception of the 8v12 comparison between fibroblasts and adipocytes, which had a FDR of 0.2 (20%). The produced nodes (biological pathways) were manually adjusted/grouped into themes for presentation purposes using the Reactome hierarchy as a reference. Annotations of select biological themes and notable genes were added manually.

## Supplemental Resources

### Pathway Analyzer

This supplemental tool integrates regularized log transformed transcript levels (mRNA-sequencing values) and enriched biological pathways identified by Gene Set Enrichment Analysis (GSEA). Data from every possible pairwise comparison of cell types is embedded in the tool, which is supplied as a MS Excel file. The file is approximately 300Mb and contains computationally intensive algorithms (e.g., we run the file on both Windows and MAC-based computers (Surface Laptop Studio [Core i7/3.3GHz with 32Gb RAM], Surface Book 2 [Corei7/4.2GHz with 16Gb RAM], Dell Inspiron Desktop [Corei5/3.2GHz with 8Gb RAM], MacBook Pro [Corei7/2.1GHz with 16Gb RAM]), and we recommend using it on computers with ≥16Gb RAM).

### Breast Cell Atlas

**Transcript Finder** http://breastcancerlab.com/breast-atlas/. This online tool facilitates exploration of mRNAs expressed by the breast cells. When provided individual gene names (official gene symbol), the tool will display the transcript levels measured in each cell type. Regularized log transformed values are provided on a log2 and linear scale (bar and donut graph, respectively), similar to those presented in the many figure elements contained herein. This application was coded using the R/Shiny package^103^.

## Author Contributions

Conceptualization, WCH; FACS & Methodology, WCH; RNA isolation, WCH; mRNA Library production, ZW; RNA-seq alignment, quality control, and count matrix formation, AC; Bioinformatics, KDT, RS, AC, and WCH; Pathway Analyzer: KDT and WCH; Breast Cell Atlas: Transcript Finder: KDT, WCH; Immunostaining, WCH, KT, KDT, YLM; Writing – Original draft, WCH and KDT; Writing – Review & Editing, WCH, KDT, RS; Funding, WCH.

## Acknowledgments

We thank Mina Bissell, Irene Kuhn, Alex Bazarov, Ritu Mukhopadhyay (Lawrence Berkeley Laboratory), Sandy Borowsky (U.C. Davis), Kornelia Polyak (Dana Farber Cancer Institute), Eric Prossnitz (University of New Mexico Health Sciences Center), Curtis Hines (Sandia National Laboratory, retired) for thoughtful scientific discussions and/or for critical review of the manuscript. We thank Tristan Oper, Alice Rindestig, Kremena Karagyozova, Alyssa Cozzo, Gaelen Stanford-Moore, Maria Rojec, Anya Afasisheva, and Sun-Young Lee for their technical assistance. We also thank Dana Pe’er, Ambrose Carr, Zhenmao Wan, and the staff at Columbia University’s Genomic Sequencing Core for Illumina RNA-sequencing. Much of the analyses presented herein would not have been possible without Reactome’s curation of genes, pathways, and publicly available resources. We thank Reactome’s Principal Investigators and the team of dedicated scientists that have contributed to the Reactome Project. We also thank Kerry Wiles and Erik Brooks of the Cooperative Human Tissue Network, and the anonymous women who graciously donated their remnant surgical tissue to science. This project depended on flow cytometry instruments provided at the University of New Mexico Flow Cytometry and High Throughput Screening Resource and we thank Cecelia Rietz and Wade Johnson for their assistance. Finally, a very special thank you to Michelle Scott, Lawrence Berkeley National Laboratory Advanced Microscopy and Flow Cytometry facility, for her expert technical guidance and support.

---

**Figure 3—Table supplement 8.** Antibodies

| **Tissue Staining (Immunofluorescence)** |  |  |  |  |
| --- | --- | --- | --- | --- |
| **Antibody specificity** | **Clone** | **Conjugated** | **Source** | **Dilution** |
| Alpha-Smooth Muscle Actin | 1A4 | No | Mouse IgG2a, $\kappa$ , Lab Vision/Neomarkers MS-113-P0 | 1:100 |
| CD29 | TS2/16 | A488 | Mouse IgG1, $\kappa$; Biolegend 303015 | $\frac{1}{2}$ assay<sup>¶</sup> |
| CD49f | GoH3 | No | Rat IgG2a, $\kappa$; Biolegend 313602 | $\frac{1}{2}$ assay<sup>¶</sup> |
| E-Cadherin (CD324) | 24E10 | No | Rabbit mAb; Cell Signaling 3195S | 1:100 |
| E-Cadherin (CD324) | 67A4 | A488 | Mouse IgG1, $\kappa$; BioLegend 324110 | $\frac{1}{2}$ assay<sup>¶</sup> |
| erbB2/Her2 (CD340) | 24D2 | PE | Mouse IgG1, $\kappa$; Biolegend 324406 | $\frac{1}{2}$ assay<sup>¶</sup> |
| Keratin 14 | Polyclonal | No | Rabbit Polyclonal; Thermo/Lab Vision RB-9020-P1 | 1:400 |
| Keratin 14 | Polyclonal | No | Chicken Polyclonal; Biolegend 906001 | 1:400 |
| Keratin 18 | DC10 | No | Mouse IgG1; Thermo/Lab Vision MS142P0 | 1:400 |
| Keratin 19 | A53-B/A2.26; same as Ks19.1 | No | IgG2a/$\lambda$; Lab Vision/Neomarkers MS-198-PABX | 1:400 |
| Keratin 8 | | No | Mouse IgG2a $\kappa$ | 1:400 |
| P63 | 4A4 | None | Abcam ab32353 | 1:100 |
| Pan-Laminin | Polyclonal | No | Rabbit Polyclonal; L9393 | 1:400 |
| Pan-Laminin | Polyclonal | No | Rabbit Polyclonal; L9393 | 1:400 |
| Podoplanin | NC-08 | No | Rat IgG2a, $\lambda$ | $\frac{1}{2}$ assay<sup>¶</sup> |
| **Facs Panel Antibodies / Viability Dyes** |  |  |  |  |
| CD49f (alpha-6 integrin) | GoH3 | BV421* | BioLegend 313624, 313612, 313607 | 2.5μl<sup>§</sup> |
| CD24 | ML5 | BV605* | BioLegend 311124, 311106, 311108 | 2.5μl<sup>§</sup> |
| CD227 (Muc1) | 16A | PE | BioLegend 355604, | 4μl<sup>§</sup> |
| Podoplanin | NC-08 | A488<sup>†</sup> | BioLegend 337006, 337004 | 2.5μl<sup>§</sup> |
| CD45 | HI30 | A488<sup>†</sup> | BioLegend 304052, 304032 | 3μl<sup>§</sup> |
| CD90 (Thy1) | 5E10 | PE-Cy5 | BioLegend 328112, 328134 | 3µl<sup>§</sup> |
| CD34 | 581 | PE-Cy7 | BioLegend 343516 | 1-2µl<sup>§</sup> |
| CD10 | HI10a | APC-Cy7 | BioLegend 312212, 312204 | 3µl<sup>§</sup> |
| TO-PRO™-3 Iodide | N/A | (642/661) | Thermo ( T3605) | 1:4000 |

\* *Brilliant Violet*; † *Alexa Fluor*; (Biolegend); § 1x volume per 6x10<sup>6</sup> cells stained in FACS experiments (up to 36x10<sup>6</sup>, after which cells received 6x volumes; max stained cells =120x10<sup>6</sup>); titration required and must be matched to cytometer. ¶ Percentage of manufacturer's suggested assay volume per 100ul volume (antibody is prediluted).

| List of Table supplements and Data |  |
| --- | --- |
| Figure 3—table supplement 1 | Sample Treatment.xlsx |
| Figure 3—table supplement 2 | CountMatrix_rLog_VST.xlsx |
| Figure 3—table supplement 3 | PCA_Loading_Factors.xlsx |
| Figure 3—table supplement 4 | GO from PCAs.xlsx |
| Figure 3—table supplement 5 | DESeq2_LRT.xlsx |
| Figure 3—table supplement 6 | Gene Families_DE Genes.xlsx |
| Figure 3—table supplement 7 | DESeq2_files.xlsx |
| OSF; https://osf.io/gfhc3/ | Cytoscape/Enrichment Map files (1v2, 2v3, 2v9, 6v8) |
| OSF; https://osf.io/hpb9t/ | Pathway Analyzer<sup>9</sup> |
| http://breastcancerlab.com/breast-atlas | Breast Cell Atlas: Transcript Finder (R/Shiny App)<sup>10</sup> |

**Figure 1—figure supplement 1.**
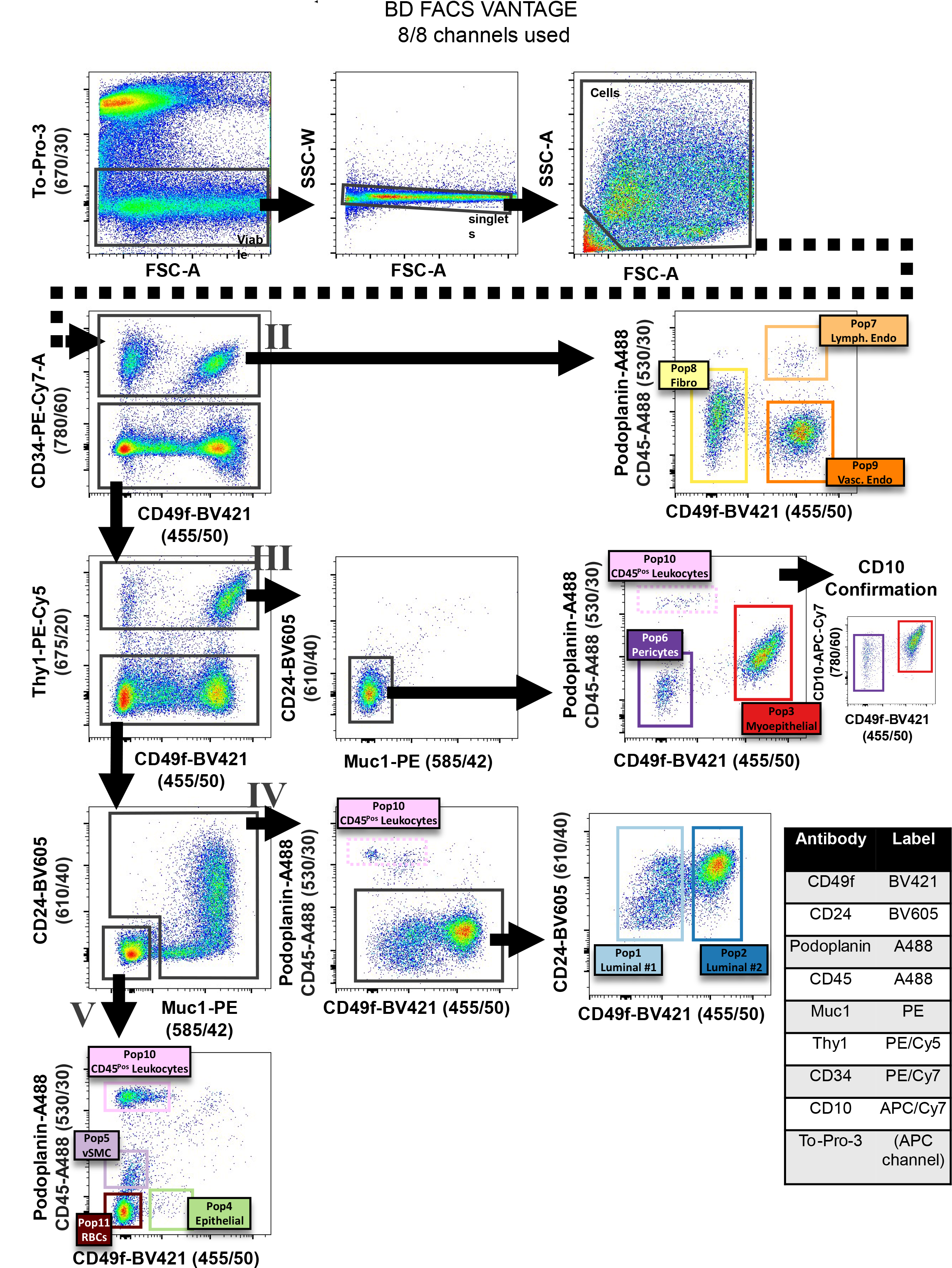
FACS strategy designed for an 8 channel FACS machine (BD FACS Vantage). (a) FACS gating scheme used to isolate the RNA-sequenced cell types (antibody panel designed for an 8 channel FACS machine- BD FACS Vantage). Representative FACS data are of cells derived from disease-free reduction mammoplasty tissue surgically resected from a 22-year-old female (sample #N239).

**Figure 2—figure supplement 1.**
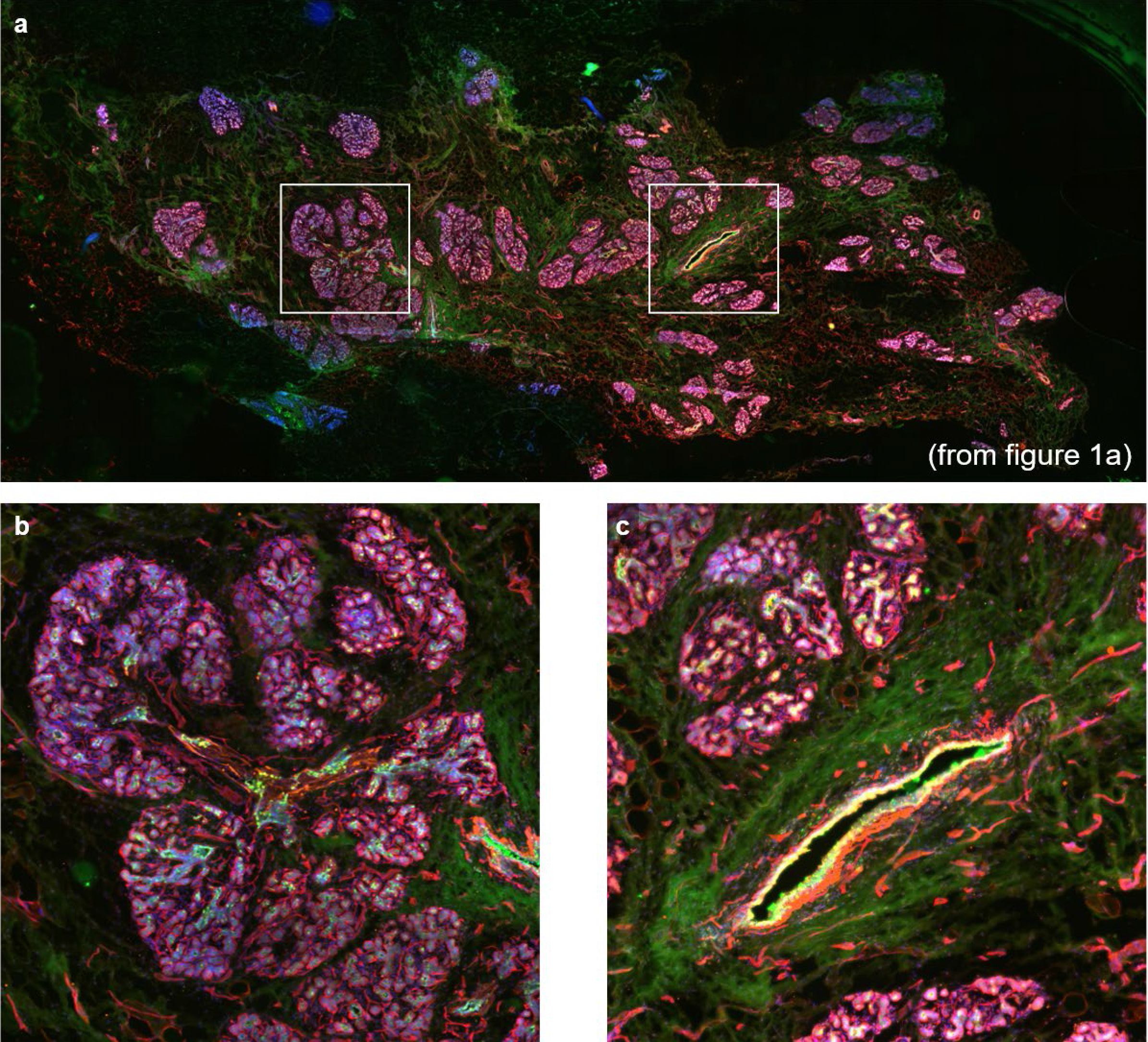
(a) High-resolution image of the immunostained breast tissues provided in Fig. 2a. **(b)** magnified area indicated by white box in a (on left) containing several breast lobules (a.k.a, terminal ductal lobular units). **(c)** magnified area indicated by white box in a (on right) containing the cross section of a large lactiferous duct. The duct is surrounded by numerous blood vessels running parallel to the duct and is embedded in a collagenous-rich matrix.

**Figure 2—figure supplement 2.**
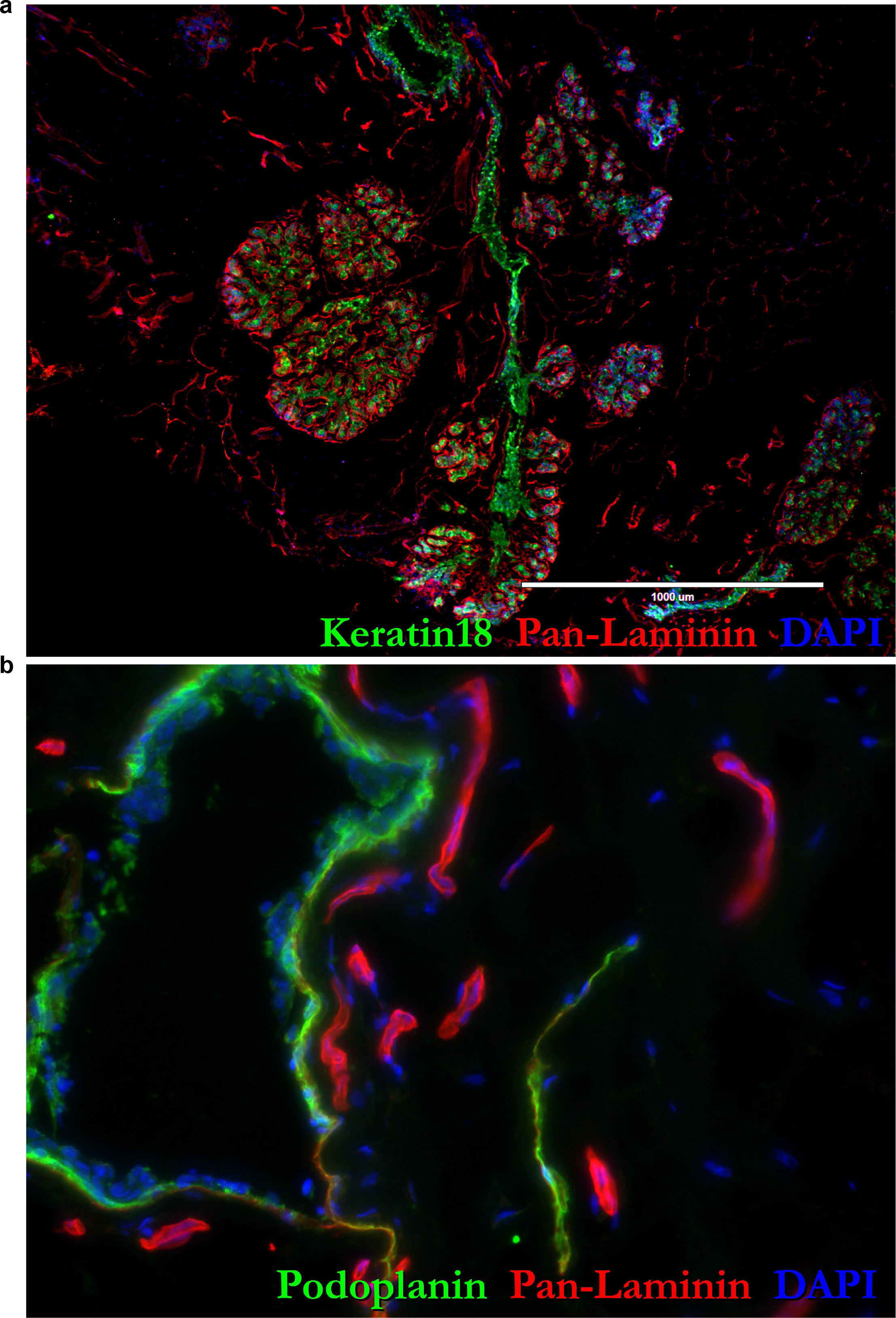
(a,b) High-resolution (label-free) images of the immunostained breast tissues provided in Figs 2c,d.

Figure 2- Figure supplement 3. (a,b) High-resolution (label-free) images of the immunostained breast tissues provided in Figs 2e.f.

**Figure 3—figure supplement 1.**
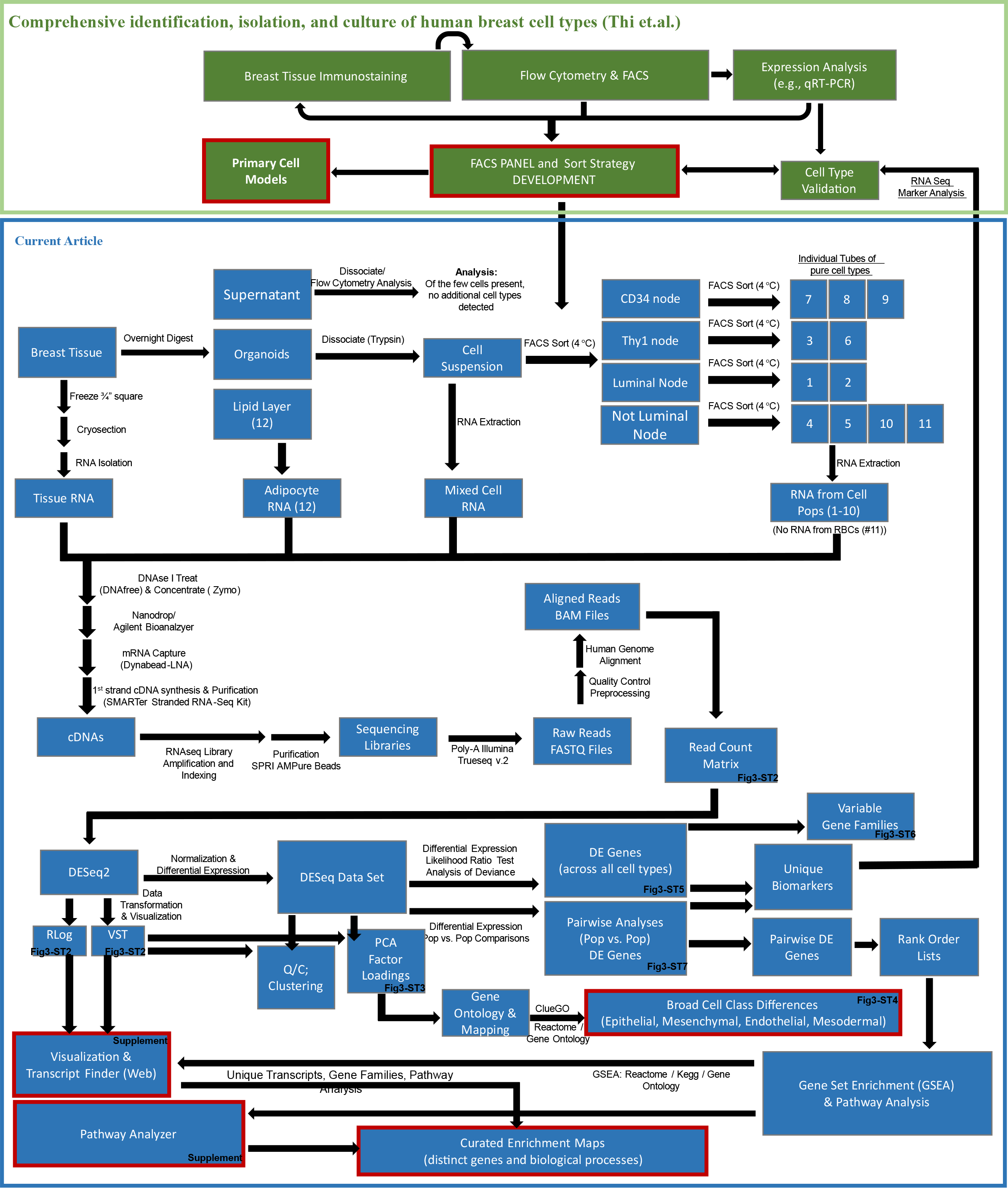
Project workflow. Diagram of the sequential processes and analyses presented in the adjoining and current articles. The adjoining article (in green) describes the development and validation of a FACS isolation strategy for separating all twelve major breast cell types, culminating with the development of primary cell models for nine major breast cell types. The current article (in blue) explores the nature of cell type through bulk-mRNA-sequencing and analysis of transcript levels measured in each FACS-purified population. Analysis included principal component analysis, differential gene expression, and gene set enrichment analysis. Red-outlined boxes indicate the major products from each publication.

Figure 3—table supplement 1. Sample information/Meta Data. Table of sample information, containing age/sex/race information supplied with the acquired tissues (B=black; W=white), Patient number, FACS gate, cell type, cell yields, RNA-isolation strategy/kit, elution volumes, nanodrop concentrations, DNase treatment used, Bioanalyzer results (concentration and integrity number), RNA-sequencing batch, and calculated reads in each sample.

Figure 3—table supplement 2. RNA-sequencing count matrix.

Figure 3—table supplement 3. PCA loading factors. Sheet 1, ‘All Pops,’ contains loading factors calculated from the analysis of all cell types (Pops 1-12). Sheet 2, ‘Endothelial Removed,’ contains loading factors calculated from the analysis of non-endothelial cell types (Pops 1-7,8,10-12).

Figure 3—table supplement 4. Tabulated Pathway Enrichment Results (ClueGo/Reactome/Gene Ontology Biological Processes) that used the 200 loading factors associated with the four cell lineages: a) Epithelial end of PC1, b) Leukocyte end of PC2, c) Mesenchymal end of PC1No Endo, d) Endothelial end of PC3, and the perivascular and myoepithelial lineage subtypes e) Perivascular cell end of PC3, and f) myoepithelial end of PC4.

Figure 3—table supplement 5. RNA-Seq DESeq2 Likelihood Ratio Test (LRT) statistics, p- values, and Benjamini Hochberg adjusted p-values for differentially expressed genes with LRT adj.p-val<0.05. These results were used to calculate the proportion of differentially expressed genes in each HGNC gene family.

Figure 3—table supplement 6. Table of the 1,421 HGNC Gene families, number of genes in each family, and the calculated number of differentially expressed genes in each family (using the DESeq2 Benjamini Hochberg Adj. p-value threshold of ≤ 0.1).

Figure 3—table supplement 7. Excel workbook containing 55 DESeq results (on separate tabulated worksheets) from each possible pairwise contrast of the 11 seqenced breast cell types; e.g. Pop1v2 tab contains results from the contrast of Pop1 (ERPos luminal epithelial cells) and Pop2 (ERNeg luminal epithelial cells). Tab 1v2 also contains the calculated rank metric used for GSEA analysis; i.e., sign([Fold Change] * -log10 {p-value}).

Figure 3—table supplement 8. Antibodies. Antibody clones used for immunostaining and FACS.

**Figure 4—figure supplement 1.**
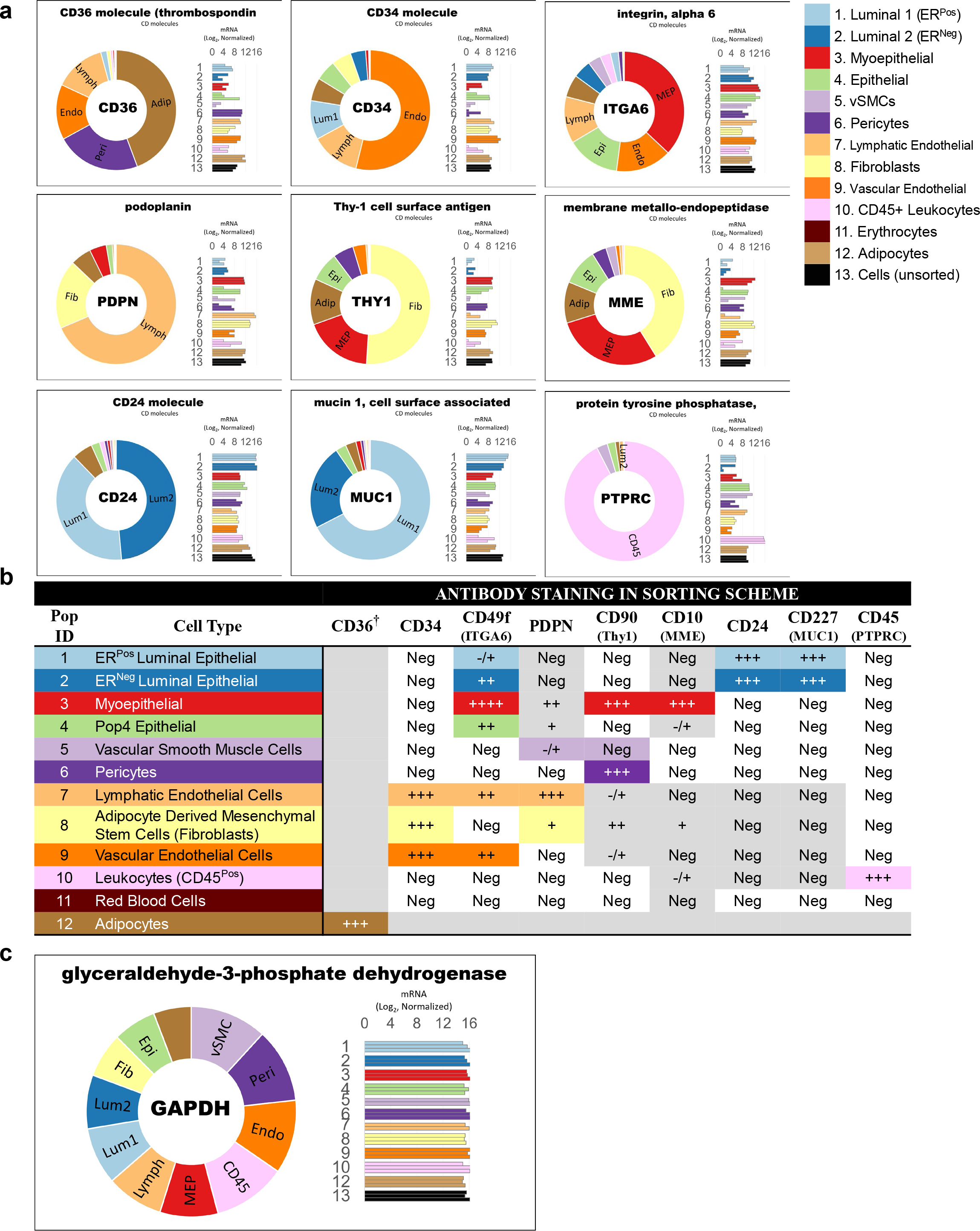
mRNA levels correspond to FACS staining. (a) Normalized mRNA values for genes encoding FACS markers used to purify cell types from tissues (and CD36, which was later used to purify cultured adipocytes). Transcript levels are provided on a) a log2 scale (bar graphs containing each biological replicate) and b) a linear scale (donut graphs of the replicate’s median value), which are both color-coded by cell type. **(b)** Schematic indicating relative FACS staining levels of each marker in the twelve breast cell types. “Neg” indicates relative negative staining (i.e., compared to other populations, not absolute negativity), “-/+” indicates near background level staining, and the increasing number of “+” indicates greater staining intensity. **^†^** CD36 is a marker used to purify cultured adipocytes but was not used to purify adipocytes prior to RNA-sequencing in this analysis. **(c)** Normalized transcript levels for the commonly used internal control gene GAPDH (rlog, DEseq2) are presented on a log2 scale (bar graph) and linear scale (donut graph of median value).

**Figure 5—figure supplement 1.**
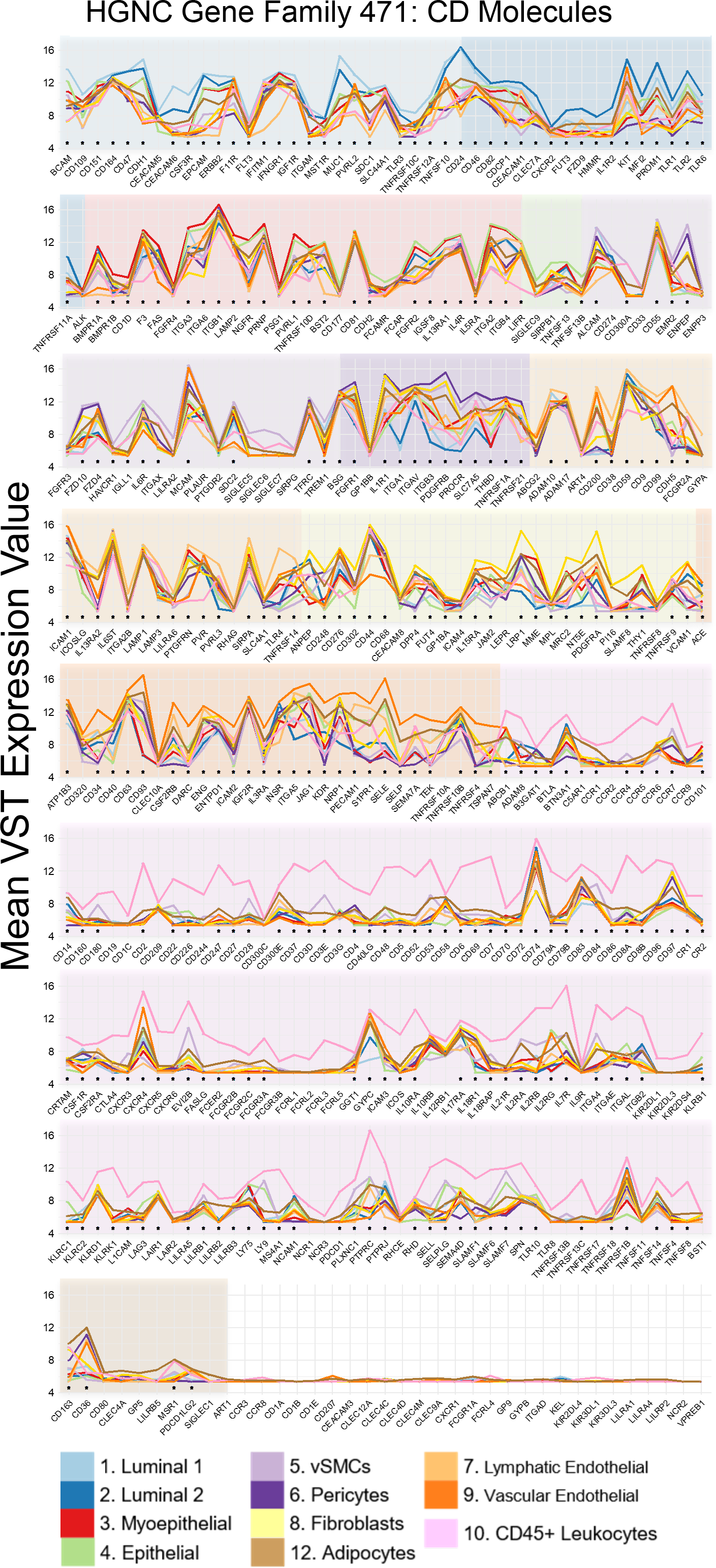
CD antigen genes. Median mRNA expression levels of genes encoding CD antigens (VST, log2 scale). The genes are arranged by the cell type that expressed them at their highest level, as indicated by the background shading specific to each cell type. Genes differentially expressed across cell populations are annotated with (*). CD antigen genes in the uncolored panel were not expressed by any breast cell type.

**Figure 7—figure supplement 1.**
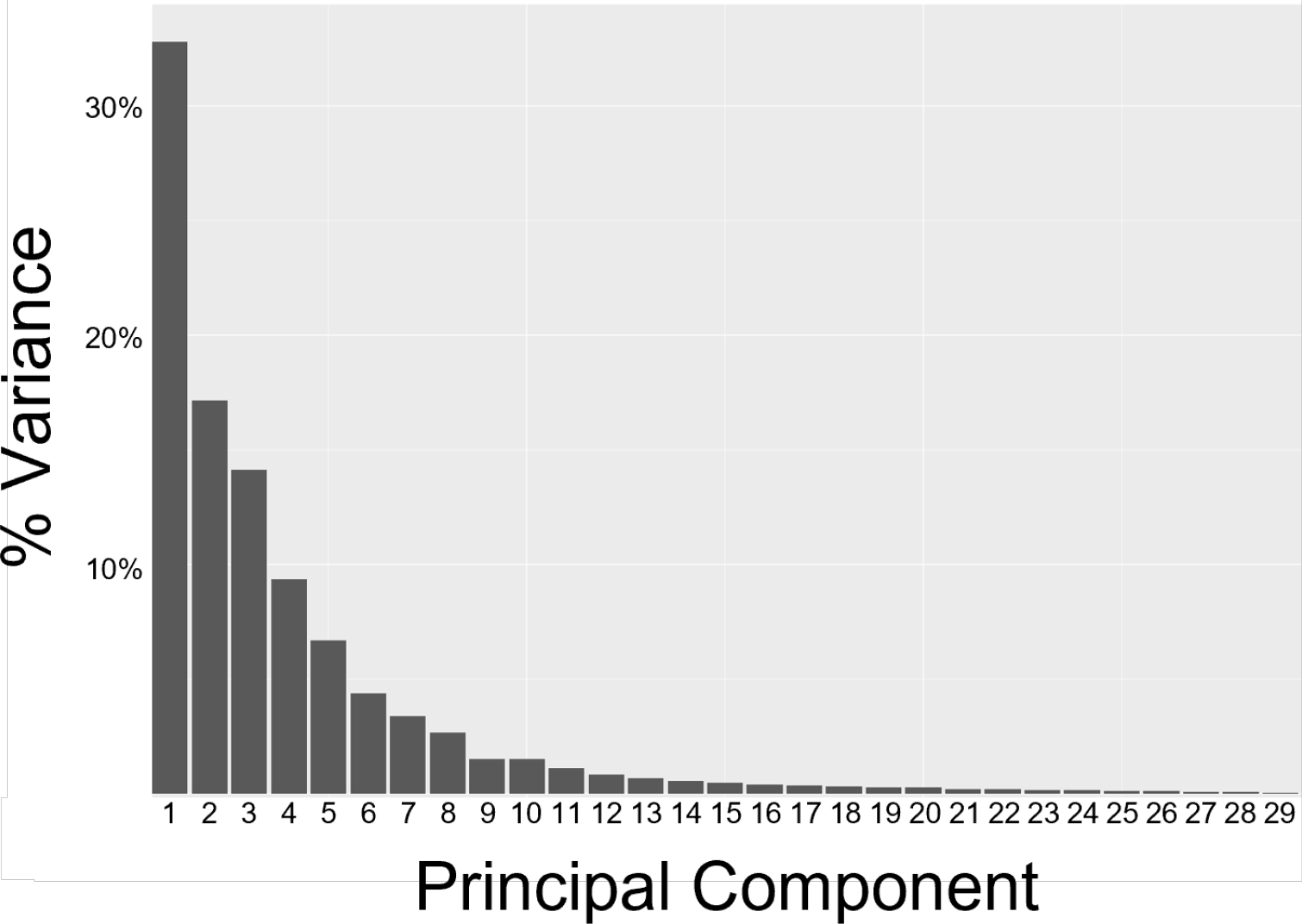
Variance captured by principal components. A screeplot shows the sample variances captured by each principal component. 71% of the total variance is captured by the first four components.

**Figure 7—figure supplement 2.**
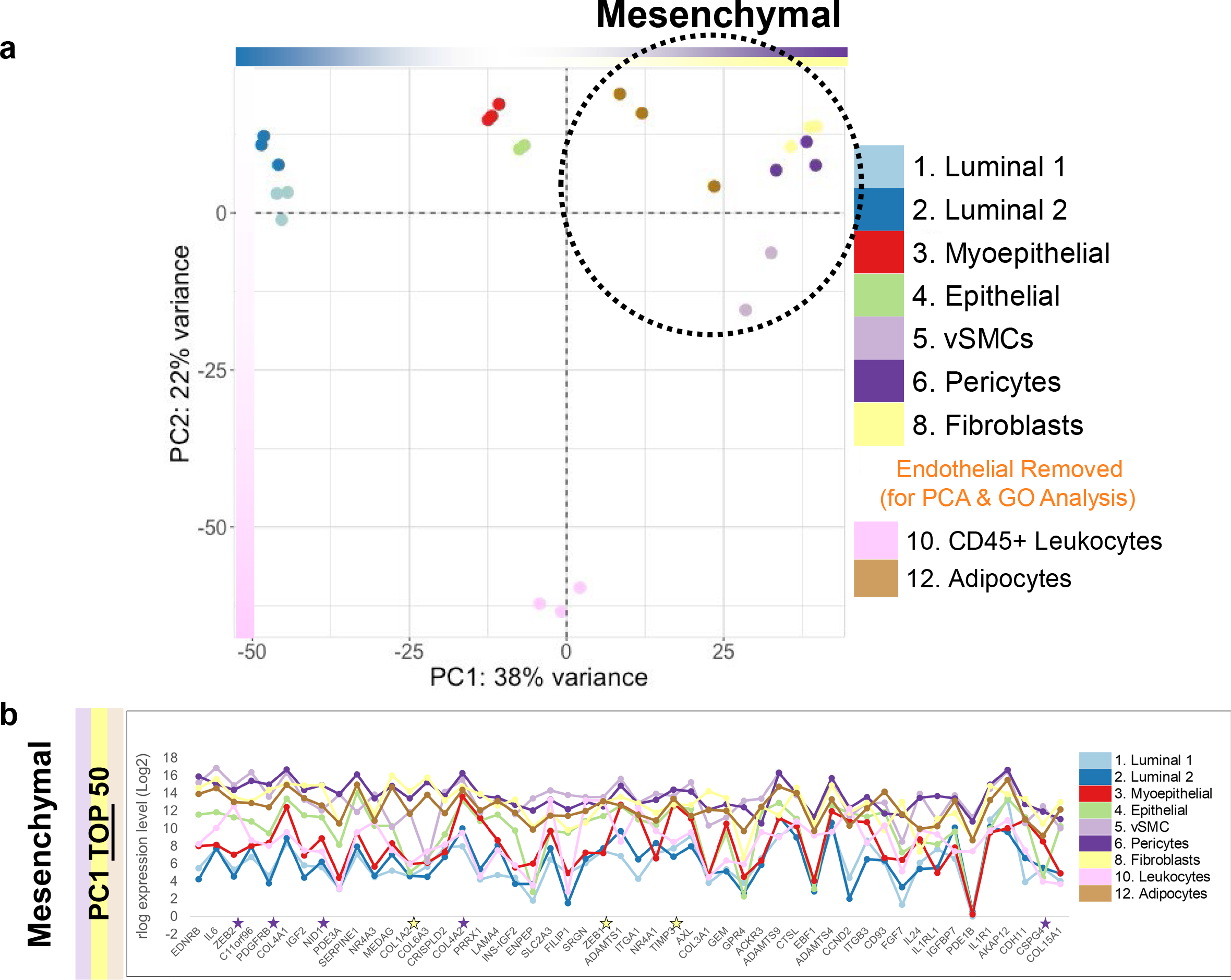
PCA with endothelial cells removed. (a) PCA projection of breast cell types, after removal of the lymphatic and vascular endothelial cells, resolves the mesenchymal cell types on PC1^NoEndo^. **(b)** Median mRNA expression of the 50 loading factors (genes) contributing to the positive end of PC2 (top end of the list; rlog, log2 scale). Stars (↔) denote common markers of pericytes (purple) and fibroblasts (yellow).

**Figure 10—figure supplement 1.**
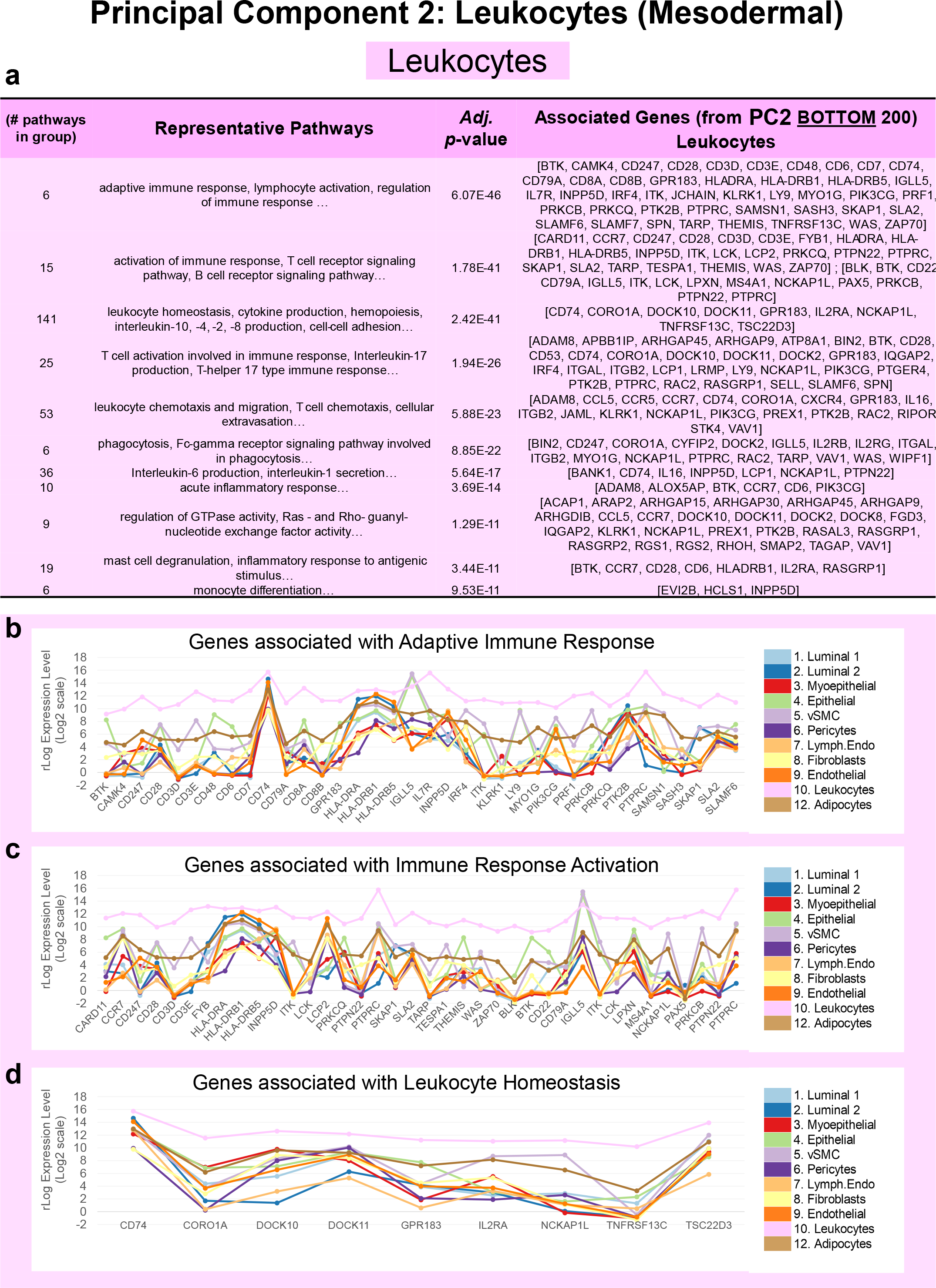
Pathway analysis of leukocyte genes (PC2). (a) Grouped GO/Reactome terms associated with the 200 genes contributing to the positive end (bottom of list) of PC1 ordered by adjusted p-value. Median mRNA expression (rlog transformed) for PC2 genes associated with select pathways, including **(b)** a*daptive immune response*, **(c)** *immune response activation*, and **(d)** *leukocyte homeostasis*.

**Figure 12—figure supplement 1.**
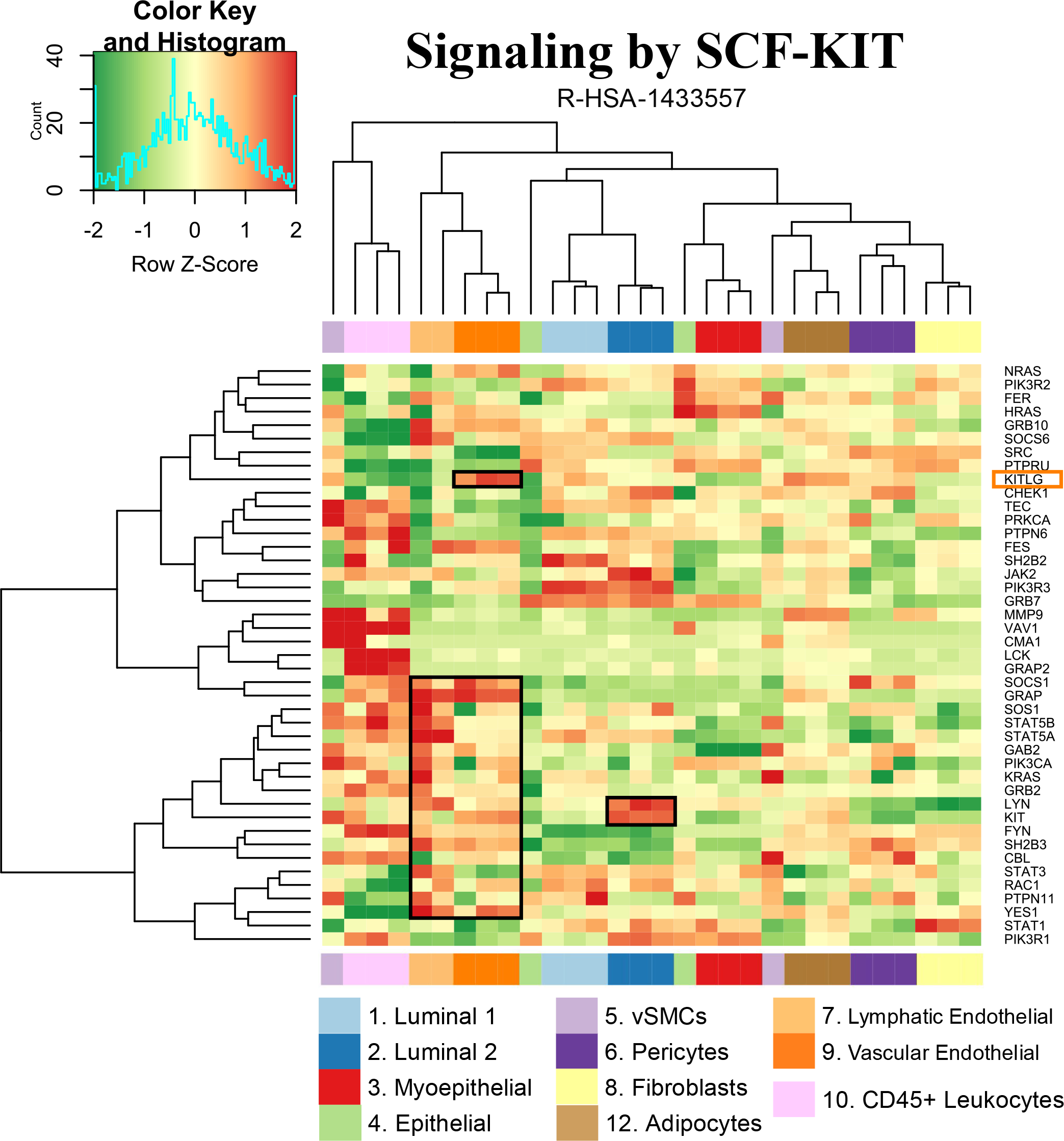
Transcript levels of genes composing the *signaling by SCF-KIT* pathway. Transcript levels for each gene in the Reactome pathway s*ignaling by SCF-KIT.* VST gene levels are relative (normalized within each row of the heatmap). Boxed areas highlight expression of select genes in lymphatic and vascular endothelial, and ER^Neg^ luminal epithelial cell types.

**Figure 12—figure supplement 2.**
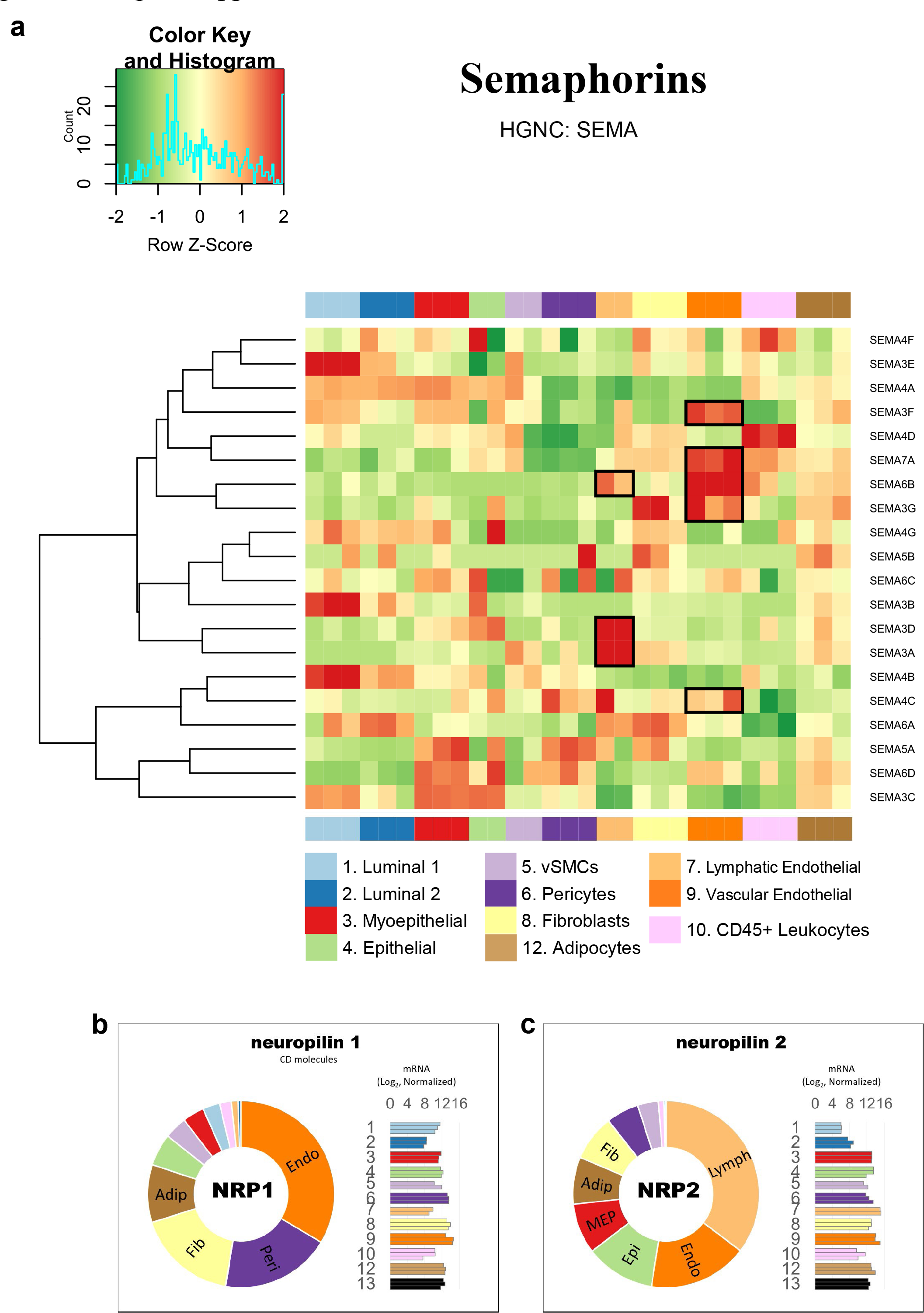
Transcript levels of genes composing the *Semaphorins* gene family. (a) Transcript levels of genes in Semaphorin gene family. VST gene levels are relative (normalized within each row of the heatmap). Boxed areas highlight select genes highly expressed by the lymphatic and vascular endothelial cell types. Normalized mRNA values of **(b)** NRP1 and **(c)** NRP2 (rlog, DEseq2) are provided on log2 scale (bar graph of each biological replicate) and linear scale (donut graph of median value), which are both color-coded by cell type.

**Figure 15—table 1.**
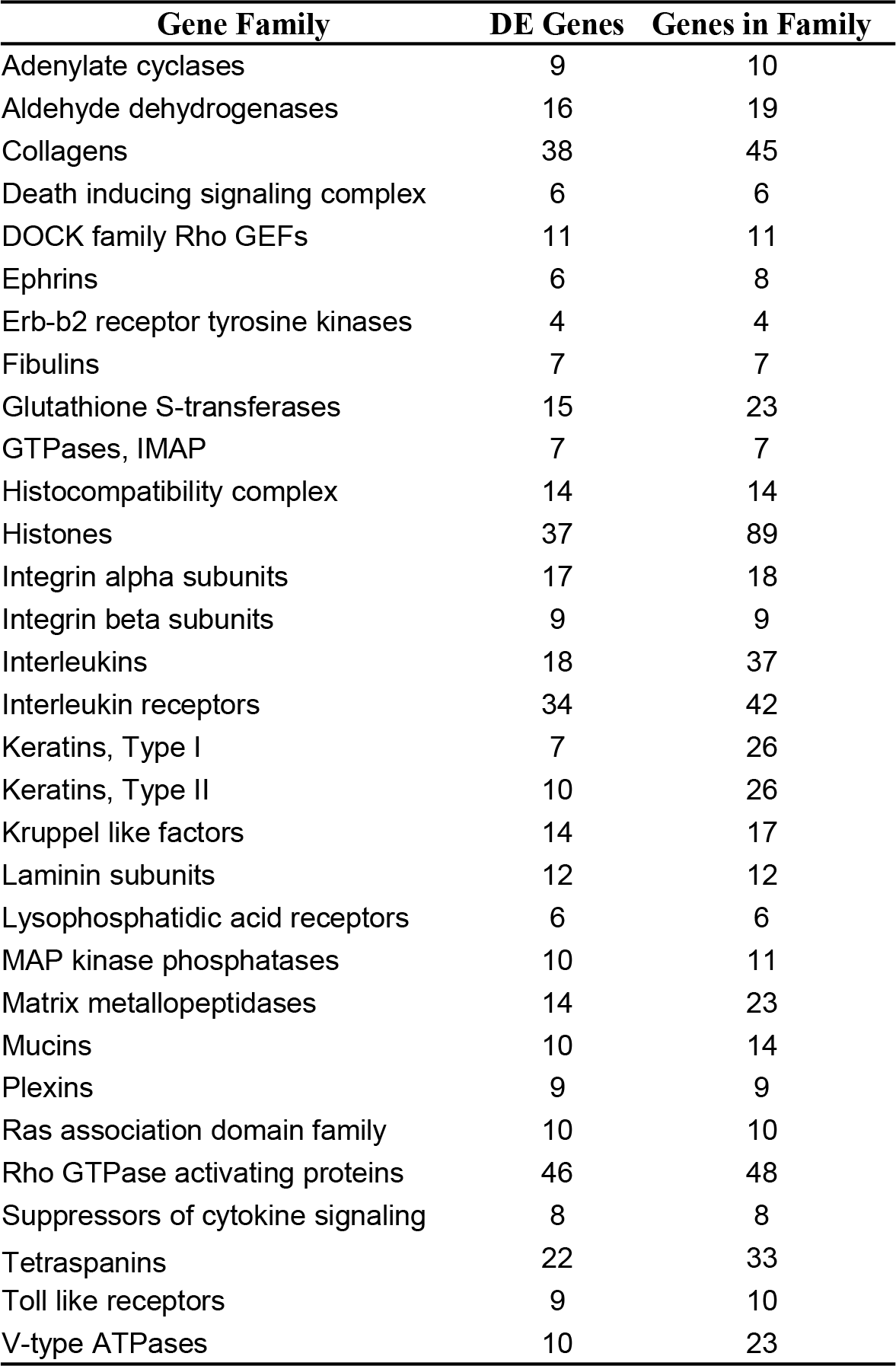
Variably expressed gene types. Genes composing each gene family (HGNC) were assessed by DESeq2 (Adj.*p*-value ≤ 1, across all cell types). The number of genes in each family, along with the numer of those found to be differentially expressed are provided (for gene families ≥ 6 genes).

**Fig.15—figure supplement 1.**
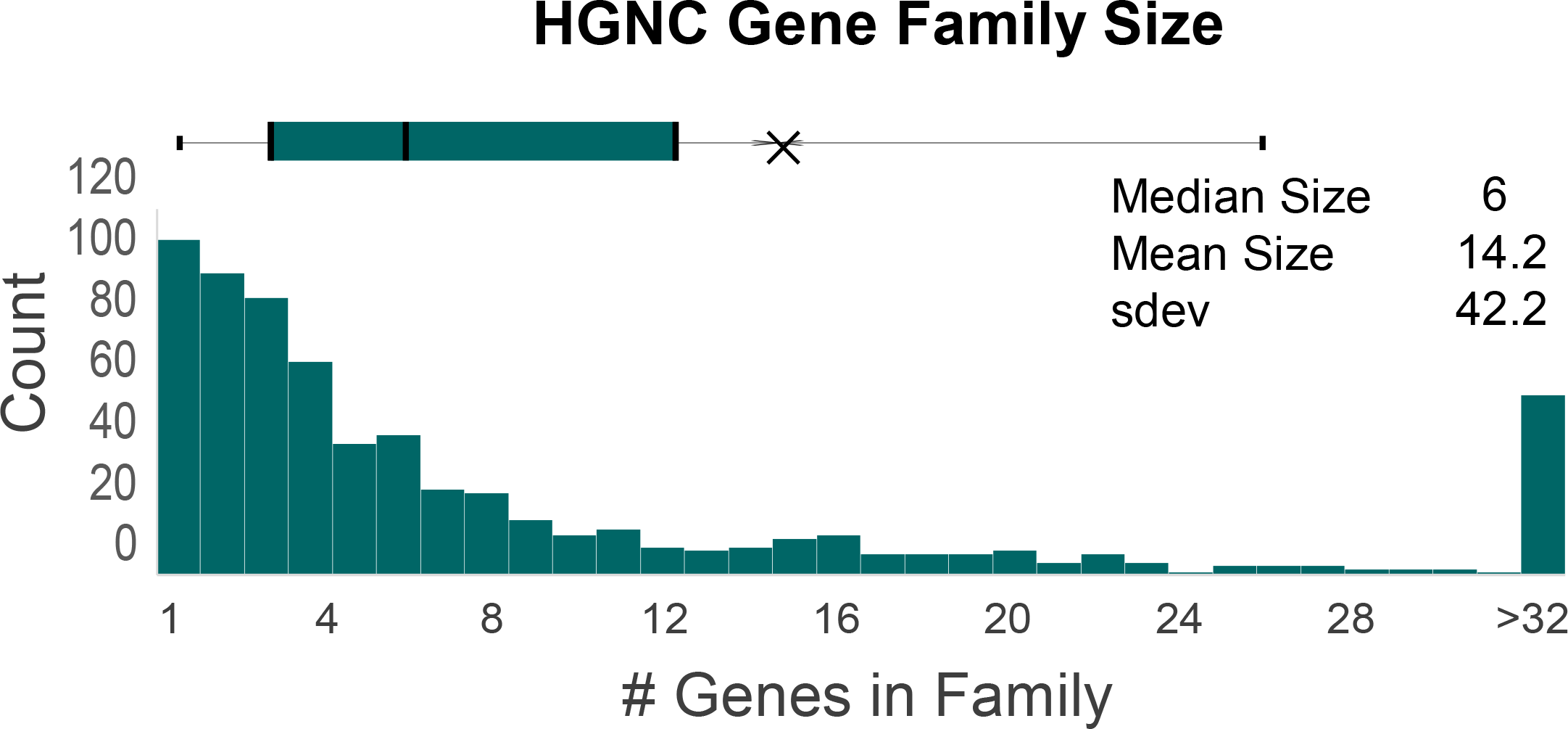
Distribution of gene family size. The sizes of the HGNC families range from 1 to 831 genes. The distribution of these family sizes is provided, along with a box & whisker plot (top). The median family size is six genes per family.

**Fig.15—figure supplement 2.**
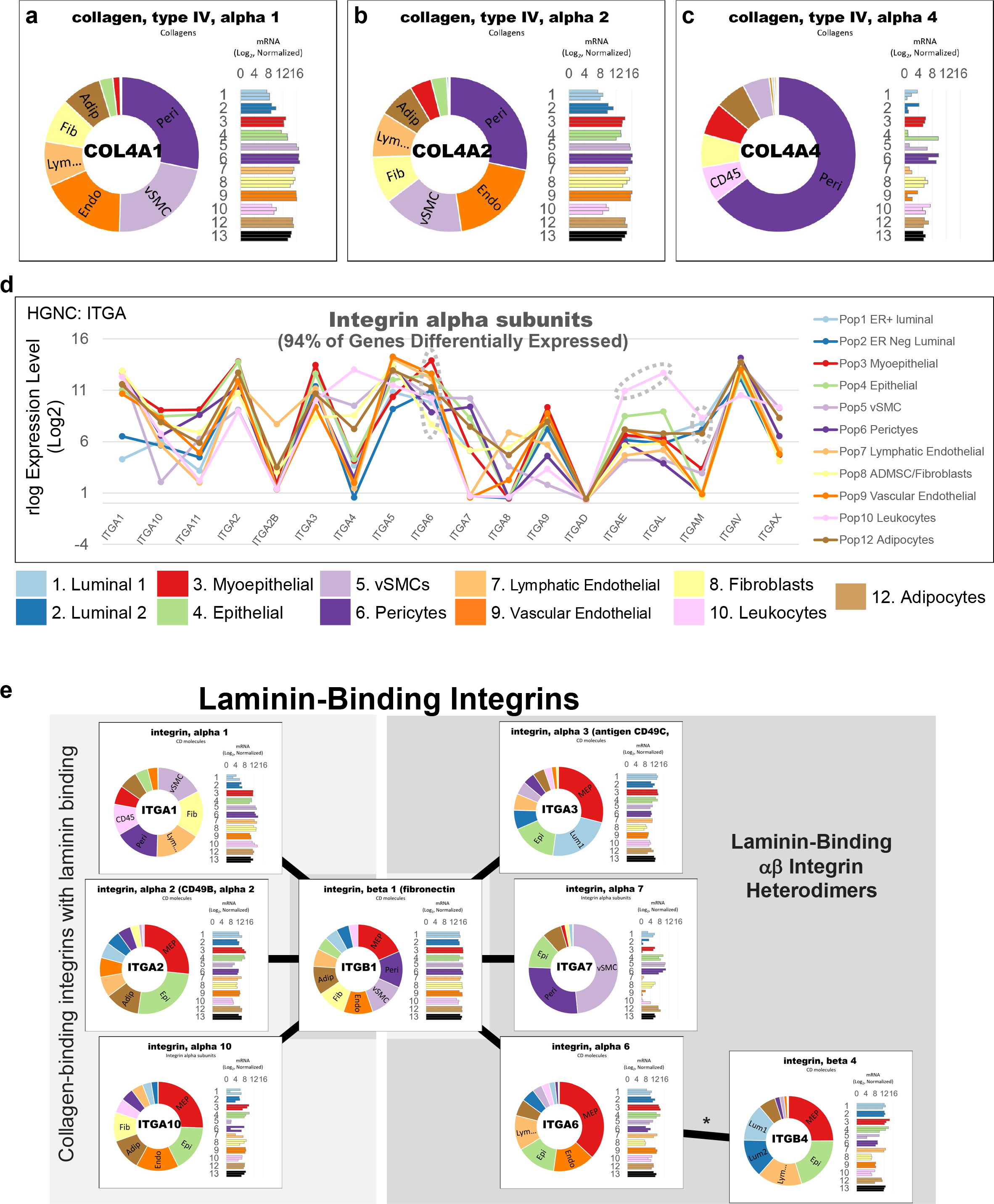
Transcript levels of DE genes. Transcript levels for select Type IV collagen chains: **(a)** COL4A1, **(b)** COL4A2, and **(c)** COL4A4. **(d)** Median mRNA expression (rlog, log2 scale) for integrin alpha subunits. The dashed circles identify notable differentially expressed genes, ITGA6, ITGAE, and ITGAM. **(e)** Transcript levels of the laminin binding integrins. The predominant laminin-binding integrins are αβ heterodimers of: α3β1, α7β1, α7β1 and α6β4. These are integrins are encoded by ITGB1, ITGA3, ITGA7, ITGA6, and ITGB4. Collagen-binding integrins that also show laminin binding properties include heterodimers of: α1β1, α2β1, and α10β1 (encoded by ITGA1, ITGA2, ITGA10, and ITGB1). ITGA7 is predominantly limited to the perivascular cell types, whereas ITGA3, ITGA6, and ITGB4 are largely expressed by the epithelial cell types.

**Figure 16—figure supplement 1.**
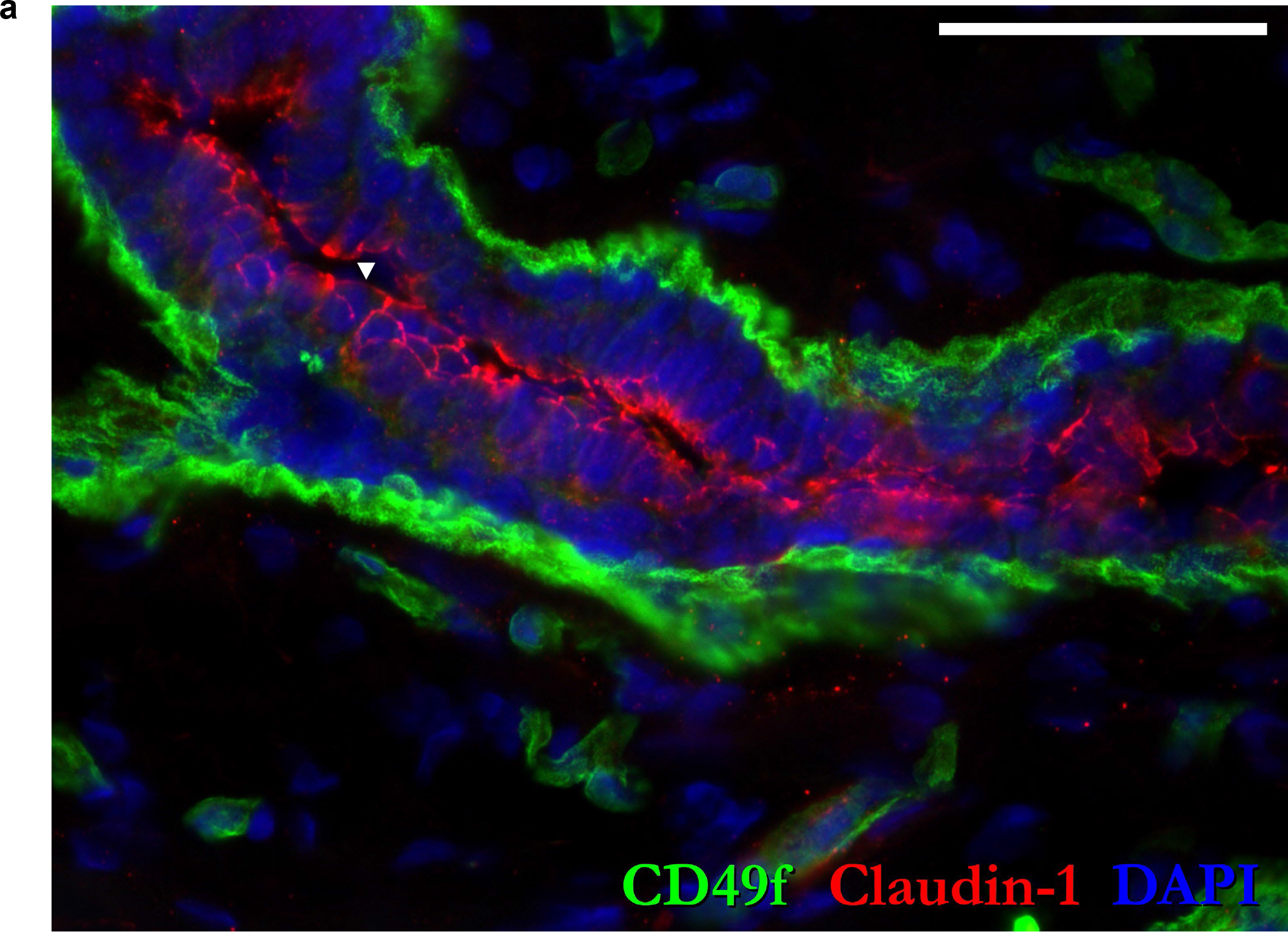
IHC staining. (a) Claudin-1 staining of normal breast tissue (reduction mammoplasty of 28-year-old female). (▾) Claudin-1 staining is found confined to the apical surface of the paracellular space of luminal epithelial cells, circumscribing the luminal surface of the cells. Scale = 50 μm.

**Figure 23—figure supplement 1.**
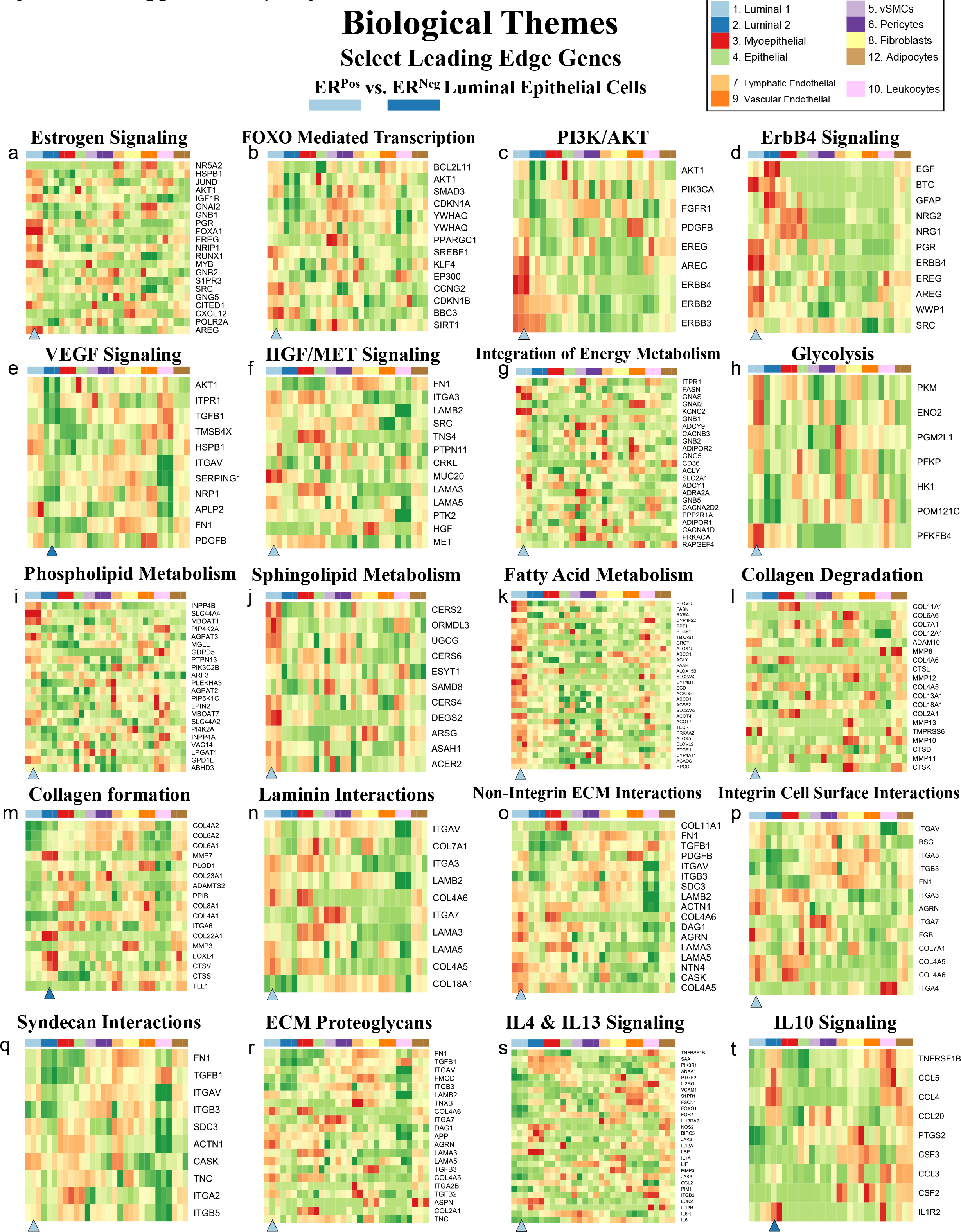
Transcript levels from enriched pathways. (a-t) Transcript levels of leading-edge genes from select pathways enriched in ER^Pos^ or ER^Neg^ luminal cells (from 325 enriched pathways with a FDR of ≤ 10%, GSEA). VST gene levels are relative (normalized by row). Arrowheads under the heatmaps identify the cell type either over- or under-expressing the pathway’s genes. Samples are color-coded. Due to space constraints and the large number of presented genes, their names (rows of the heatmap) must be inspected digitally (magnified in the PDF).

**Figure 23—figure supplement 2.**
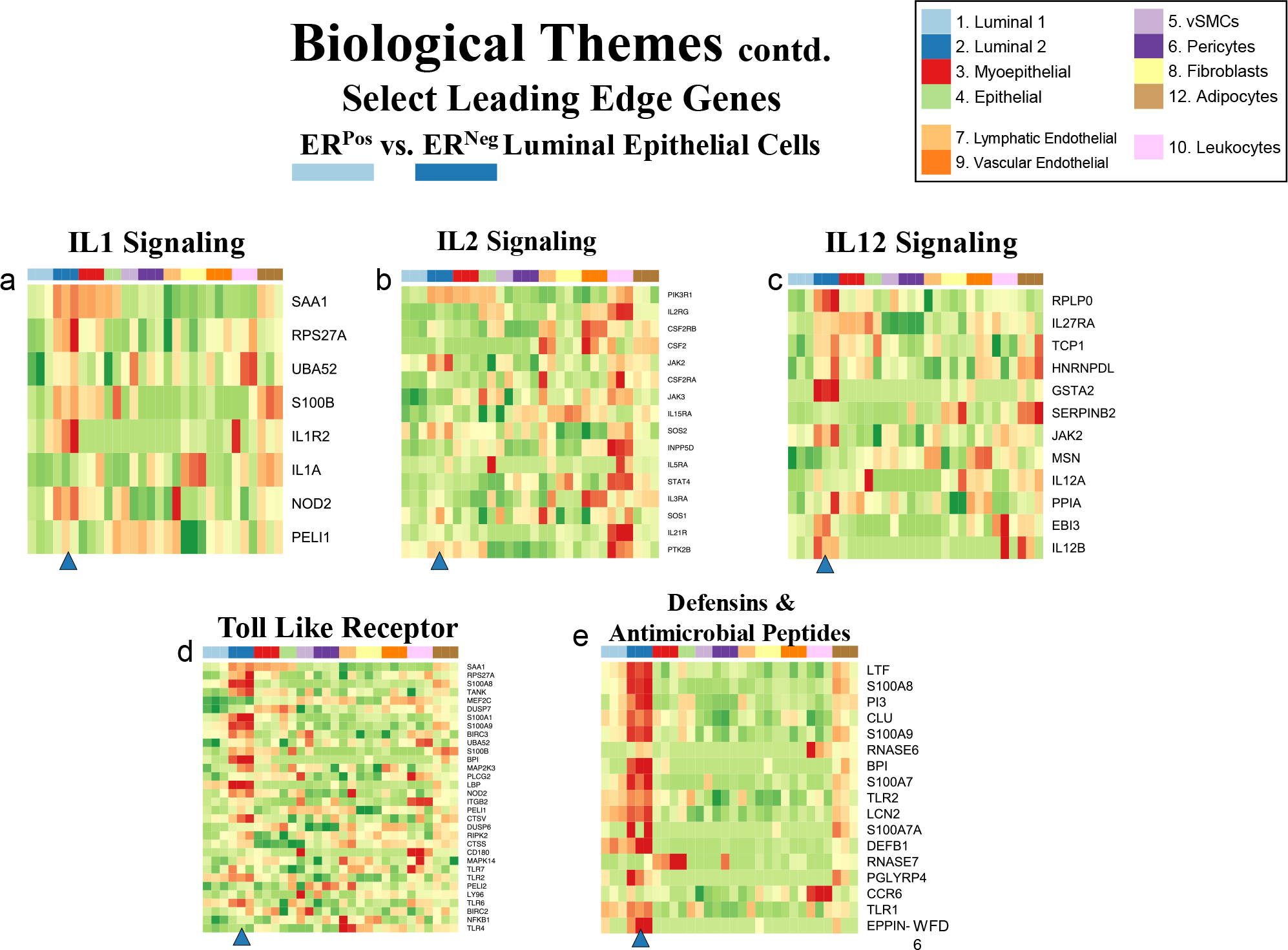
Transcript levels from enriched pathways. (a-e) Transcript levels of leading-edge genes from select pathways enriched in ER^Pos^ or ER^Neg^ luminal cells (from 325 enriched pathways with a FDR of ≤ 10%, GSEA). VST gene levels are relative (normalized by row). Arrowheads under the heatmaps identify the cell type either over- or under-expressing the pathway’s genes. Samples are color-coded. Due to space constraints and the large number of presented genes, their names (rows of the heatmap) must be inspected digitally (magnified in the PDF).

**Figure 23—figure supplement 3.**
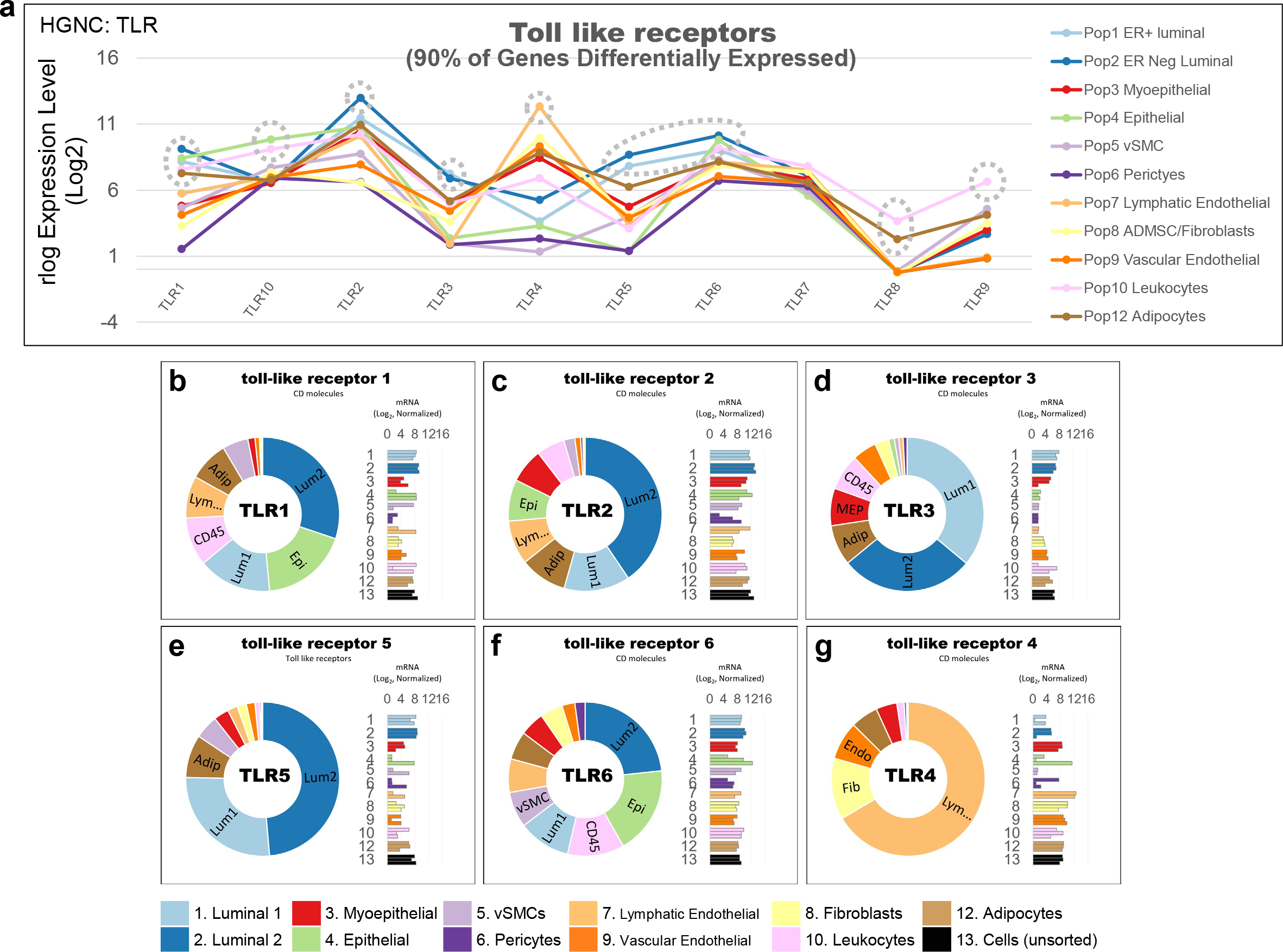
Transcript levels of toll-like receptors. (a) Transcript levels of TLR gene family. **(b-f)** Normalized mRNA values (rlog, DEseq2) provided on log2 scale (bar graph of each biological replicate) and linear scale (donut graph of median value) which are both color-coded by cell type.

**Figure 23—figure supplement 4.**
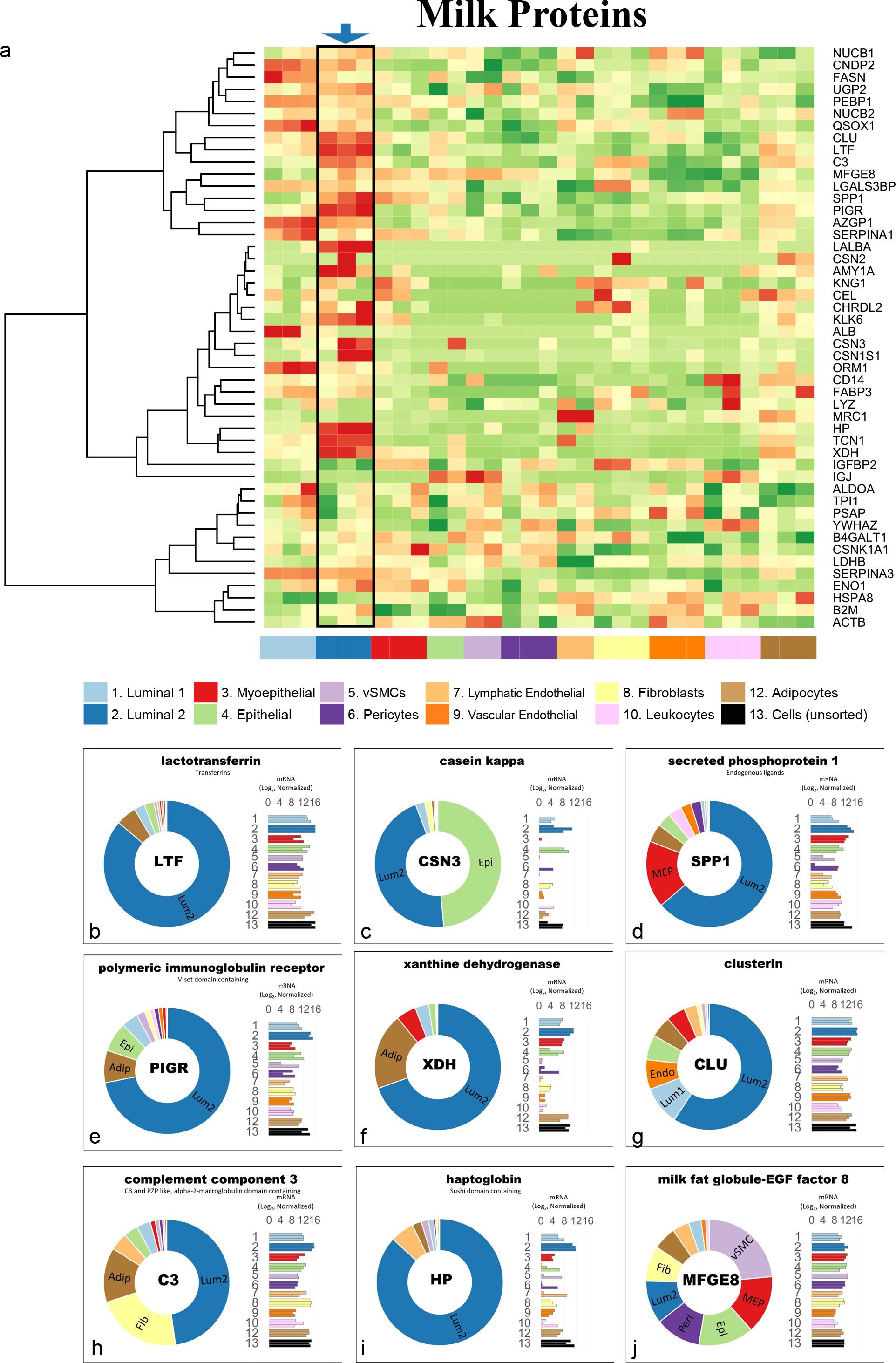
Transcript levels of milk proteins. (a) Transcript levels of milk proteins. VST gene values are relative (normalized by row). **(b-j)** Normalized mRNA values (rlog, DEseq2) provided on log2 scale (bar graph of each biological replicate) and linear scale (donut graph of median value), which are both color-coded by cell type.

**Figure 25—figure supplement 1.**
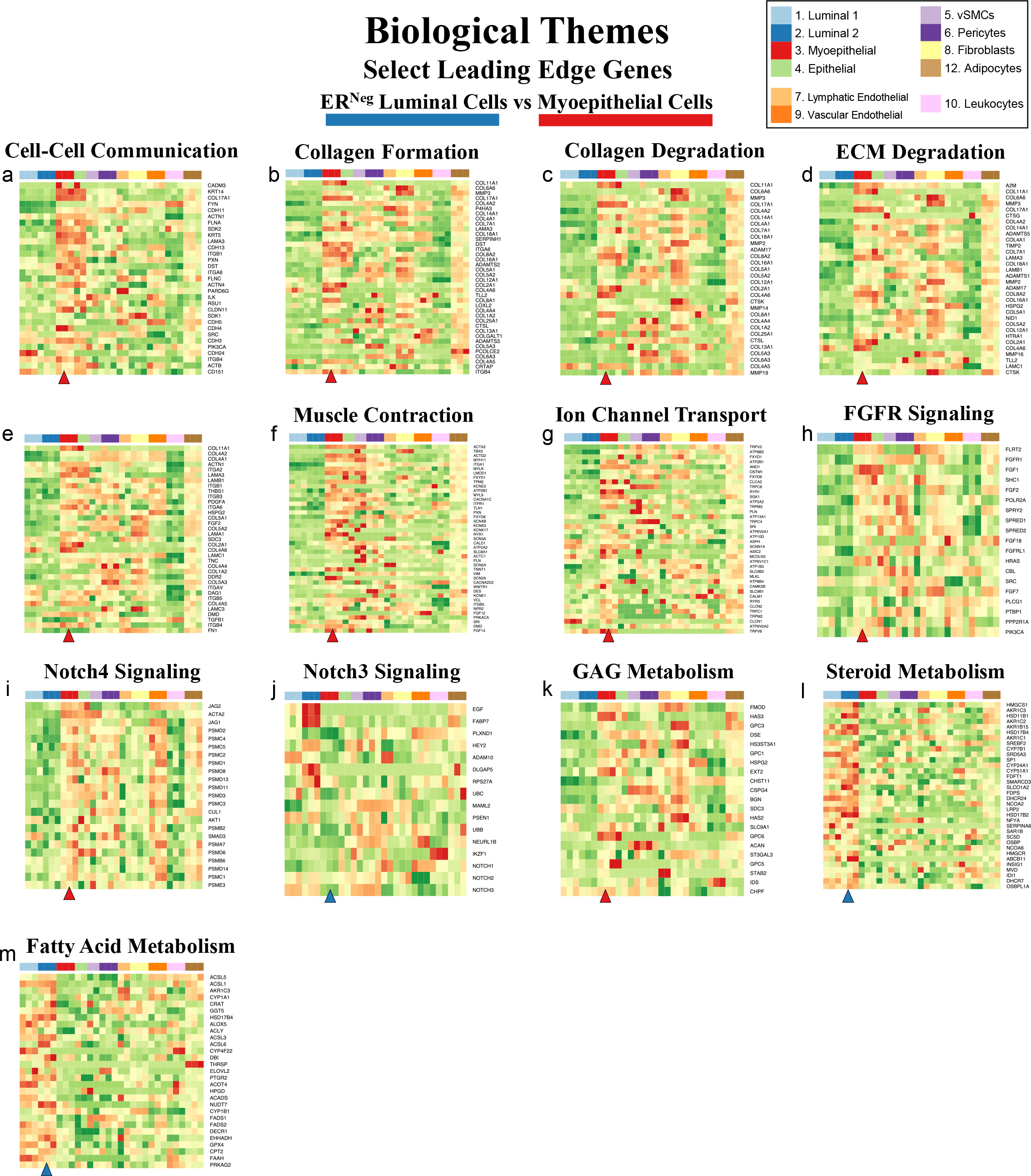
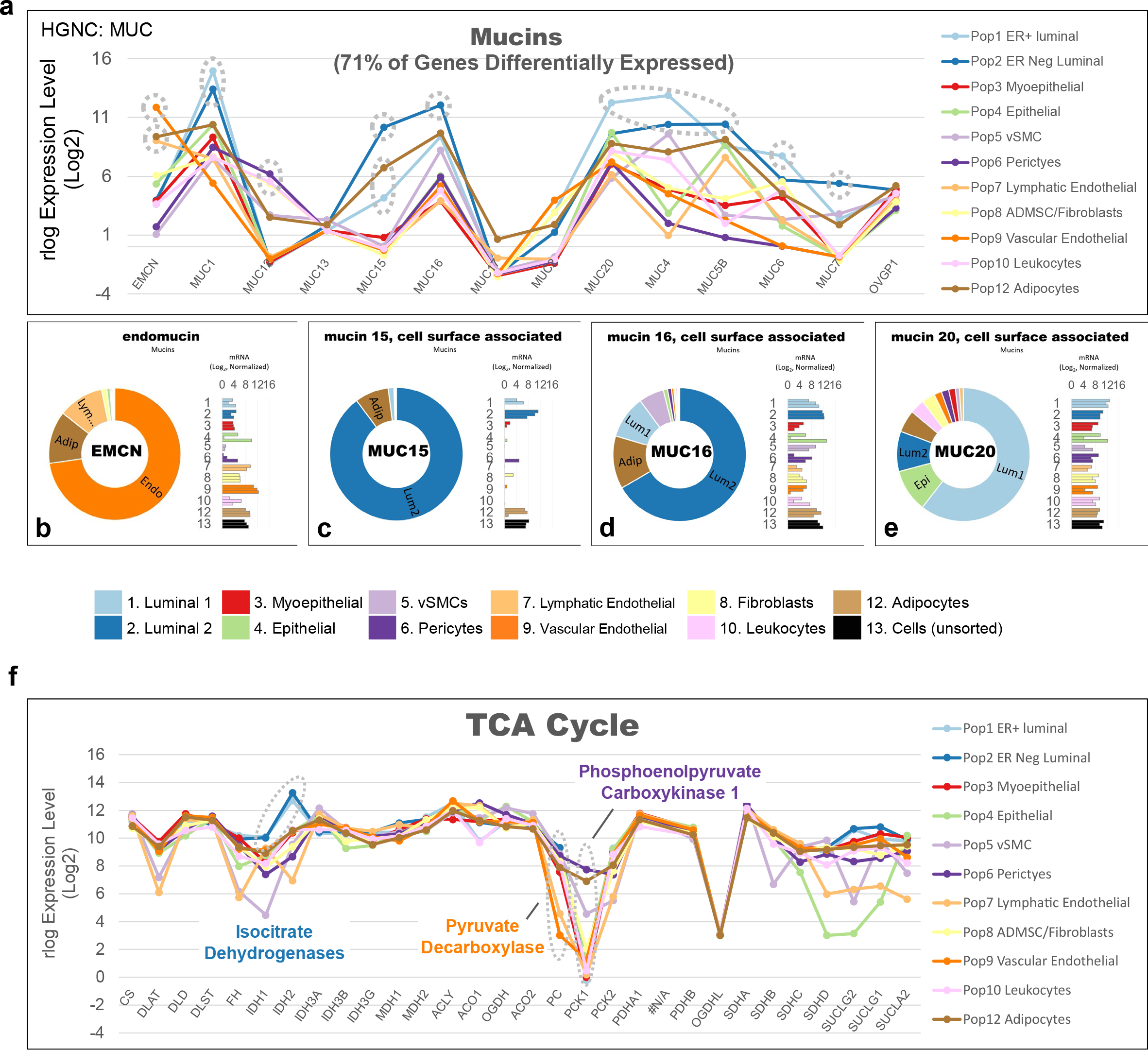
Transcript levels from enriched pathways. Transcript levels of leading-edge genes from select pathways enriched in ER^Pos^ or myoepithelial cells (from 426 enriched pathways with a FDR of ≤ 10%, GSEA). VST gene levels are relative (normalized by row). Arrowheads under the heatmaps identify the cell type, either over- or under-expressing the pathway’s genes. Samples are color-coded. Due to space constraints and the large number of presented genes, their names (rows of the heatmap) must be inspected digitally (magnified in the PDF).

**Figure 27—figure supplement 1.**
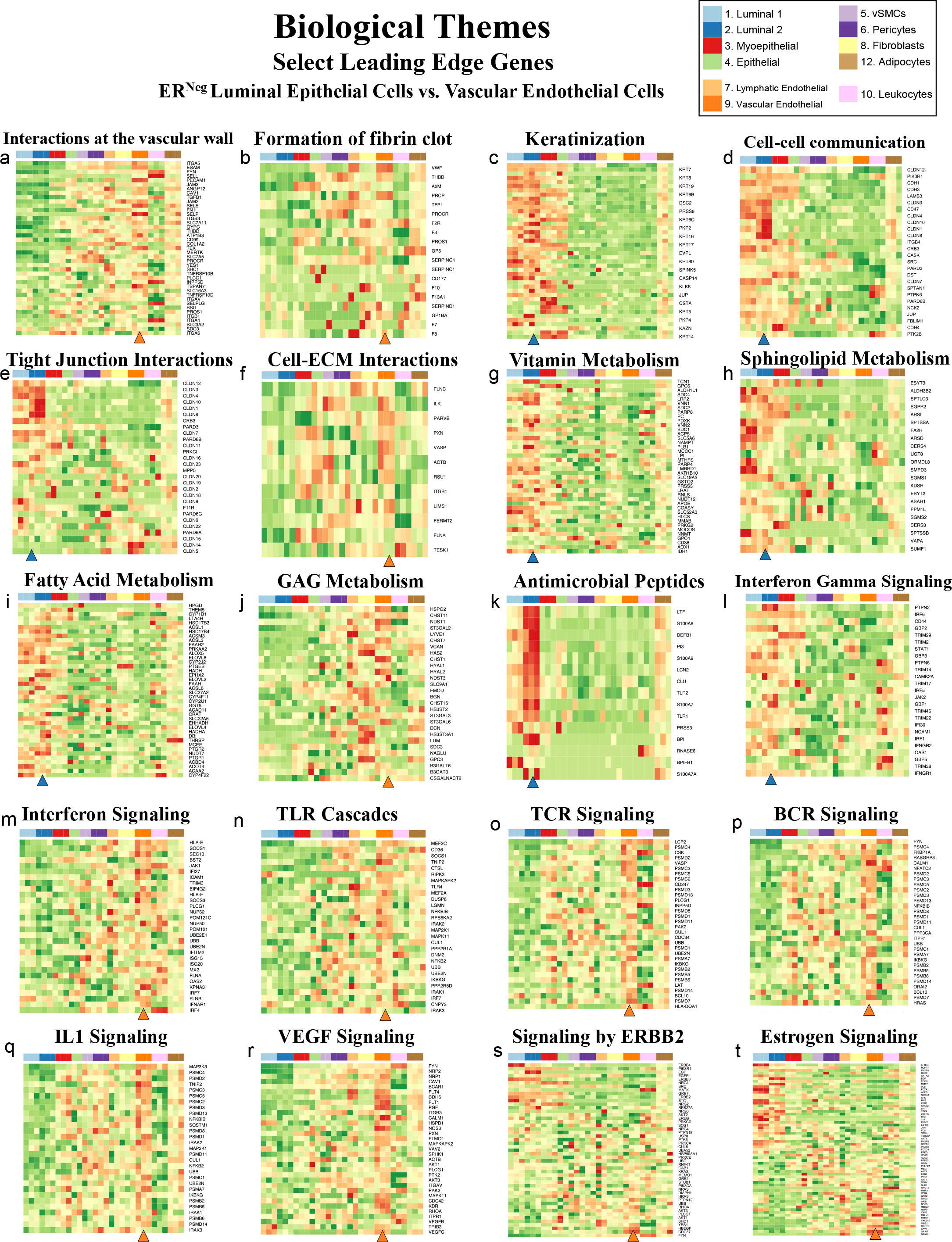
Transcript levels from enriched pathways. Transcript levels of leading-edge genes from select pathways enriched in ER^Pos^ luminal and vascular endothelial cells (from 500 enriched pathways with a FDR of ≤ 10%, GSEA). VST gene levels are relative (normalized by row). Arrowheads under the heatmaps identify the cell type, either over- or under-expressing the pathway’s genes.

Figure 29—figure supplement 1. Transcript levels from enriched pathways. Transcript levels of leading-edge genes from select pathways enriched in lymphatic and vascular endothelial cell (from 240 enriched pathways with a FDR of ≤ 10%, GSEA). VST gene levels are relative (normalized by row). Arrowheads under the heatmaps identify the cell type, either over- or under-expressing the pathway’s genes.

Figure 31—figure supplement 1. Transcript levels from enriched pathways. Transcript levels of leading-edge genes from select pathways enriched in fibroblasts and adipocytes (from 387 enriched pathways with a FDR of ≤ 10%, GSEA). VST gene levels are relative (normalized by row). Arrowheads under the heatmaps identify the cell type, either over- or under-expressing the pathway’s genes.

**Figure 33—figure supplement 1.**
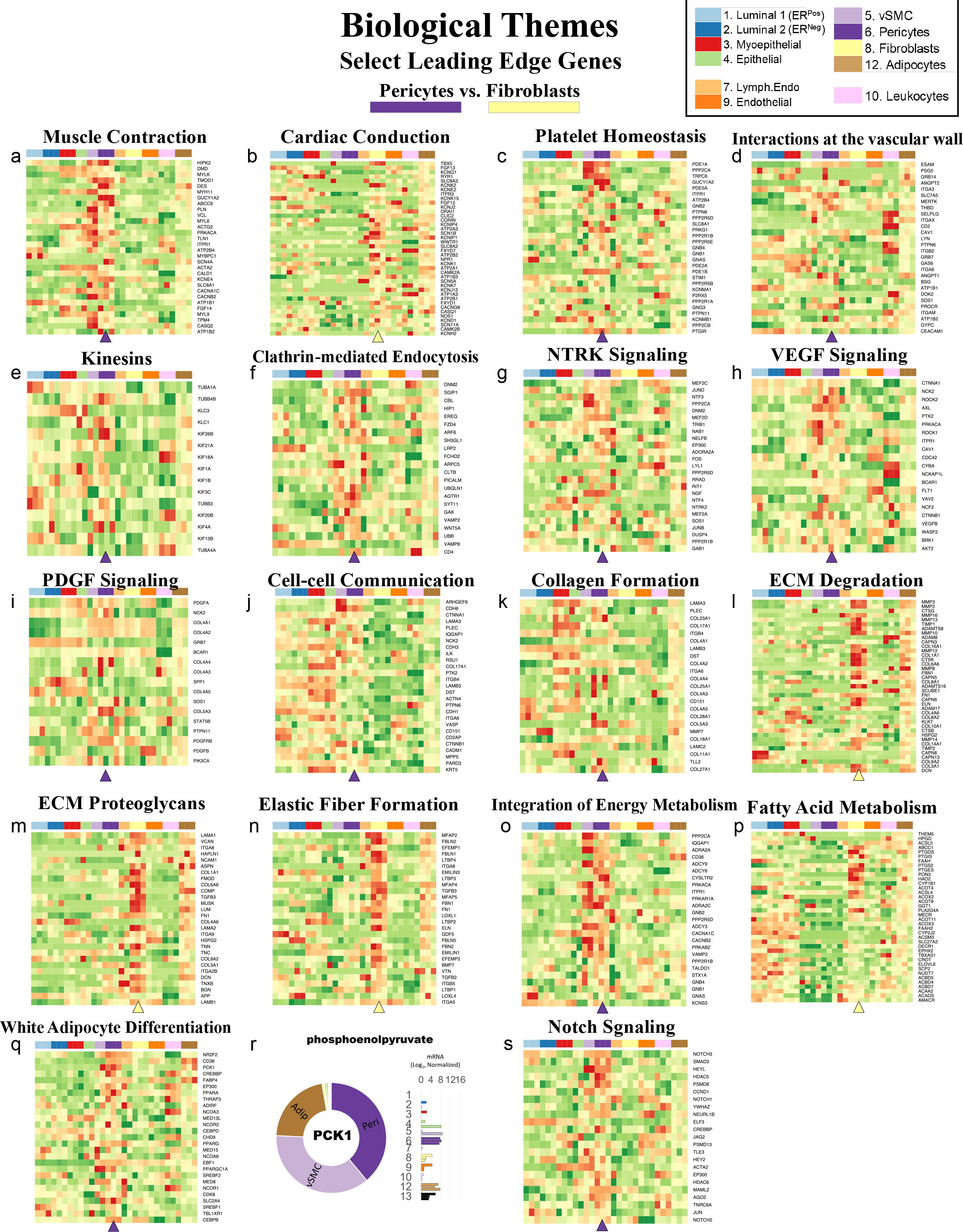
Transcript levels from enriched pathways. Transcript levels of leading-edge genes from select pathways enriched in pericytes and fibroblasts (from 477 enriched pathways with a FDR of ≤ 10%, GSEA). VST gene levels are relative (normalized by row). Arrowheads under the heatmaps identify the cell type, either over- or under-expressing the pathway’s genes.

